# Simplifying bioinformatics data analysis through conversation

**DOI:** 10.1101/2023.10.29.564479

**Authors:** Zhengyuan Dong, Han Zhou, Yifan Jiang, Victor Zhong, Yang Young Lu

## Abstract

The rapid advancements in high-throughput sequencing technologies have produced a wealth of omics data, facilitating significant biological insights but presenting immense computational challenges. Traditional bioinformatics tools require substantial programming expertise, limiting accessibility for experimental researchers. Despite efforts to develop user-friendly platforms, the complexity of these tools continues to hinder efficient biological data analysis. In this paper, we introduce BioMANIA– an AI-driven, natural language-oriented bioinformatics pipeline that addresses these challenges by enabling the automatic and codeless execution of biological analyses. BioMANIA leverages large language models (LLMs) to interpret user instructions and execute sophisticated bioinformatics workflows, integrating API knowledge from existing Python tools. By streamlining the analysis process, BioMANIA simplifies complex omics data exploration and accelerates bioinformatics research. Compared to relying on general-purpose LLMs to conduct analysis from scratch, BioMANIA, informed by domain-specific biological tools, helps mitigate hallucinations and significantly reduces the likelihood of confusion and errors. Through comprehensive benchmarking and application to diverse biological data, ranging from single-cell omics to electronic health records, we demonstrate BioMANIA’s ability to lower technical barriers, enabling more accurate and comprehensive biological discoveries.

## 1 Introduction

Recent advancements in high-throughput sequencing technologies have enabled researchers to collect vast amounts of omics data—including genomics, transcriptomics, epigenomics, proteomics, metabolomics, and microbiomics—across various tissues, organs, individuals, and species. Large-scale consortium projects, such as The Cancer Genome Atlas (TCGA) (Weinstein et al, 2013) and the Encyclopedia of DNA Elements (ENCODE) (ENCODE Project Consortium, 2012), have made genomic, transcriptomic, and epigenomic data from diverse samples openly accessible. While this wealth of data has sparked interest in understanding molecular mechanisms and exploring therapeutic applications, it also poses significant challenges in extracting meaningful insights from complex data analyses.

Over the years, significant efforts have been made to tackle these data analysis challenges. These efforts have produced valuable insights from two key perspectives (Zhou et al, 2023): (1) computational researchers develop innovative methods and tools for biological data analysis, and (2) experimental researchers apply existing tools and pipelines to analyze and interpret the data. Initially, bioinformatics tools were developed within specialized programming frameworks, including BioPerl (Stajich et al, 2002), BioRuby (Goto et al, 2010), Bioconductor (Gentleman et al, 2004), Biopython (Cock et al, 2009), and BioJava (Holland et al, 2008). However, utilizing these tools requires considerable bioinformatics programming expertise, which many experimental researchers may not possess. To lower the programming barrier, more advanced tools were later developed, allowing users to interactively explore and visualize omics data without requiring extensive programming skills. Representative tools include Galaxy (Giardine et al, 2005)–a platform offering pre-assembled bioinformatics pipelines for interactive large-scale genome analysis and the cBio Cancer Genomics Portal (Cerami et al, 2012), which enables interactive exploration and visualization of preloaded multidimensional cancer genomics datasets, among others. These tools have predefined analyses and often require users to navigate complex interfaces and procedures. Additionally, with advancements in algorithms, bioinformatics pipelines evolve rapidly to integrate cutting-edge solutions.

Despite the intention to assist experimental researchers, the advancement of bioinformatics tools presents a learning barrier, requiring substantial professional skills and domain-specific expertise. For example, in single-cell RNA sequencing (scRNA-seq) data analysis, there are over 1,000 tools available (Zappia and Theis, 2021), offering diverse computational pipelines, software libraries, and analytical workflows to examine the data from various perspectives. Furthermore, the complexity of scRNA-seq data analysis becomes evident through its multi-step pipeline, which requires substantial specialized knowledge and expertise. The analysis involves numerous considerations (He et al, 2022), including quality control, feature selection, batch correction, dimensionality reduction, cell clustering, cell annotation, trajectory inference, marker gene detection, and more. Completing these steps requires researchers to have both advanced programming skills and a strong biological background, which increases the cost and complexity of conducting single-cell analysis. Thus, there is an urgent need to bridge the gap between researchers’ intentions and their understanding of existing bioinformatics tools, facilitating the automatic execution of these tools to produce analysis results for biological data.

From a practitioner’s perspective, an ideal bioinformatics framework should lower technical barriers in data analysis for biological scientists by incorporating the following features: (1) standardized and automated pipelines that enable shareable and reproducible analysis (Wratten et al, 2021), and (2) user-friendly, codeless interfaces that help overcome the learning barrier for researchers with limited programming skills (Li et al, 2021). Coincidentally, large language models (LLMs) such as GPT-3 (Brown et al, 2020) and GPT-4 (Bubeck et al, 2023) have demonstrated exceptional capabilities in natural language processing (NLP) tasks, including human-like conversation (Touvron et al, 2023), mathematical reasoning (Wei et al, 2022), code generation (Chen et al, 2021), and more. With the advancement of LLMs, they have increasingly shown potential in utilizing external tools to solve complex problems across various application domains (Qu et al, 2024), much like humans do. For example, in the biological domain, LLMs have proven valuable in bioinformatics education (Shue et al, 2023; Tu et al, 2024) and in assisting with programming tasks (Piccolo et al, 2023; Zhou et al, 2023). Building on these advancements, a natural question arises: “How can we enable effective and user-friendly bioinformatics data analysis using LLMs?”

Although leveraging LLMs to assist in bioinformatics data analysis is a promising prospect, directly applying current state-of-the-art general-purpose LLMs to specific biological data analysis tasks presents significant challenges. Despite their capabilities in NLP, these general-purpose LLMs lack accurate, up-to-date knowledge of domain-specific biological tools and concepts, often resulting in outputs that lack meaningful biological significance (Qu et al, 2024). A major limitation of general-purpose LLMs is their tendency to “hallucinate”, generating confident yet inaccurate responses when faced with specialized biological queries (Refer to Sec. A.4 for failure cases). These hallucinated responses can cause confusion and misdirection, making it harder for researchers to achieve their objectives. These limitations highlight the need for tool-specific LLMs that integrate deep, accurate, tool-specific knowledge to enable efficient, professional, and automated execution of biological data analysis tasks.

In this paper, surmount the bioinformatics data analysis barrier using natural languages and propose an artificial intelligence (AI)-driven, natural language-oriented bioinformatics data analysis pipeline, named BioMANIA (BioinforMatics dAta aNalysIs through conversAtion), as illustrated in Fig. 1. BioMANIA is designed to directly understand natural language instructions from users and efficiently execute complex biological data analysis tasks. Importantly, considering the limitations of LLMs in lack of domain-specific biological knowledge, BioMANIA explicitly learns about API information and their interactions from the source code (*e*.*g*., hosted in GitHub) and documentation (*e*.*g*., hosted in ReadTheDocs) of any off-the-shelf well-documented, open-source Python tools. BioMANIA’s innovation enables automated, codeless biological data analysis by simplifying the complex process into manageable steps: interpreting the user’s intent, breaking it down into specific tasks if needed, mapping tasks to API calls, executing the APIs with inferred arguments, and troubleshooting any coding errors. For the purpose of reproducibility, BioMANIA summarizes the execution results in a user-friendly format, organizes the API calls into a downloadable script, and generates a comprehensive report documenting the entire data analysis process. As a result, with minimum human intervention, BioMANIA enables the end-to-end automatic and codeless execution of any off-the-shelf biological analysis workflow.

**Fig. 1:**
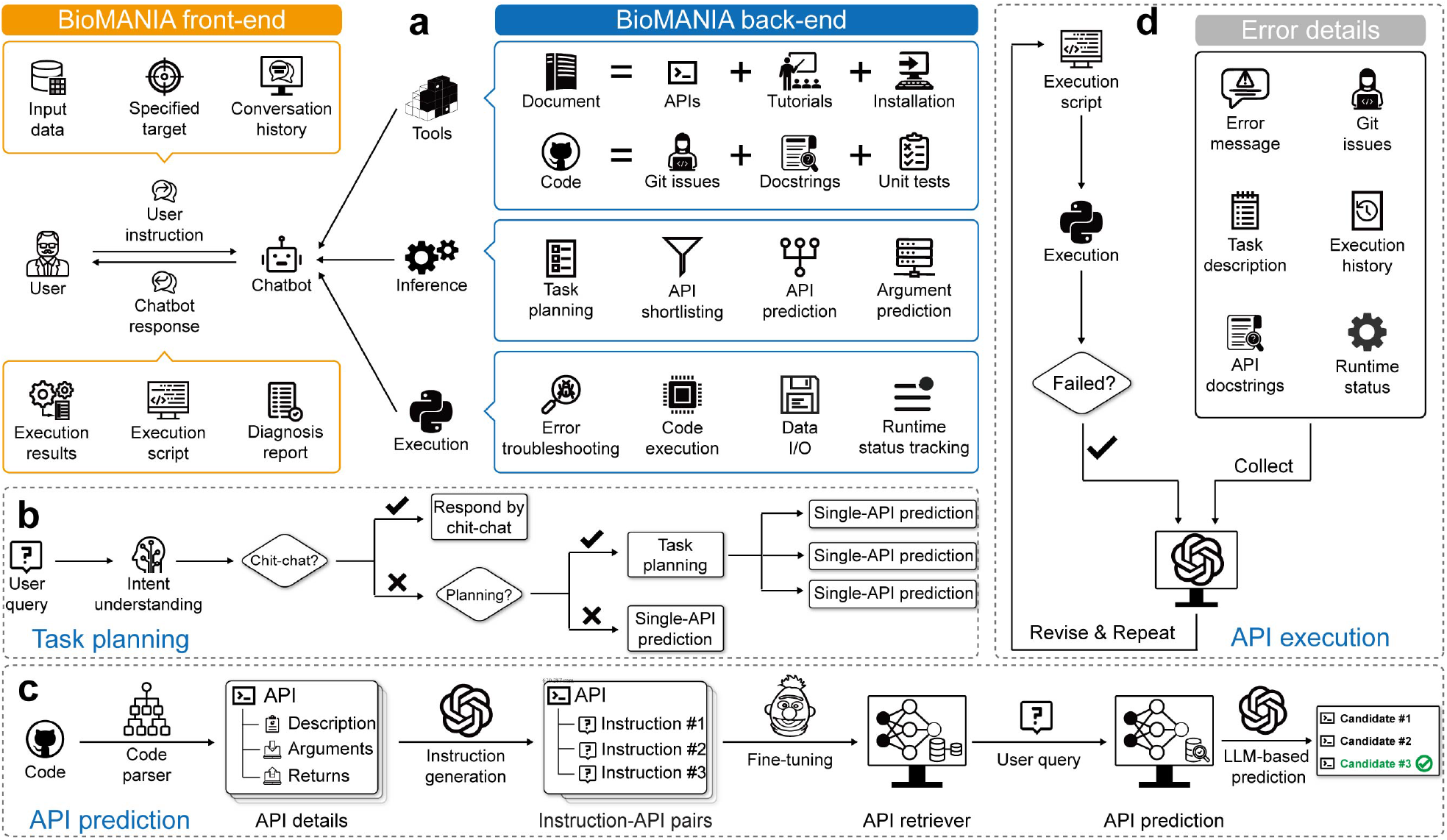
Overview of BioMANIA. (**a**) BioMANIA is composed of two main components: the front-end and the back-end. The BioMANIA front-end receives the user’s data and instructions, then sends this information, along with the chat history, to the BioMANIA back-end for intent understanding, task planning, and execution. It then collects the outcomes from the back-end, which include execution results with code and a detailed report. The BioMANIA back-end first learns about API information and their interactions from the source code and documentation of a well-documented, open-source Python tool. The back-end also processes the information received from the front-end, interprets the user’s intent, breaks it down into specific tasks, maps these tasks to API calls, executes the APIs with the inferred arguments, and sends the execution results back to the front-end. (**b**) Given the user’s instruction, BioMANIA determines their intent in two steps. First, it identifies whether the user is asking an API-specific data analysis question or engaging in chit-chat. For API-specific questions, it further determines whether the task requires multiple API calls. If multiple API calls are required, BioMANIA breaks the user’s instruction into several tasks and addresses each one individually. (**c**) BioMANIA analyzes the source code to extract potential APIs, prompts LLMs to understand their functionalities, and generates synthetic instructions corresponding to API calls. These instruction-API pairs are used to train an API retriever, which narrows down candidate APIs to the top-k for a given instruction, and an API predictor, which selects the exact API from the retrieved subset. (**d**) BioMANIA uses a Python interpreter to execute the predicted API calls. In case of failure, BioMANIA gathers all relevant error information and prompts LLMs to troubleshoot and iteratively revise the API call code.

We have conducted comprehensive evaluation to demonstrate BioMANIA’s empirical utility. We first rigorously benchmarked the performance of each individual step of BioMANIA, using both LLM-generated and human-annotated instructions for hundreds of APIs. Next, we applied BioMANIA to a range of commonly used bioinformatics tools, covering diverse application domains–from single-cell RNA-seq analysis (Wolf et al, 2018), single-cell spatial omics analysis (Palla et al, 2022), single-cell epigenomics analysis (Zhang et al, 2024), to electronic health record analysis (Heumos et al, 2024). Compared to using LLMs to conduct analysis from scratch, which can lead to hallucinated responses and potential misdirection, the structured steps in BioMANIA, informed by domain-specific biological tools, help mitigate hallucinations, significantly reducing the likelihood of confusion and errors. For example, BioMANIA successfully completed 18 out of 21 single-cell transcriptome data analysis tasks, with minimal troubleshooting required for most tasks. In contrast, general-purpose LLMs, when used to perform the analysis from scratch, succeeded in only 5 out of 21 tasks. In summary, BioMANIA simplifies the exploration of omics data and accelerates bioinformatics research and enables more accurate and comprehensive biological discoveries. Also, BioMANIA significantly lowers the technical barriers to conducting state-of-the-art biological data analysis, with the potential to usher in a new research paradigm.

## 2 Results

### 2.1 The design of Python tools influence the effectiveness of conversational data analysis

BioMANIA is designed to enable conversational, codeless analysis for any off-the-shelf Python tools. However, it’s important to recognize that these tools were not originally designed for conversational analysis. This raises a natural question: “What are the challenges in making existing tools suitable for conversational analysis?” To assess how the design and implementation of existing Python tools impact the effectiveness of conversational data analysis, we gathered a variety of commonly used bioinformatics tools, spanning diverse application domains such as single-cell omics analysis and electronic health records (Refer to Sec. 4.2 for more details). These Python tools used by BioMANIA exhibit varying levels of API ambiguity, validity, tutorial and unit-test coverage, argument distribution, tutorial comprehensiveness, and Git support, reflecting a wide range of complexity (Fig. 2**a**).

**Fig. 2:**
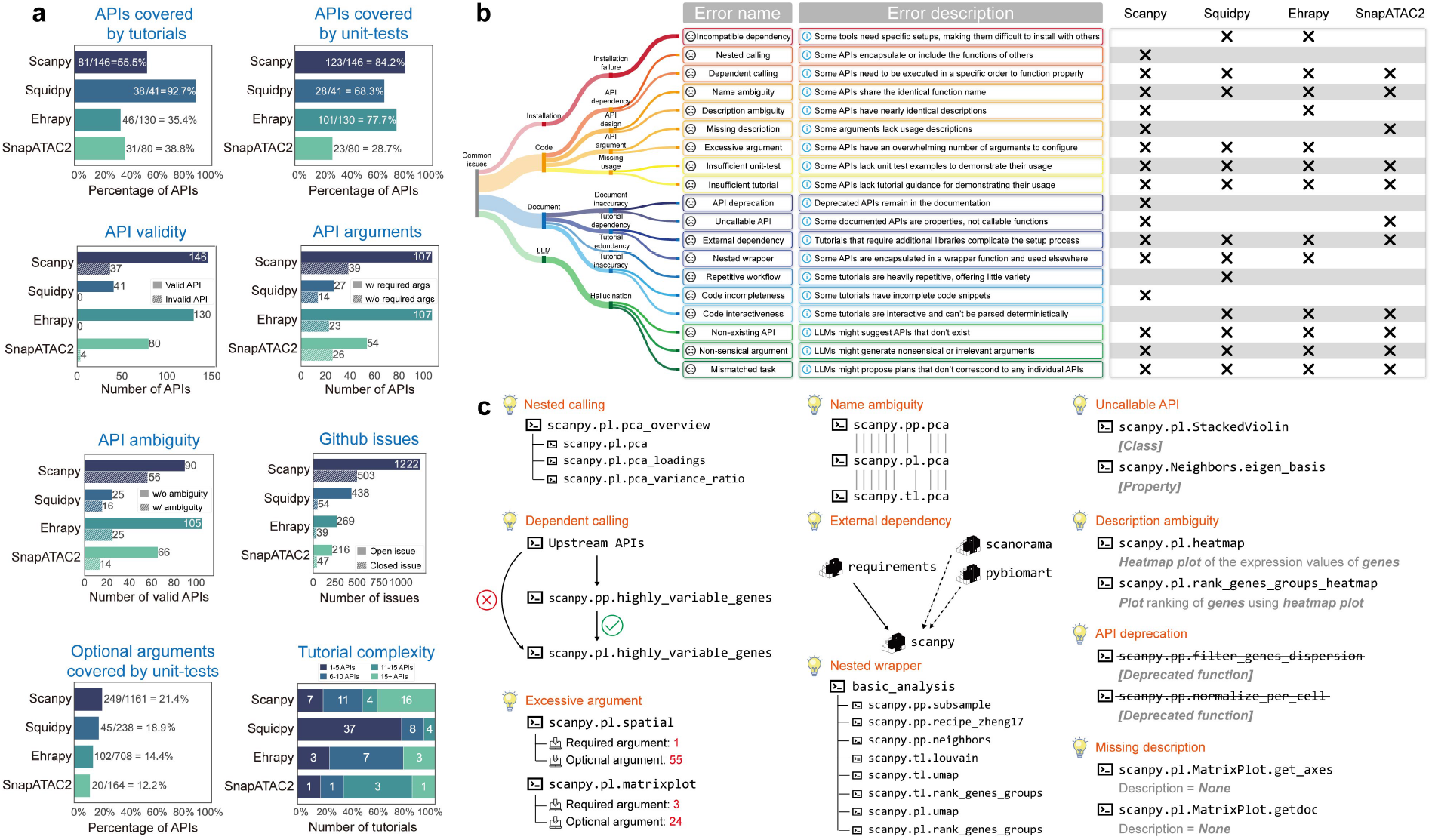
The design and implementation of Python tools influence the effectiveness of conversational data analysis. (**a**) The Python tools used by BioMANIA display varying levels of API ambiguity, validity, tutorial and unit-test coverage, argument distribution, tutorial comprehensiveness, and Git support. (**b**) The common issues encountered by BioMANIA are categorized into four groups, each based on their role in the BioMANIA pipeline. A concise description is provided for each issue, along with the indication of its presence or absence in each Python tool. (**c**) The major issues are illustrated with real examples we encountered in the Python tool Scanpy.

During our analysis of these tools, we identified 19 common issues encountered throughout the BioMANIA pipeline. These issues were categorized into four groups based on their role (Fig. 2**b**). We break down the issue categories and discuss each one in the order it occurs within the BioMANIA analysis pipeline. We observed that some tools could not be installed smoothly by following the documentation instructions. For example, the Python tool Squidpy v1.3.0 is incompatible with versions of the dependency library Napari earlier than v0.4,as confirmed by its Github issue (https://github.com/scverse/squidpy/issues/743).

We observed that many issues stem from the Python tool’s code, which are critical for BioMANIA tool parsing. These code-related issues encompass various aspects of code design and implementation, including API design, argument structure, and usage guidance. In the API design, some APIs either share identical function names or have nearly identical descriptions, making them difficult to distinguish and use correctly. For example, the Python tool Scanpy has three nearly identical APIs related to principal component analysis, *i*.*e*., “scanpy.pp.pca”, “scanpy.pl.pca”, and “scanpy.tl.pca”, differing by only one edit distance. As a result, improper API design can hinder effective API shortlisting and prediction from user instructions (Refer to Sec. 2.2 for more details). In terms of argument design, some tools feature highly complex APIs with up to dozens of arguments, which can overwhelm the argument prediction process and make conversational analysis more laborious. For example, Scanpy has an API called “scanpy.pl.spatial” for scatter plots in spatial coordinates, which has a total of 56 arguments, most of which are optional (Fig. 2**c**). As a result, the excessive number of arguments poses challenges for BioMANIA, as most of them are not demonstrated through unit tests or tutorials. Even Scanpy, which has the highest argument coverage, only manages to cover 21.4% of argument usage (Fig. 2**a**).

Apart from code issues, documentation-related problems further exacerbate the challenges in BioMANIA tool parsing. Although documentation is expected to provide detailed information about API usage, inaccuracies and complications in the documentation can adversely affect its effectiveness. However, we observed that some tools lack up-to-date documentation, listing APIs that have already been deprecated (Fig. 2**c**). This inconsistency between the documentation and the code leads to confusion when parsing the tool for conversational analysis. Furthermore, we consider tutorials to be valuable resources for demonstrating the interplay among APIs, offering guidance on how to coordinate multiple APIs to complete real-world tasks. However, we observed that some tutorials are overly complicated, making it difficult for BioMANIA to effectively capture the interplay among APIs. For example, some tutorials are designed to be interactive (*e*.*g*., The CellProfiler tutorial in Squidpy), requiring user input within loops to run correctly, which disrupts the automated process of BioMANIA. Additionally, some tutorials follow software engineering principles by encapsulating multiple APIs within a wrapper function for repetitive use, making them more difficult to analyze (Fig. 2**c**).

Lastly, BioMANIA leverages LLMs for predicting APIs and arguments (Fig.1), but the use of LLMs inevitably leads to occasional hallucinations, where confident yet inaccurate responses are generated. For example, LLMs may suggest APIs that don’t exist or generate irrelevant, even nonsensical, arguments based on the user’s instruction (Fig. 2**b**). Although hallucinated responses are inevitable when using LLMs, having the detailed descriptions of APIs and arguments can help easily identify these hallucinations, thereby reducing potential confusion and misdirection.

### 2.2 BioMANIA accurately manages each step in its pipeline

BioMANIA relies on several key components to enable automated, codeless biological data analysis. That said, the effectiveness of BioMANIA critically depends on these components working together seamlessly and efficiently. Thus, we break down the BioMANIA pipeline and assess the effectiveness of BioMANIA in each of its key components, which contribute at different stages of the pipeline.

We first investigated BioMANIA’s performance in distinguishing between user queries related to API-encoded instructions and casual chit-chat (Refer to Sec. 4.1.2 for more details). We randomly sampled an equal number of chit-chat samples from two existing chit-chat datasets: Topical-Chat (Gopalakrishnan et al, 2019) and Qnamaker (Agrawal et al, 2020). Qualitatively, the t-SNE projection reveals a clear separation between TF-IDF embeddings of chit-chat samples from two datasets and API-encoded instructions generated by LLMs (Fig. 3**a**). Quantitatively, this clear separation enables a simple nearest centroid classifier to achieve nearly perfect classification accuracy on the test dataset, with a minimum accuracy of 99.19% across different Python tools. The accurate recognition of chit-chat samples allows BioMANIA to better interpret the user’s intent and decide whether to proceed with determining the corresponding API calls.

**Fig. 3:**
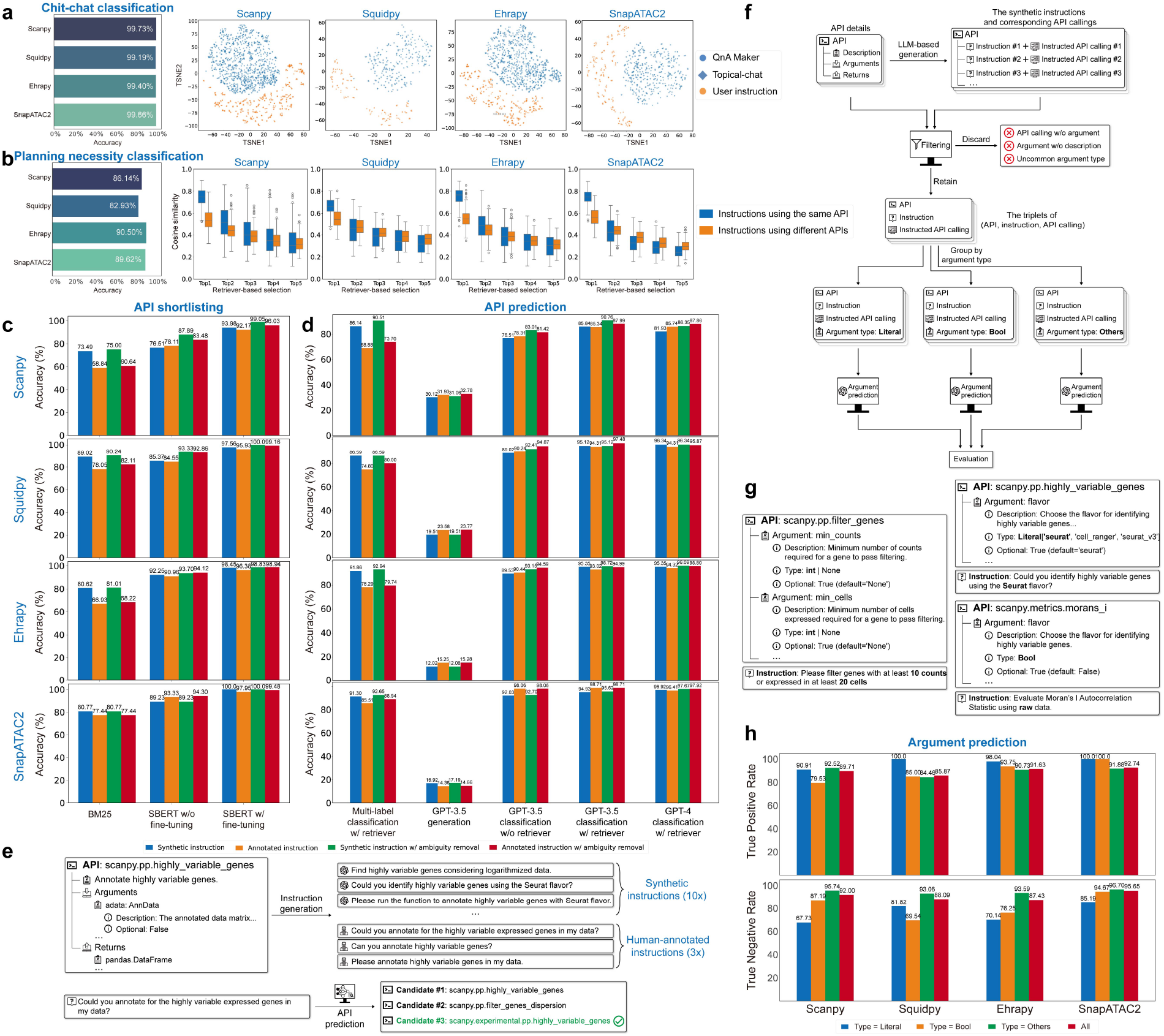
Evaluating the effectiveness of BioMANIA in each of its key components. (**a**) BioMANIA accurately differentiates between chit-chat samples and API-encoded instructions generated by LLMs across different Python tools. The performance is further supported qualitatively by the clear separation observed in the t-SNE projection. (**b**) BioMANIA accurately distinguishes between instructions that encode a single API and those that encode multiple APIs across different Python tools. The cosine similarity of the top-ranked matches reported by the API retriever, the top-3 in particular, reveals a significant disparity between instructions encoding a single API and those encoding multiple APIs. (**c**) BioMANIA uses a fine-tuned Sentence-BERT-based API retriever to accurately identify the top three candidate APIs containing the target, outperforming two baselines: BM25 and Sentence-BERT without fine-tuning. (**d**) BioMANIA frames API prediction as an in-context API selection task and outperforms other baselines in terms of accuracy, including alternative problem formulations (*e*.*g*., multi-label classification or sequence generation) and different LLM choices (*e*.*g*., GPT-3.5). (**e**) Here, we provide illustrative examples of synthetic instructions generated by LLMs for a randomly selected API, along with API retrieval and prediction for a randomly selected user instruction. (**f**) BioMANIA categorizes argument prediction into three groups based on argument type. (**g**) Each argument type is exemplified by a randomly selected example. (**h**) BioMANIA accurately predicts arguments of different types, achieving high true positive and true negative rates.

We next investigated BioMANIA’s performance distinguishes between instructions that encode a single API and those that encode multiple APIs (Refer to Sec. 4.1.3 for more details). We generated two population of API-encoded instructions: one consisting of concatenated instructions encoding the same API generated by LLMs, and the other consisting of concatenated instructions encoding multiple different APIs generated by LLMs. Qualitatively, the cosine similarity of the top-ranked matches reported by the API retriever reveals a significant disparity between instructions encoding a single API and those encoding multiple APIs (Fig. 3**b**). By fitting a multivariate Gaussian model using the cosine similarity values of the top-3 matches reported by the API retriever for each instruction population, the model accurately distinguishes between instructions that encode a single API and those that encode multiple APIs. This approach achieves accuracies of 86.14%, 82.93%, 90.50%, and 89.62% across four different Python tools, respectively. Accurate classification of planning necessity enables BioMANIA to better interpret the user’s intent and determine whether the task requires multiple API calls.

After initiating an API call, we evaluated BioMANIA’s performance in shortlisting the top three candidate APIs that contain the target (Refer to Sec. 4.1.4 for more details). BioMANIA employs a finetuned Sentence-BERT-based API retriever (Reimers and Gurevych, 2019) for shortlisting. We evaluated its performance by comparing it to two baseline methods: Sentence-BERT without fine-tuning and BM25, (Robertson et al, 2009), a commonly used deterministic relevance measure for document comparison. To assess whether the synthesized instructions generated by the LLMs accurately mimic the user’s instructions, we used two test datasets: one with synthesized instructions generated by the LLM and the other manually labeled by a diverse group of human annotators (Refer to Sec. 4.3 for more details). The fine-tuned API retriever achieves an average accuracy of 97.50% for synthetic instructions and 95.61% for human-annotated instructions in shortlisting the target API within the top three candidates across different Python tools (Fig. 3**c**). The slightly lower performance with human-annotated instructions is expected, as the test synthetic instructions were generated by GPT-4, which was also used during training. In contrast, Sentence-BERT without fine-tuning and BM25, two baseline methods that lack access to training data, achieve average accuracies of 85.84% and 80.98% for synthetic instructions, and 86.74% and 70.32% for human-annotated instructions, respectively. It is important to note that improper API design can hinder effective API shortlisting, with a significant portion of misclassifications attributed to the presence of ambiguously designed APIs (Refer to Sec. 2.1 for more details). By focusing on target APIs without ambiguity, the fine-tuned API retriever achieves an increased average accuracy of 99.47% for synthetic instructions and 98.40% for human-annotated instructions.

With the shortlisted APIs from the API retriever, we evaluated BioMANIA’s performance in predicting the target API from the shortlist (Refer to Sec. 4.1.5 for more details). We assessed prediction performance across multiple settings, considering factors such as the use of LLMs for prediction, the inclusion of the API Retriever, the formulation of the problem as either a prediction task or a sequence generation task (*i*.*e*., generating the API from scratch), and the choice between GPT-3.5 and GPT-4 (Fig. 3**d**). The in-context API classification model based on GPT-4 achieves an average accuracy of 92.64% for synthetic instructions and 92.70% for human-annotated instructions when predicting the target API from the top three candidates across different Python tools. After excluding ambiguous APIs, the average accuracy improves to 94.11% for synthetic instructions and 94.36% for human-annotated instructions.

Refer to Fig. 3**e** for illustrative examples of API retrieval and prediction given a user instruction. Furthermore, we observed that: (1) Utilizing the LLM for prediction significantly outperforms multi-label classification; (2) Framing the problem as a prediction task yields better results compared to formulating it as a sequence generation task; (3) Employing the API Retriever helps filter out incorrect answers, thereby improving prediction accuracy; and (4) GPT-4 better comprehends in-context demonstrations and predicts the target API more effectively than GPT-3.5.

Lastly, we evaluated BioMANIA’s performance in predicting the argument values based on the predicted API call (Refer to Sec. 4.1.6 for more details). To prevent the generation of confidently incorrect or incompatible values for API calls, we categorize argument prediction into three groups based on argument type (Fig. 3**e**): *Literal, Bool*, and other types. We quantified the argument prediction performance using two metrics: the Area Under the True Positive Rate (TPR) and the True Negative Rate (TNR). A high TPR reflects better sensitivity in predicting the desired argument, while a high TNR indicates stronger resistance against hallucinations. As a result, BioMANIA achieves an average accuracy of 97.24%, 89.57%, and 89.90% for TPR, and 78.47%, 81.91%, and 94.77% for TNR across different Python tools, respectively. These results suggest potential areas for improvement in future work, particularly in reducing LLM hallucinations.

It is important to note that while BioMANIA performs effectively across its key components, it is not without flaws, and errors at each stage of the pipeline may accumulate. To prevent these errors from compounding into major issues, BioMANIA, though designed to operate autonomously, still involves the user in the process. Users are encouraged to monitor task progression and actively interact with BioMANIA, fostering a collaborative approach.

### 2.3 BioMANIA enables conversational single-cell transcriptome data analysis

To assess the effectiveness of BioMANIA in real-world conversational data analysis, we began by examining single-cell gene expression data using the widely-adopted tool, Scanpy (Wolf et al, 2018). Scanpy (https://scanpy.readthedocs.io) is a widely-used toolkit for analyzing single-cell gene expression data across a range of functionalities (Refer to Sec. 4.2 for more details). We gathered 21 representative singlecell transcriptome data analysis tasks from Scanpy, covering a broad range of functionalities including data loading, preprocessing, quality control, dimensionality reduction, summary statistics, gene ranking, and various visualization techniques (Refer to Fig. A.7 for more details). Among these tasks, BioMANIA successfully completed 18 out of 21 using Scanpy, with the majority of tasks requiring minimal or no troubleshooting (Fig. 4**a**). In contrast, general-purpose LLMs, when used to perform the analysis from scratch, succeeded in only 5 out of 21 tasks (Refer to Sec. A.4 for more details).

**Fig. 4:**
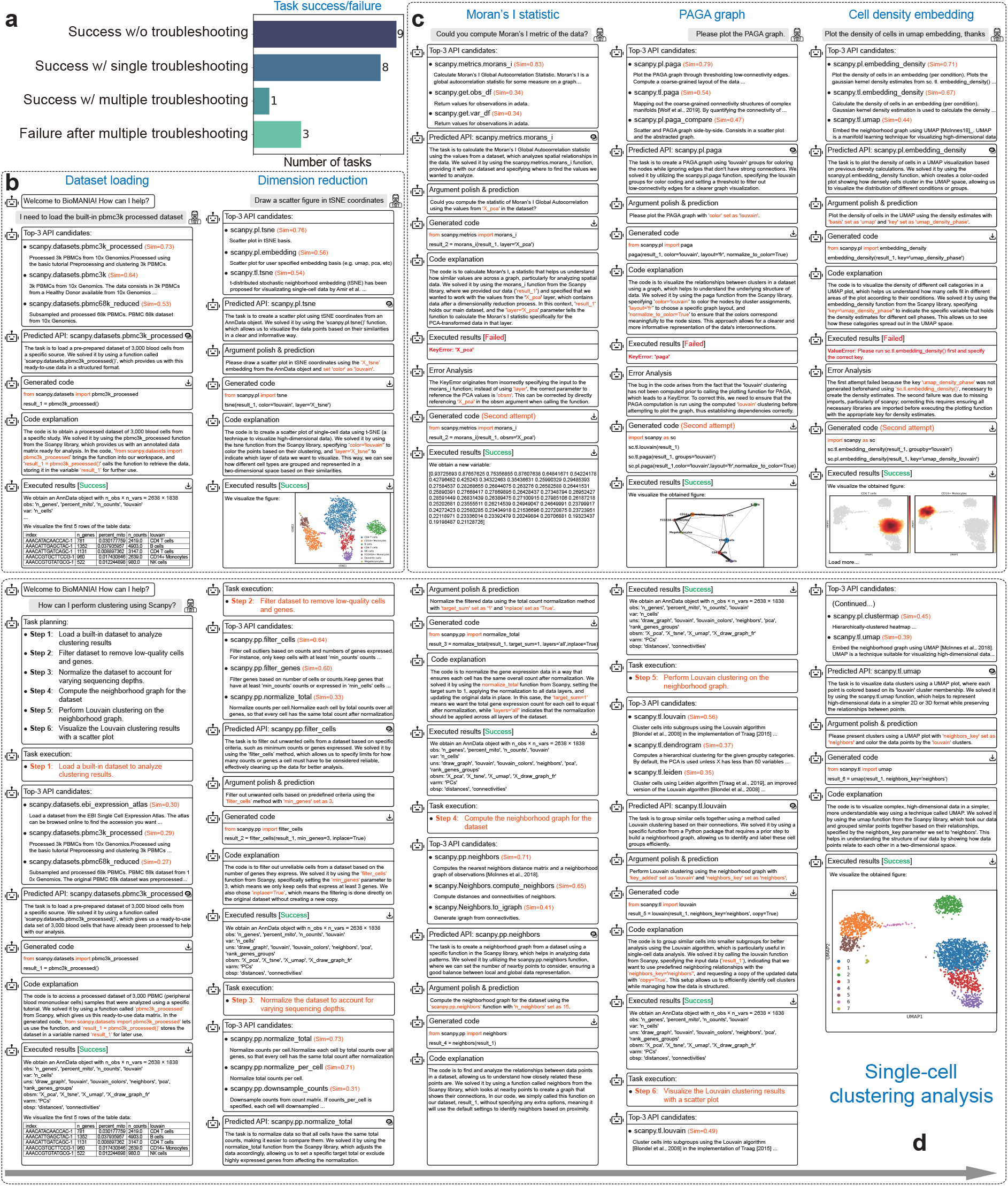
Illustrative examples of using BioMANIA for single-cell transcriptome data analysis. (**a**) BioMANIA successfully completed 18 out of 21 single-cell transcriptome data analysis tasks using Scanpy, with most tasks requiring little to no troubleshooting. (**b**) Successfully tasks accomplished by BioMANIA without any troubleshooting. (**c**) Successfully tasks accomplished by BioMANIA with troubleshooting. (**d**) BioMANIA is capable of task planning and decomposition, as demonstrated in a case study of single-cell clustering analysis.

At the start of the analysis, users can either upload their own datasets in the input area and instruct BioMANIA to parse them, or quickly load built-in datasets for a proof-of-principle analysis. After receiving instructions from users, BioMANIA lists the top three potential API candidates, each accompanied by a concise summary and the relevance score between the API and the input instruction, as determined by the fine-tuned API retriever. After predicting the target API from the shortlist, BioMANIA provides a summary based on the API information, explaining how the predicted API could address the user’s request and fulfill their needs. Once the target API and required arguments are identified, BioMANIA generates the corresponding code, including necessary library imports, and executes the API call using a built-in Python interpreter. After the API’s success execution, BioMANIA construct comprehensive natural language-based responses from three perspectives: (1) A step-by-step code explanation to illustrate each component for educational purposes, from the argument settings to the output; (2) The execution results in a user-friendly format, like a plotting figure or a snapshot of the data matrix; and (3) The API calls organized into a downloadable script for reproducibility, allowing users to customize the script and re-run it as needed. For example, BioMANIA can be instructed to load a processed 3k PBMC dataset from 10x Genomics and visualize the cells using t-SNE projection (Fig. 4**b**).

It’s important to note that BioMANIA cannot be guaranteed to be flawless, and errors may occur during API calls due to various reasons, such as incorrect formatting of input arguments or missing dependencies. When errors are encountered during code execution, BioMANIA refines the API calling code by gathering all relevant error information and prompting LLMs to troubleshoot and iteratively revise the code. For example, through troubleshooting, BioMANIA can pinpoint the erroneous argument needed for calculating Moran’s I Global Autocorrelation Statistic. For another example, it can identify that the root cause of a Key Error in plotting a trajectory-related PAGA graph (Wolf et al, 2019) lies in the missing invocation of dependent functions (Fig. 4**c**). While troubleshooting can help mitigate errors that occur during API calls, there remains a possibility that some issues may not be resolved effectively (Refer to Fig. A.7 for more details). Among the failed tasks, one involved creating a visual representation of pseudotime data using the “scanpy.pl.dpt groups pseudotime” function. The task encountered a Key Error due to missing dependencies. However, the challenge arose from the large number of prerequisite APIs that needed to be executed beforehand, and at times, the error messages were not informative enough to pinpoint the exact missing API call within the specified retry attempts. The other failed task involved annotating the genes with the most variation using the “scanpy.pp.highly_variable genes” function. Unfortunately, this task encountered a Key Error, and the error message provided little helpful information. The issue remains unresolved and is currently an open GitHub issue (https://github.com/scverse/scanpy/issues/2853).

Lastly, when the user provides a relatively vague instruction, BioMANIA is capable of understanding the user’s needs and breaking down the instruction into several tasks, addressing each one individually. For example, the user might vaguely request a clustering analysis on the data without specifying the exact strategy. BioMANIA can be prompted to break down the request into six sequential sub-tasks (Fig. 4**d**): (1) Load a built-in dataset; (2) Filter dataset to remove low-quality cells and genes; (3) Normalize the dataset to account for varying sequencing depths; (4) Compute the neighborhood graph to understand the relationships between data points in the dataset; (5) Perform Louvain clustering on the computed neighborhood graph; and (6) Visualize the Louvain clustering results with a scatter plot. The planned sub-tasks encompass the entire data analysis pipeline, including data loading, preprocessing, analysis, and visualization. Importantly, each sub-task is concrete and can be addressed sequentially through individual API calls. Refer to Fig. A.8 for more examples of single-cell transcriptome data task planning.

### 2.4 BioMANIA empowers conversational analysis across much broader domains

Although we have demonstrated BioMANIA for conversational analysis in single-cell transcriptome data, we believe that BioMANIA holds potential for broader applications across various domains. We next examined the effectiveness of BioMANIA in analyzing single-cell spatial omics data using a widely-adopted tool, Squidpy (Palla et al, 2022). Squidpy (https://squidpy.readthedocs.io) is a widely-used toolkit for analyzing single-cell spatial omics data (Refer to Sec. 4.2 for more details). We demonstrated the usefulness of BioMANIA in six representative single-cell spatial omics data analysis tasks, encompassing a broad range of functionalities, including data loading, Ripley’s statistics for spatial pattern analysis, co-occurrence probability of clusters, centrality scores for different cell types, neighborhood enrichment analysis, and cluster interaction analysis (Fig. 5). Since Squidpy separates data analysis from visualization, the six selected tasks effectively involve 12 API calls in total. These showcases present a new representative scenario where additional argument information is required, highlighting BioMANIA’s ability to handle such requests effectively. For example, one task involved plotting the centrality scores for different cell types, but the user neglected to mention the key that stores the cell type information in the conversation. BioMANIA accurately predicts the target API for plotting centrality scores but encounters an issue due to the absence of the key name that holds the cell type information. As a result, BioMANIA explains to the user what is needed to proceed and requests the additional argument information necessary for execution.

**Fig. 5:**
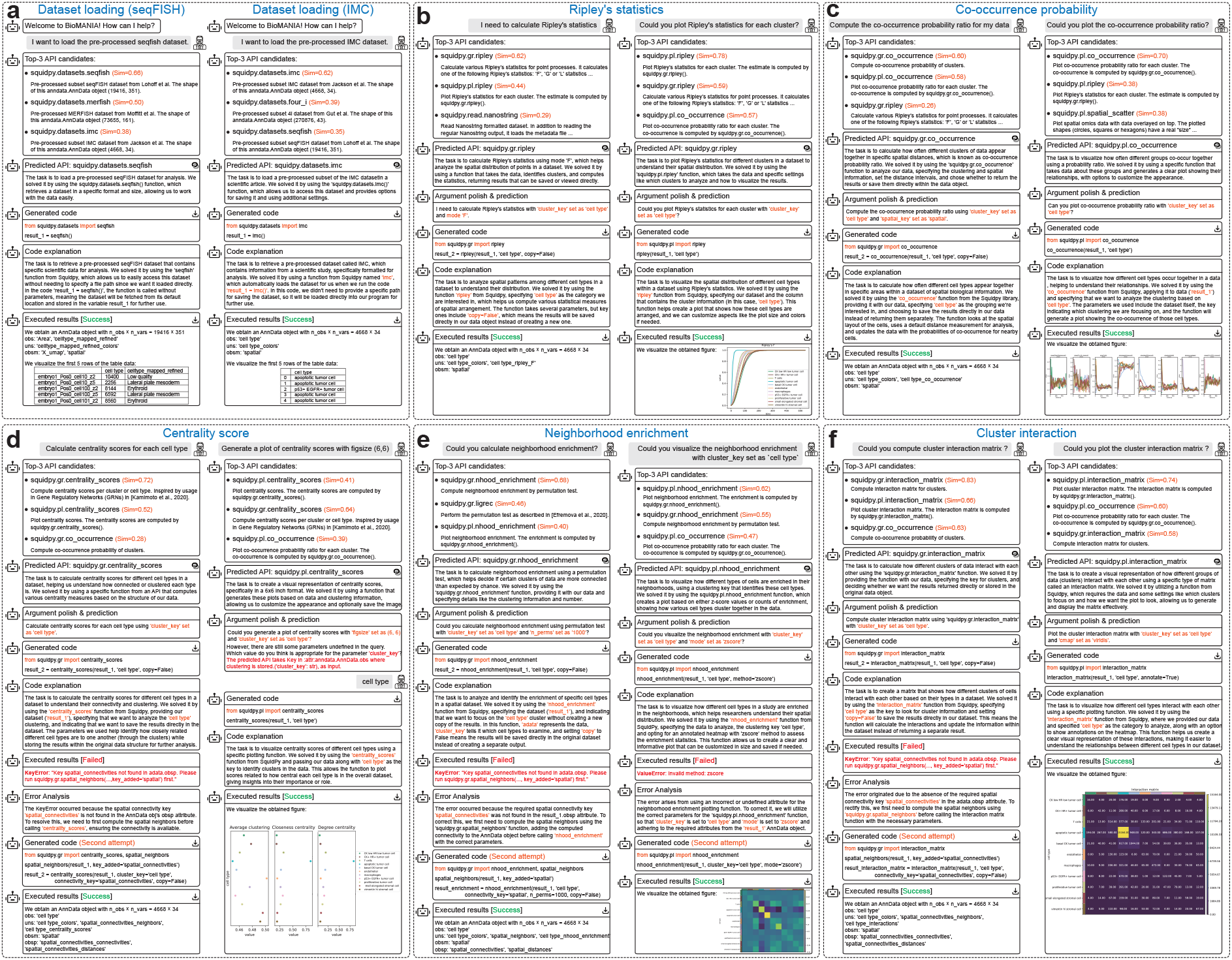
Illustrative examples of using BioMANIA for single-cell spatial omics data analysis. These examples encompass a wide range of single-cell spatial omics data analysis functionalities, including (**a**) Data loading, (**b**) Ripley’s statistics to analyze spatial patterns, (**c**) Co-occurrence probability of clusters, (**d**) Centrality scores for different cell types, (**e**) Neighborhood enrichment analysis, (**f**) Cluster interaction analysis.

In addition to single-cell spatial omics data analysis, We also examined the effectiveness of BioMANIA in analyzing single-cell epigenomics data using a widely-adopted tool, **SnapATAC2** (Zhang et al, 2024). SnapATAC2 (https://kzhang.org/SnapATAC2) is a widely-used toolkit for single-cell epigenomics analysis especially for single-cell ATAC-seq data (Refer to Sec. 4.2 for more details). We showcased the utility of BioMANIA in a lengthy yet commonly used pipeline for identifying enriched transcription factor (TF) motifs. This TF motif enrichment identification pipeline consists of several key tasks, including data loading, peak calling, peak merging, peak counting, marker region identification, TF motif retrieval, and TF motif enrichment analysis (Fig. 6). As a result, BioMANIA produces visualized motif enrichment results, aiding in the identification of crucial enriched motifs in the input data while maintaining a low false discovery rate.

**Fig. 6:**
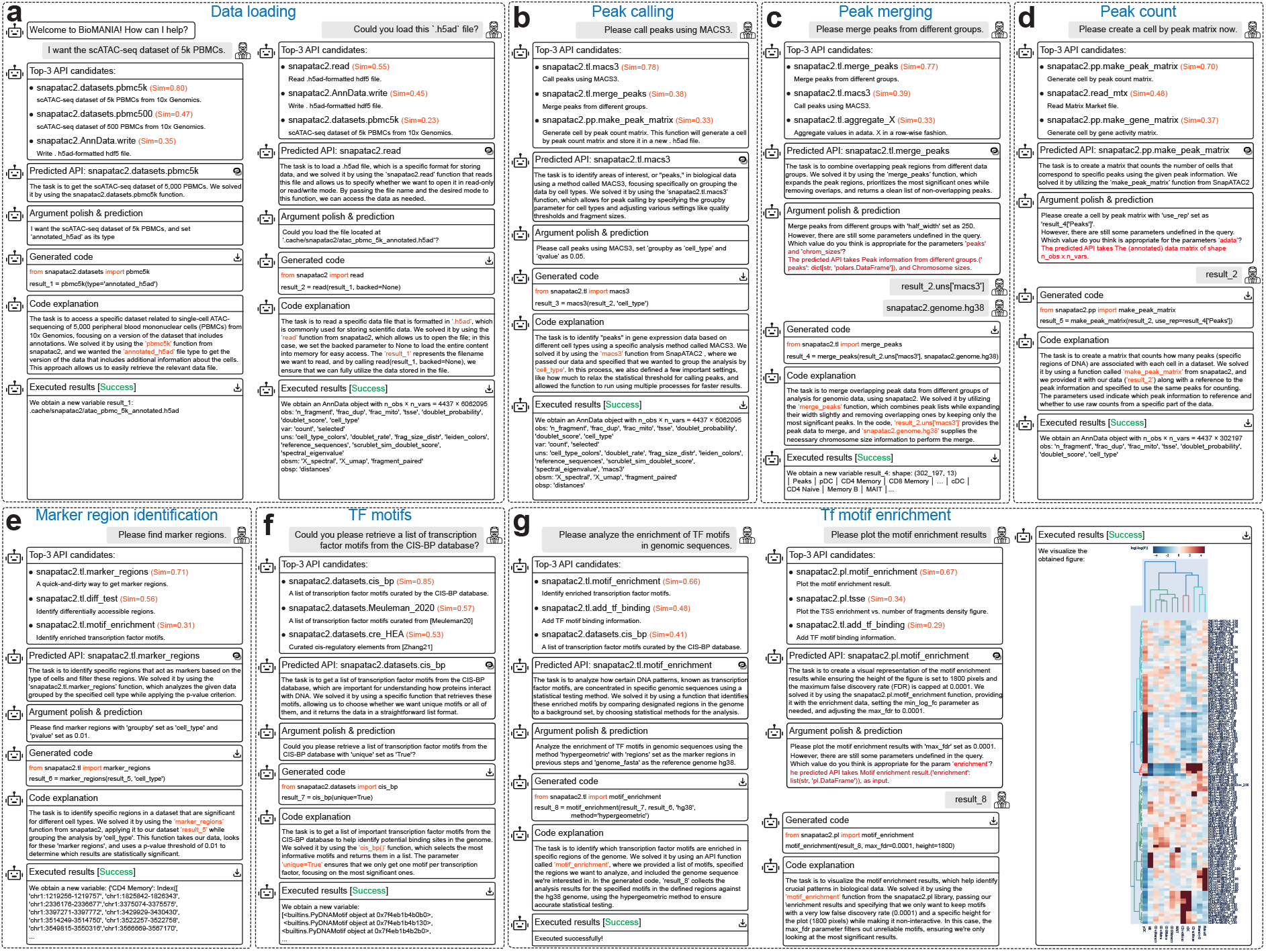
Illustrative examples of using BioMANIA for single-cell epigenomics data analysis. These examples encompass a wide range of single-cell epigenomics data analysis functionalities, including (**a**) Data loading, (**b**) Peak calling, (**c**) Peak merging, (**d**) Peak count, (**e**) Marker region identification (**f**) Transcription factor (TF) motifs retrieval, (**g**) Tf motif enrichment analysis.

Lastly, BioMANIA is applicable beyond the single-cell domain. We demonstrated its utility in electronic health record data analysis, showcasing a variety of tasks that encompass a broad range of functionalities, including data loading, missing value visualization, missing value imputation, and survival analysis. Refer to Fig. A.9 for more details of using **Ehrapy** (Heumos et al, 2024) for conversational electronic health record data analysis.

## 3 Discussion

In this study, we aim to bridge the learning barriers in data analysis for biological scientists, addressing the need for substantial technical skills and domain-specific expertise. For this purpose, we propose BioMANIA, an AI-driven, natural language-oriented bioinformatics data analysis pipeline that facilitates the automatic execution of off-the-shelf Python-based bioinformatics tools, enabling the generation of analysis results for biological data with minimal tool-specific knowledge and coding expertise required. The key novelties of BioMANIA lie in its design, where BioMANIA explicitly learns about API information and their interactions directly from the source code and documentation of any well-documented, open-source Python tools. This allows for a deeper, more accurate understanding of the tools’ functionalities, enabling seamless integration into the conversational analysis pipeline. BioMANIA follows a rigorous and comprehensively validated process: interpreting the user’s intent, breaking it down into specific tasks when necessary, mapping these tasks to API calls, executing the APIs with inferred arguments, and troubleshooting any coding errors encountered during execution. Compared to using LLMs to conduct analysis from scratch, which can lead to hallucinated responses and potential misdirection, the structured steps in BioMANIA help mitigate hallucinations, significantly reducing the likelihood of confusion and errors. This approach ensures a more reliable and accurate data analysis process.

Traditionally, LLM-driven bioinformatics data analysis tools are specialized and designed for predefined tasks, lacking the flexibility to easily extend to new, unseen application domains (Li et al, 2021; Xiao et al, 2024). This rigid structure limits their adaptability, making it challenging to apply them to emerging datasets, novel analytical methods, or other areas that require dynamic adjustments beyond their original design. This contrasts with approaches like BioMANIA, which offer more versatility by leveraging open-source Python tools and adapting dynamically to user queries across diverse domains. In this way, our approach provides a path toward end-to-end automatic and codeless execution of any off-the-shelf biological analysis workflow. Furthermore, BioMANIA takes full consideration of the goals of reproducibility and education. It provides user-friendly, detailed descriptions for each step of the analysis, ensuring that users understand the reasoning behind every API call. BioMANIA organizes these API calls into a downloadable script for easy reuse, offers step-by-step explanations of the code, and generates a comprehensive report documenting the entire data analysis process. This approach not only ensures transparency and reproducibility but also serves as an educational tool, helping users to better understand and learn from the analysis. We have conducted comprehensive evaluation to demonstrate BioMANIA’s empirical utility. we applied BioMANIA to a range of commonly used bioinformatics tools, covering diverse application domains–from single-cell RNA-seq analysis (Wolf et al, 2018), single-cell spatial omics analysis (Palla et al, 2022), single-cell epigenomics analysis (Zhang et al, 2024), to electronic health record analysis (Heumos et al, 2024).

Lastly, this study highlights several promising directions for future research. First of all, we recognize that off-the-shelf Python tools were not originally designed for conversational analysis. The Python tools used by BioMANIA exhibit varying levels of API design ambiguity, validity, and complexity, which are demonstrated to influence the effectiveness of conversational data analysis. These factors can lead to challenges such as ambiguous function names, excessive and poorly documented parameters, or outdated documentation, all of which can hinder accurate API prediction, argument selection, and overall performance of the conversational interface. This limitation highlights an area for future development. For example, BioMANIA currently employs a static Abstract Syntax Tree parser to analyze source code and extract potential APIs. While this approach helps identify individual APIs and their arguments, it primarily focuses on static code analysis. It would be interesting to employ a dynamic parser, which could analyze runtime behavior and the interplay among APIs. This would enable BioMANIA to aggregate multiple APIs into higher-level functionalities that align more closely with the user’s needs, offering a more seamless and intuitive data analysis experience. Secondly, BioMANIA currently offers a preliminary task planning step to decompose the user’s instruction into several tasks and address each one individually. However, this approach implicitly assumes that the user has some knowledge of the scope and capabilities of the Python tool being used, providing only a vague description of their needs. This differs from the scenario of engaging with unknown data for data-driven scientific discovery, where the user’s expectations might be less defined. For future development, it would be interesting to build a pool of specialized chatbots, with each chatbot representing a different Python tool and deeply understanding its capabilities. A more powerful, overarching planner could then coordinate these chatbots, enabling them to work together to better explore the data, identify patterns, and make scientific discoveries. This system would offer a more dynamic, multi-tool approach, leveraging the strength of each tool to guide the user through more complex analyses and discoveries, even with unknown datasets.

Existing Python tools passively analyze the provided data and perform their predefined functionalities. However, in many cases, researchers face the challenge of insufficient data, as generating data through experiments can be costly and time-consuming (Huang et al, 2024). From the practitioner’s perspective, it would be highly beneficial to develop an agent capable of actively exploring the data that has been acquired so far. This agent could reverse the process by guiding optimal experimental design to generate better, more informative data. By incorporating both data generation and analysis within the same feedback loop, such an agent could assist in designing more efficient experiments that maximize the insights gained from minimal data. This integration would empower practitioners to not only analyze but also proactively plan for future data collection, leading to more targeted, cost-effective, and scientifically valuable experimental designs. Combining data generation, analysis, and experimental planning would create a unified system for comprehensive data-driven research.

In conclusion,BioMANIA simplifies the exploration of omics data and accelerates bioinformatics research and enables more accurate and comprehensive biological discoveries. Also, BioMANIA significantly lowers the technical barriers to conducting state-of-the-art biological data analysis, with the potential to usher in a new research paradigm. We anticipate that BioMANIA will reshape the landscape of bioinformatics research, enhancing data analysis accessibility and expediting discoveries in the field.

## 4 Methods

### 4.1 The key components of BioMANIA

BioMANIA relies on several key components to enable automated, codeless biological data analysis (Fig. 1). Through the front-end chatbot interface, these components work together effectively and seamlessly to interpret the user’s intent, plan analysis tasks, translate them into specific API calls with concrete arguments, and execute the tasks while troubleshooting any runtime errors.

#### 4.1.1 Tool parsing

BioMANIA is designed to directly use natural language to perform complex biological data analysis, leveraging existing software libraries and workflows to accomplish these tasks. To achieve this goal, BioMANIA explicitly learns API information and their interactions from the source code (*e*.*g*., hosted in GitHub) and documentation (*e*.*g*., hosted in ReadTheDocs) of any off-the-shelf well-documented, opensource Python tools. Specifically, BioMANIA employs an Abstract Syntax Tree (AST) parser to analyze the source code and extract potential APIs. For each API, we further extract key details, including its requirements (argument list), operations (name, description), and return values, to ensure its effective utilization. Arguments can be either required or optional, with each having an argument name, description, data type, and, for optional arguments, a default value.

Source code analysis often yields a large number of APIs with varying quality and reliability. BioMANIA employs a rigorous filtering process to remove low-quality and unreliable APIs from direct source code analysis. We start by removing APIs that lack documentation, are not executable, or are deprecated Furthermore, BioMANIA cross-compares the curated list of APIs from the tool’s documentation with those identified from the source code, retaining only the APIs that appear in both. Finally, BioMANIA double-checks the remaining APIs by testing their functionalities to ensure they are fully operational.

#### 4.1.2 Chit-chat recognition

Based on the user’s instruction, BioMANIA determines their intent by assessing whether they are asking an API-specific data analysis question. In natural conversation, a significant portion of the dialogue does not convey user intent – instead, it consists of greetings or remarks that do not request information or program execution (Stewart et al, 2006). Consequently, we incorporate a chit-chat detection stage to determine whether the user is inquiring about API-based data analysis or simply engaging in casual conversation. To differentiate between API calls and casual conversation, we compile a substantial corpus of open-domain conversational samples from two existing chit-chat datasets: Topical-Chat (Gopalakrishnan et al, 2019) and Qnamaker (Agrawal et al, 2020). These two datasets complement each other, with the Topical-Chat dataset covering across different topics (*e*.*g*., Sports, Music, Science and Technology, etc.) and the Qnamaker dataset focusing on various personalities (*e*.*g*., enthusiastic, professional, witty, etc.). The goal of chit-chat detection is to differentiate between chit-chat samples and API-encoded instructions generated by LLMs. Specifically, we represent both chit-chat samples and API-encoded instructions as TF-IDF embeddings (Ramos, 2003). This process involves vectorizing the TF-IDF (Term Frequency-Inverse Document Frequency) scores, a widely used metric for determining word importance, for each word in the dictionary. We calculate the centroids for three groups (*i*.*e*., API-encoded instructions, Topical-Chat samples, and Qnamaker samples) and use these to build a nearest centroid classifier for chit-chat detection. For a new user instruction, we compute the cosine similarity between its TF-IDF embedding and these centroids, predicting the label based on the closest centroid. Despite its simplicity, we found that the nearest centroid classifier performs both accurately and efficiently in practice (Refer to Fig. A.2 for more details). Finally, chit-chat detection leads to one of two outcomes: either providing a chit-chat response or recognizing the API from the question. Accordingly, the chatbot will either use an LLM to engage with users or proceed to determine the corresponding API calls (Fig. 1**b**).

#### 4.1.3 Task planning

If the user is asking an API-specific data analysis question, BioMANIA further evaluates whether the task requires multiple API calls. We frame the assessment of whether multiple API calls are required as a binary classification problem. To achieve this, we generate two populations of API-encoded instructions: one consisting of concatenated instructions encoding the same API generated by LLMs, and the other consisting of concatenated instructions encoding multiple different APIs generated by LLMs. Since concatenated instructions are generally longer than single instructions, we use LLMs to shorten them to prevent instruction length from becoming a confounding factor. Each concatenated instruction, from either population, is searched against an API retriever, which encodes the instruction and API documentation (including its name, description, and argument list) as embeddings (Refer to Sec. 4.1.4 for more details). The relevance between the instruction and API is determined by the cosine similarity between the two embeddings. We observed that the cosine similarity of the top-ranked matches reported by the API retriever reveals a significant disparity between instructions encoding a single API and those encoding multiple APIs (Refer to Fig. A.3 for more details). Following this observation, we fit a multivariate Gaussian model using the cosine similarity values of the top-3 matches reported by the API retriever for each instruction population. For a new user instruction, we search it against the API retriever, obtain the cosine similarity values of the top matches, compute the probability under both fitted Gaussian models, and determine the necessity of multiple API calls based on which model having a higher probability.

If multiple API calls are required, BioMANIA breaks the user’s instruction into several tasks and addresses each one individually. Due to the inherent dependencies among certain tasks, the execution sequence of the decomposed tasks must be considered to establish interconnections between the tasks effectively. We provide the following information as part of the prompt to the LLMs: (1) A description of the task goal, including details about the input data and the desired output format; (2) The complexity and scope of each subtask; and (3) Guidelines for describing the subtasks. Based on the LLM’s built-in knowledge and the descriptions provided for task planning, the LLM intelligently understands the user’s instruction and breaks it into a sequence of tasks, while considering task dependencies. Detailed prompts for task planning, including examples, can be found in Sec. A.5.1.

#### 4.1.4 API shortlisting

After planning, BioMANIA aims to complete all the tasks sequentially by translating each task into the corresponding API call. However, predicting the most suitable API from the user’s instruction is challenging because real-world bioinformatics tools often have a vast number of API candidates. This makes it impractical to include descriptions of all APIs as input for LLMs due to token length and cost constraints. To address this issue, we implement an API shortlisting process by using an API retriever to prune candidate APIs. When there are too many APIs, the retriever identifies the top-k most relevant APIs to present to the LLMs.

Shortlisting the relevant API from user instructions typically requires a large-scale dataset of instruction-API pairs (Chen et al, 2021). Instead of collecting such data through human annotation, which is expensive and time-consuming, we automatically synthesize language instructions given information about an API (Wang et al, 2022). Specifically, we provide LLMs with the API name, description, and argument list as input and prompt them to generate five varied, detailed, and verbose instructions for making API calls. To improve robustness and generalization across different usage scenarios, we also generate five precise and concise instructions. In total, we synthesize ten language instructions for each API. Detailed prompts for instruction generation, including examples, can be found in Sec. A.5.4.

With instruction-API pairs available, we train an API retriever that encodes both the instruction and API documentation as embeddings. The relevance between an instruction and API is then determined by the cosine similarity between these embeddings. The retriever is fine-tuned from a Sentence-BERT model (Reimers and Gurevych, 2019) to maximize the similarity of matching API-instruction pairs and minimize the similarity of randomly sampled negative API-instruction pairs. Consequently, for a new user instruction, the trained API retriever generates a ranked list of APIs based on embedding-based cosine similarity, where higher-ranked APIs are considered more relevant to the user’s instruction (Refer to Fig. A.4 for more details).

#### 4.1.5 API prediction

With the shortlisted APIs from the API retriever, the API predictor further refines the list to identify the best match. For this purpose, we frame API prediction as an in-context API selection task. Specifically, we supply the API shortlist alongside several demonstrations, where each demonstration includes a synthetic instruction paired with its corresponding API information. These randomly selected demonstrations assist LLMs (*e*.*g*., GPT-4) in grasping the underlying relationships between instructions and APIs, thereby enabling more accurate API predictions. It is worth noting that in the tool parsing stage (Sec. 4.1.1), we faced challenges related to API design, such as different APIs having similar descriptions or identical names, which introduced ambiguity and could negatively affect API prediction accuracy (Refer to Sec. 2.1 for more details). In cases where ambiguous APIs with similar descriptions or identical names are identified, we generate an ambiguity list for each API and prompt users for clarification to determine the correct target API. Detailed prompts for API prediction, including examples, can be found in Sec. A.5.5.

#### 4.1.6 Argument prediction

To generate an executable API call, we further predict both the required and optional argument values based on the predicted API call. For each argument, we supply the LLMs with the user instruction, the predicted API (including its name and description), and details of the argument under consideration (such as its name, description, data type, and default value), along with a list of possible values. A key prerequisite for accurately predicting argument values is ensuring that the LLMs strictly follow the required argument format, avoiding the generation of confidently incorrect or incompatible values for the API call. To accomplish this, we categorize argument types into three groups: (1) The *Literal* type, where the choice is restricted to a small set of predefined values; (2) The *Bool* type, where the choice is “true” or “false”, depending on whether the argument is present; and (3) Other types, which include “string”, “int”, “float”, and similar basic data types. LLMs are instructed to handle each group separately, selecting the argument’s value from the user instruction or returning the default value if no prediction is made (Refer to Fig. A.6 for more details).

It’s important to highlight the challenges of argument prediction in BioMANIA, as not all arguments can be inferred directly from the user’s instruction. For example, predicting a Pandas dataframe or Numpy multidimensional array based solely on user input is nearly impossible. We reasoned that such information should be derived from earlier API call events in the conversation. Therefore, for these more complex arguments, rather than predicting them, we extract variables from the stack traces of previously executed API calls that match the required argument type. Detailed prompts for argument prediction, including examples, can be found in Sec. A.5.6.

#### 4.1.7 API execution

With the API and arguments predicted, BioMANIA uses a Python interpreter to execute the API call. However, calling errors are to be expected due to various reasons, such as incorrect formatting of input arguments or missing dependencies. Therefore, it is crucial to have robust troubleshooting mechanisms in place to continuously resolve these errors until successful code execution is achieved. BioMANIA is designed to refine API calling code by collecting all relevant error information when a call fails and prompting LLMs to troubleshoot and iteratively revise the code. The collected error information includes the error message, relevant Git issues, a description of the task and API, execution history, and other related details. Drawing on their built-in knowledge and the provided error information, the LLMs can intelligently diagnose the cause of the error and revise the code accordingly. Detailed prompts for code troubleshooting, including examples, can be found in Sec. A.5.8.

After the API’s success execution, BioMANIA construct comprehensive natural language-based responses from three perspectives. First, BioMANIA integrates all information from previous stages into a concise summary, encompassing API prediction and execution from API execution. Refer to Sec. A.5.3 for detailed prompts for task summarization). Second, BioMANIA presents the execution results in a user-friendly format, like a plotting figure or a snapshot of the data matrix. Third, BioMANIA organizes the API calls into a downloadable script for reproducibility and generates a comprehensive report recording the entire data analysis process. In addition, BioMANIA offers step-by-step explanations of the generated code for educational purposes. Detailed prompts for code explanation, including examples, can be found in Sec. A.5.7.

### 4.2 Tools used for evaluation

To evaluate the effectiveness of BioMANIA, we applied BioMANIA to a range of commonly used bioinformatics tools, covering diverse application domains (Refer to Fig. A.1 for more details):

- **Scanpy** (Wolf et al, 2018). Scanpy (https://scanpy.readthedocs.io) is a widely-used toolkit for analyzing single-cell gene expression data across a range of functionalities. Scanpy offers 146 valid and executable Python-based APIs for a range of analyses, including preprocessing, visualization, clustering, trajectory inference, and differential expression testing. Out of 146 APIs, 81 are covered by Scanpy tutorials and 123 are covered by unit-test examples to demonstrate their usage, respectively.
- **Squidpy** (Palla et al, 2022). Squidpy (https://squidpy.readthedocs.io) is a widely-used toolkit for analyzing single-cell spatial omics data. Squidpy offers 41 valid and executable Python-based APIs for various analyses, including spatial graph-based analysis, spatial statistics computation, and spatial image processing, among others. Out of 41 APIs, 38 are covered by Squidpy tutorials and 28 are covered by unit-test examples to demonstrate their usage, respectively.
- **SnapATAC2** (Zhang et al, 2024). SnapATAC2 (https://kzhang.org/SnapATAC2) is a widely-used toolkit for single-cell epigenomics analysis. SnapATAC2 offers 80 valid and executable Python-based APIs for a range of single-cell ATAC-seq data analyses, including preprocessing, dimension reduction, clustering, data integration, peak calling, differential analysis, motif analysis, regulatory network analysis. Out of 80 APIs, 31 are covered by SnapATAC2 tutorials and 23 are covered by unit-test examples to demonstrate their usage, respectively.
- **Ehrapy** (Heumos et al, 2024). Ehrapy (https://ehrapy.readthedocs.io) is a recent toolkit for analyzing heterogeneous epidemiology and electronic health record data. Ehrapy offers 130 valid and executable Python-based APIs for a range of analyses, including patient fate analysis, survival analysis, causal inference, among others. Out of 130 APIs, 46 are covered by Ehrapy tutorials and 101 are covered by unit-test examples to demonstrate their usage, respectively.

### 4.3 Human annotation

Several key components of BioMANIA, including chit-chat recognition (Sec. 4.1.2), task planning (Sec. 4.1.3), and API shortlisting (Sec. 4.1.4), rely on synthesized instructions generated by the LLMs. BioMANIA implicitly assumes that the synthesized instructions generated by the LLMs can accurately mimic the user’s instructions. However, whether this assumption holds true in real-world scenarios remains uncertain. To assess the extent to which this assumption is violated, we assembled a diverse group of human annotators to manually label three different instructions for each API across various tools, creating an independent test set. To provide a comparative perspective, we used GPT-4 to generate five detailed and five concise synthetic instructions with varying tones for each API. For each API, eight out of the ten synthetic instructions were assigned to the training split to train the API retriever and predictor, while the remaining two were reserved for evaluation.

### 4.4 Code and data availability

The software implementation in this study are available at https://github.com/batmen-lab/BioMANIA. The Docker images for the tools used by BioMANIA are available at https://hub.docker.com/repositories/chatbotuibiomania. The usage demos of BioMANIA are available at https://biomania.ngrok.io/en. The datasets and the models are available at https://drive.google.com/uc?id=1nT28pIJ dsdvi2yD8ffWt aePXsSWdqI.

## Declarations

- The research is supported by the Canadian NSERC Discovery Grant RGPIN-03270-2023.
- The authors declare that they have no conflict of interest.
- There is no ethics approval and consent to participate involved in this study.
- There is no consent for publication involved in this study.
- Author contribution: Z.D. implemented the code and did the analysis. Z.D., H.Z, and Y.J. annotated the API-specific instructions. Z.D. and Y.L. prepared the figures. Z.D., V.Z, and Y.L. participated the discussion. Y.L. wrote the manuscript. All authors reviewed the manuscript.

## A Appendix

### A.1 Details of the tools used by BioMANIA

**Fig. A.1:**
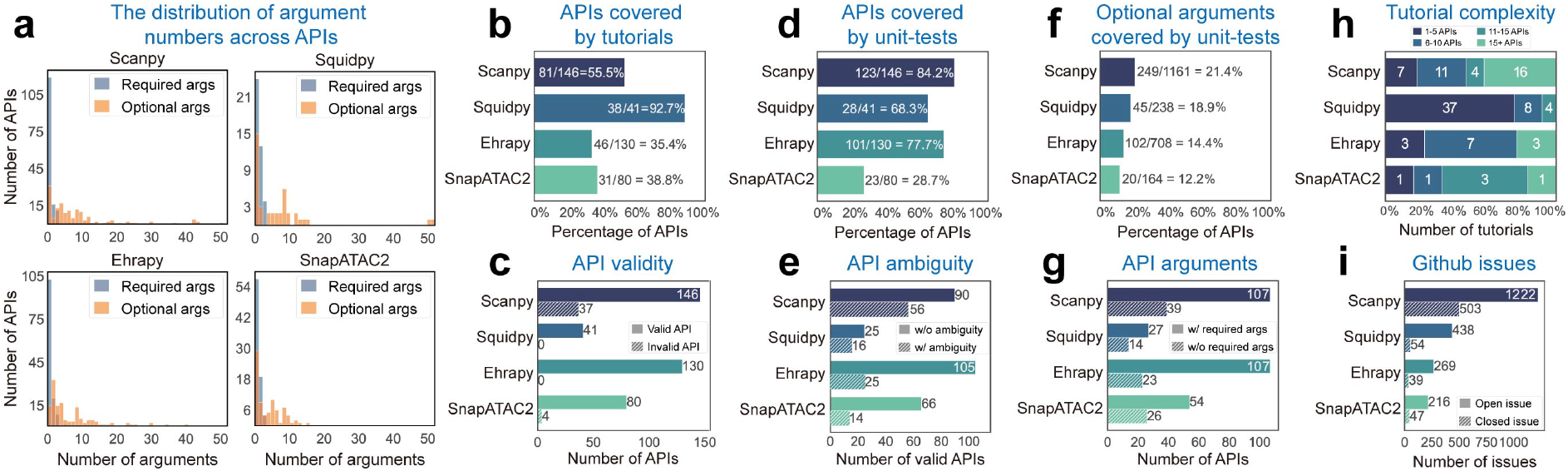
Detailed statistics for the Python tools used by BioMANIA. These statistics include: (**a**) The number of API arguments, (**b**) The number of APIs covered by tutorials, (**c**) The number of valid APIs, (**d**) The number of APIs covered by unit-tests, (**e**) The number of ambiguous APIs, (**f**) The number of optional arguments covered by unit-tests, (**g**) The number of APIs having required arguments, (**h**) The complexity of tutorials in terms of the API composition, and (**i**) The number of Git issues.

### A.2 Evaluating the performance of BioMANIA key components

**Fig. A.2:**
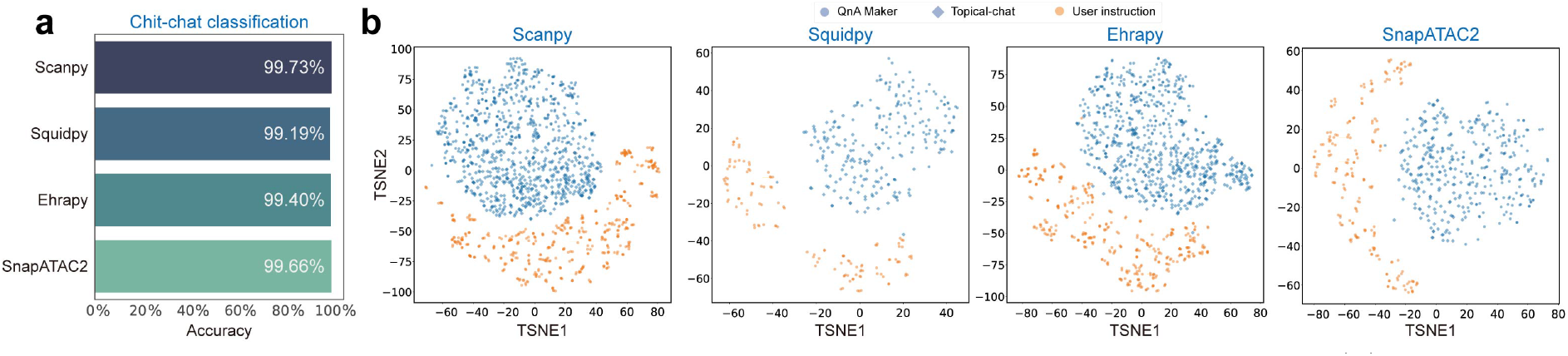
Evaluating the effectiveness of BioMANIA in chit-chat recognition. (**a**) BioMANIA accurately differentiates between chit-chat samples and API-encoded instructions generated by LLMs across different Python tools. (**b**) The performance is further supported qualitatively by the clear separation observed in the t-SNE projection.

**Fig. A.3:**
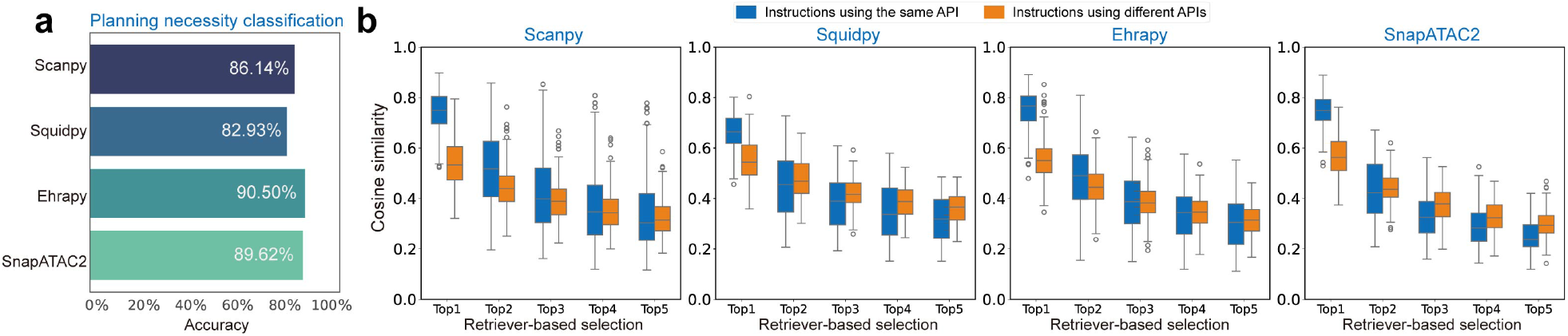
Evaluating the effectiveness of BioMANIA in planning necessity classification. (**a**) BioMANIA accurately distinguishes between instructions that encode a single API and those that encode multiple APIs across different Python tools. (**b**) The cosine similarity of the top-ranked matches reported by the API retriever reveals a significant disparity between instructions encoding a single API and those encoding multiple APIs. The disparity diminishes after the top three ranked matches.

**Fig. A.4:**
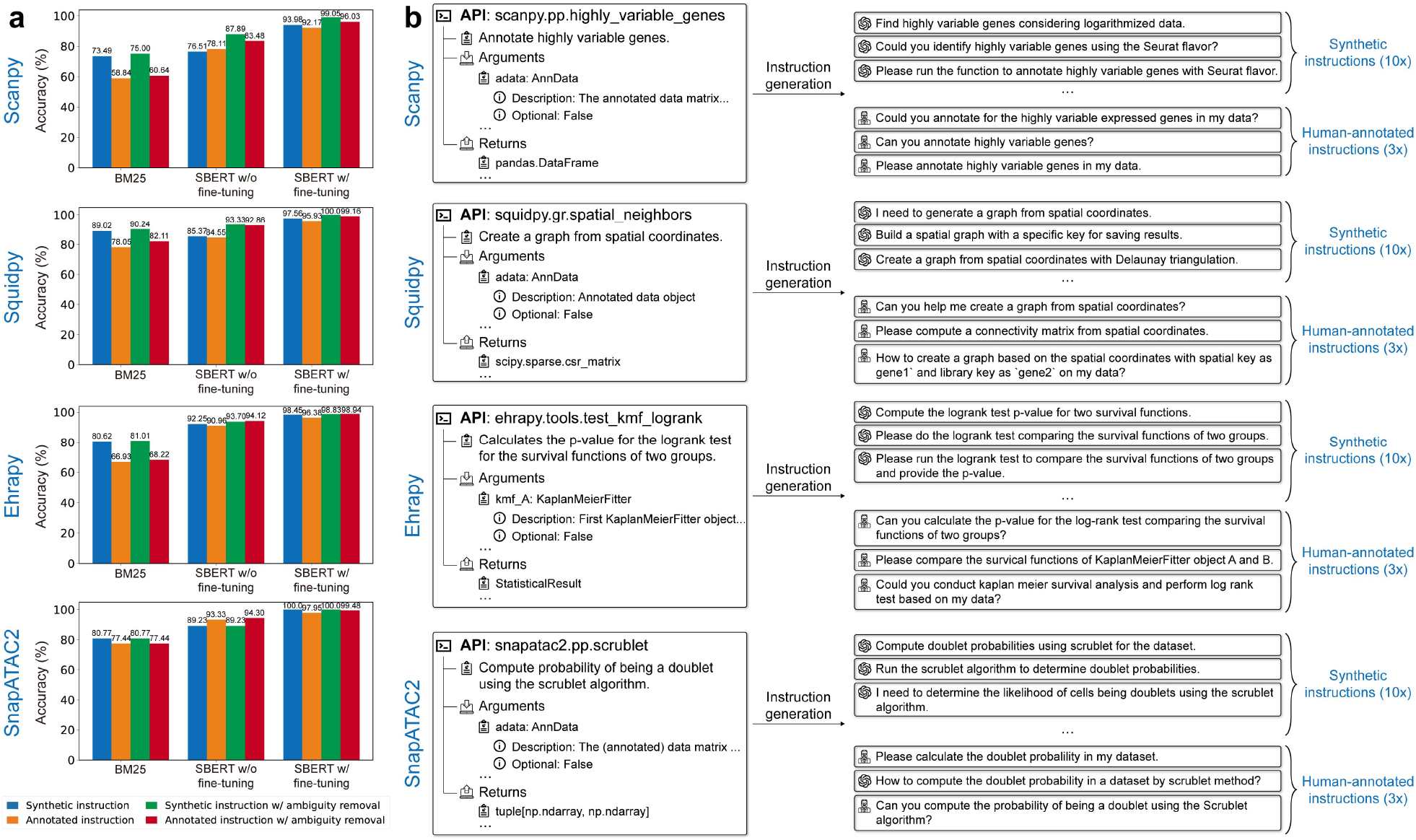
Evaluating the effectiveness of BioMANIA in API shortlisting. (**a**) BioMANIA uses a fine-tuned Sentence-BERT-based API retriever to accurately identify the top three candidate APIs containing the target, outperforming two baselines: BM25 and Sentence-BERT without fine-tuning. (**b**) Fine-tuning the Sentence-BERT model requires synthesized instructions based on information about an API. Here, we provide illustrative examples of synthetic instructions generated by LLMs for randomly selected APIs across different Python tools.

**Fig. A.5:**
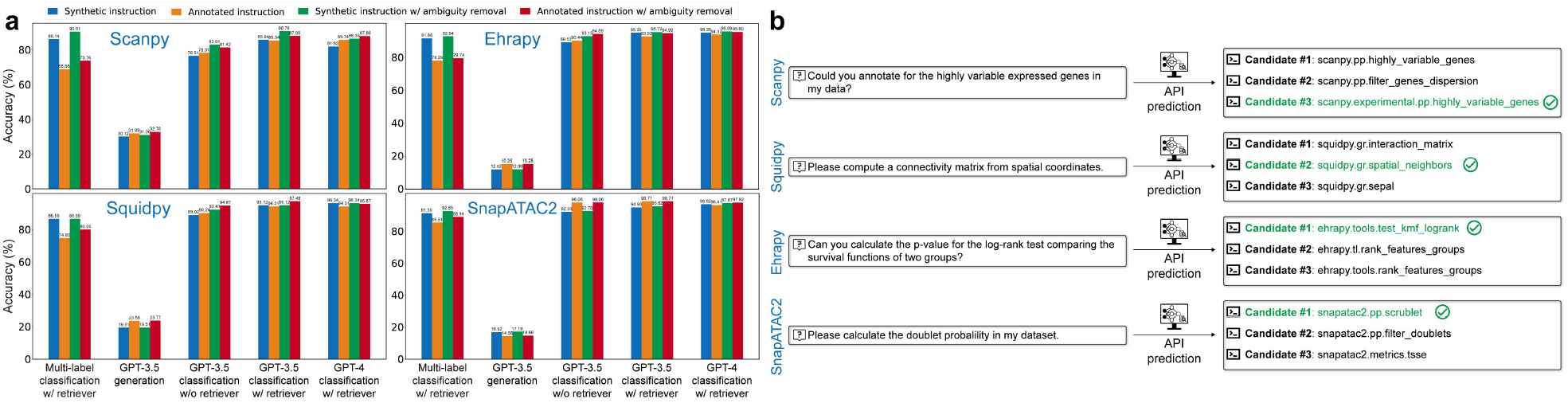
Evaluating the effectiveness of BioMANIA in API prediction. (**a**) BioMANIA frames API prediction as an in-context API selection task and outperforms other baselines in terms of accuracy, including alternative problem formulations (*e*.*g*., multi-label classification or sequence generation) and different LLM choices (*e*.*g*., GPT-3.5). (**b**) We also provide illustrative examples of API retrieval and prediction for randomly selected user instructions across different Python tools.

**Fig. A.6:**
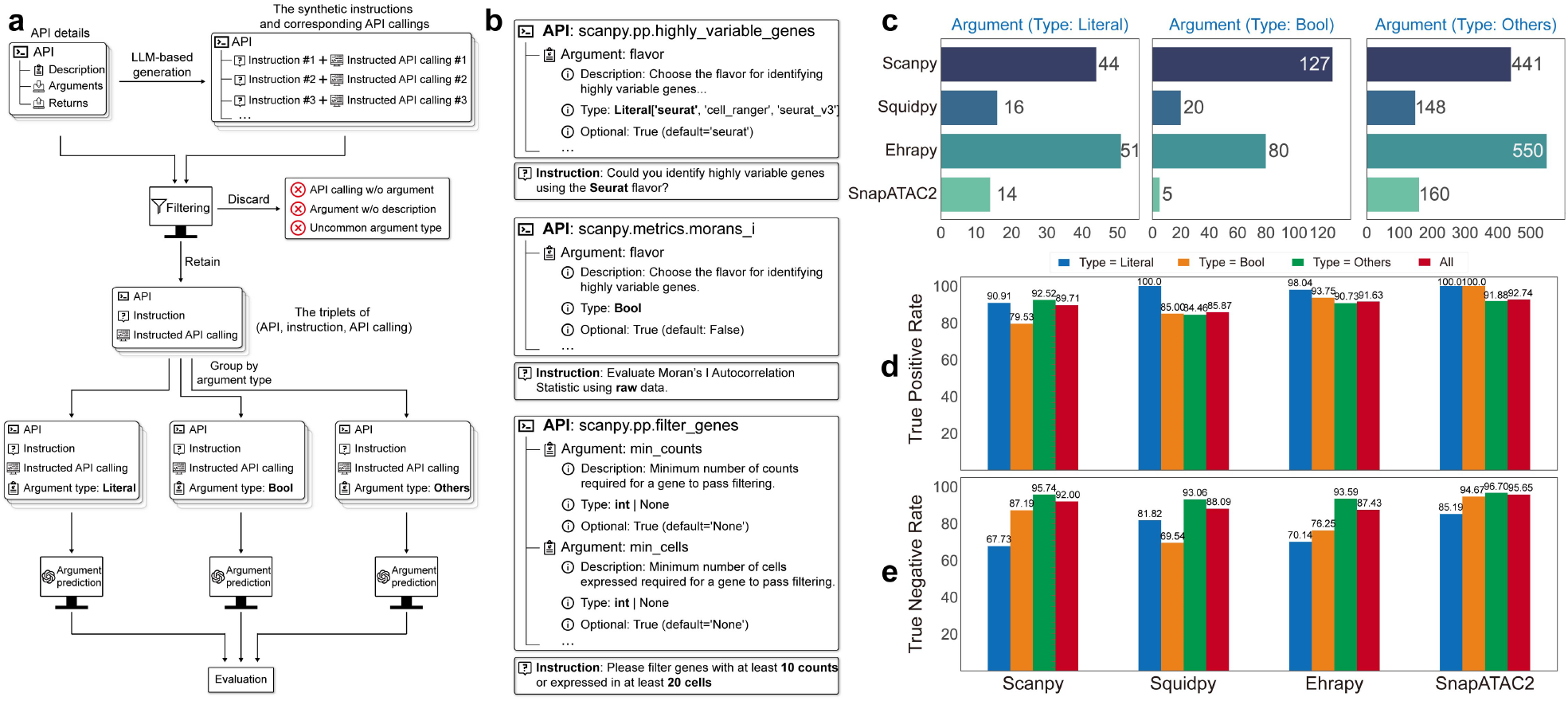
Evaluating the effectiveness of BioMANIA in argument prediction. (**a**) BioMANIA categorizes argument prediction into three groups based on argument type. (**b**) Each argument type is exemplified by a randomly selected example. (**c**) Each argument type appears unevenly in the instructions generated by LLMs across different Python tools. BioMANIA accurately predicts arguments of different types, achieving high (**d**) true positive and (**e**) true negative rates.

### A.3 Case studies of BioMANIA

**Fig. A.7:**
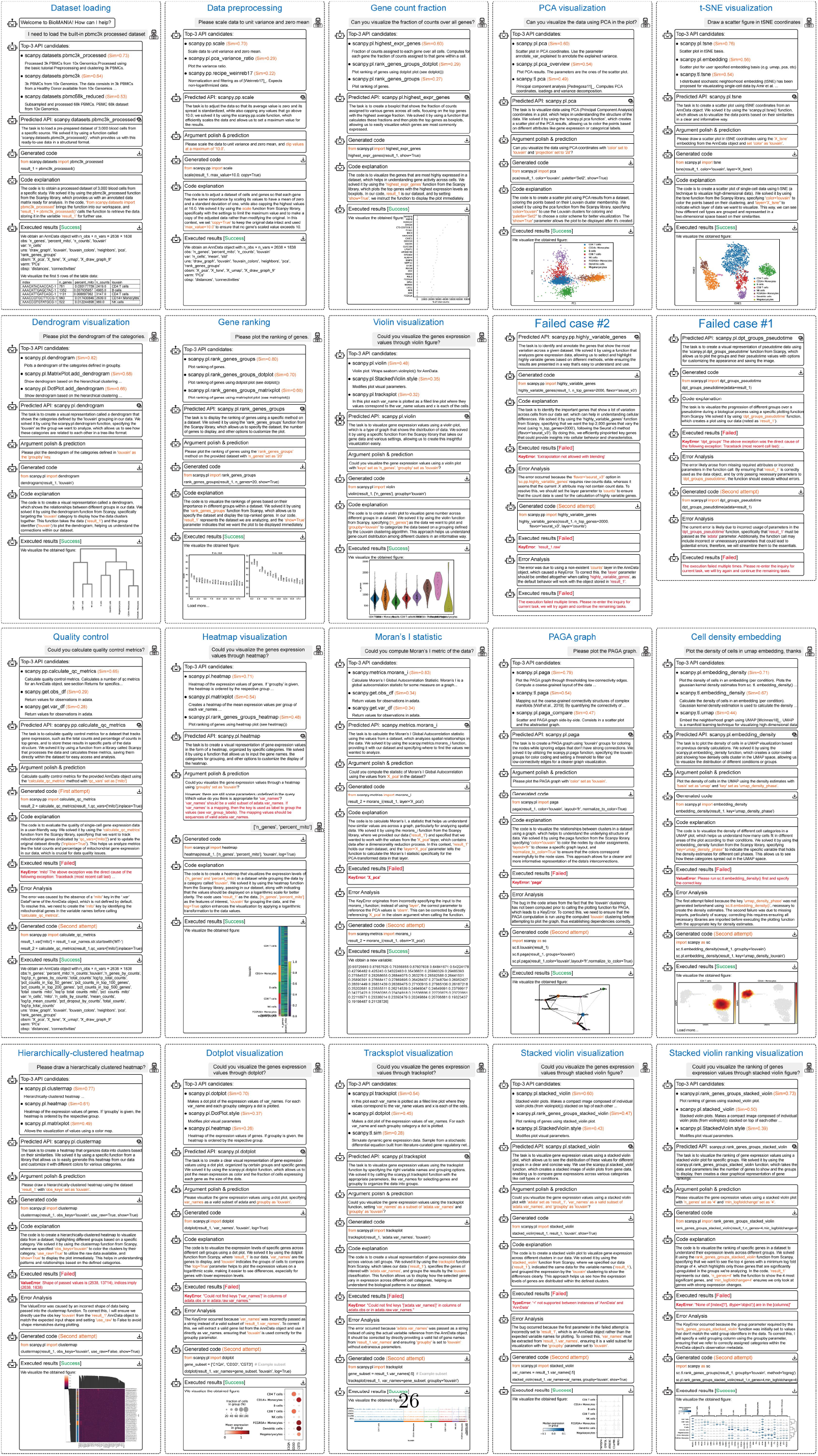
More examples of using BioMANIA for single-cell transcriptome data analysis tasks.

**Fig. A.8:**
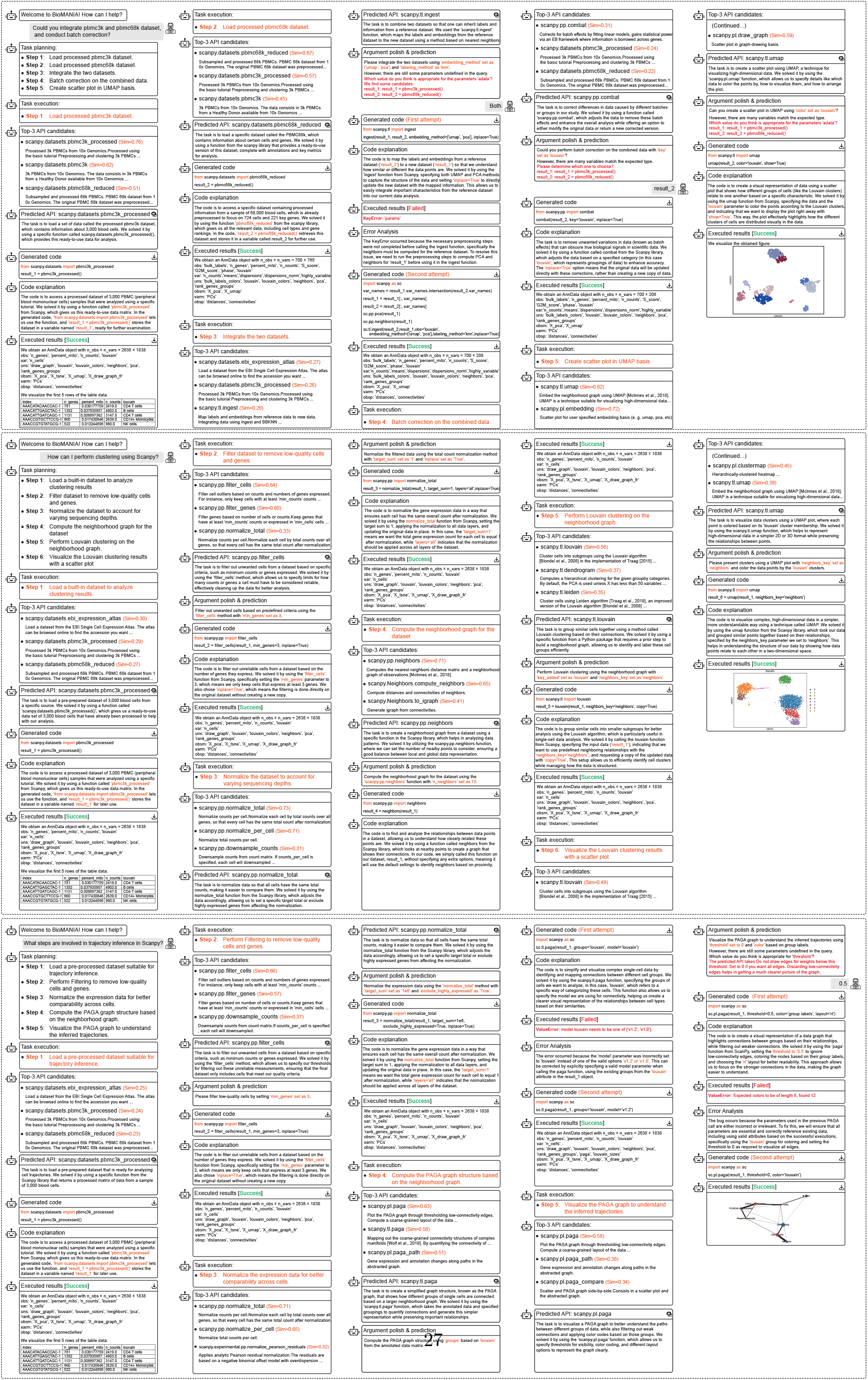
More examples of using BioMANIA for single-cell transcriptome data task planning.

**Fig. A.9:**
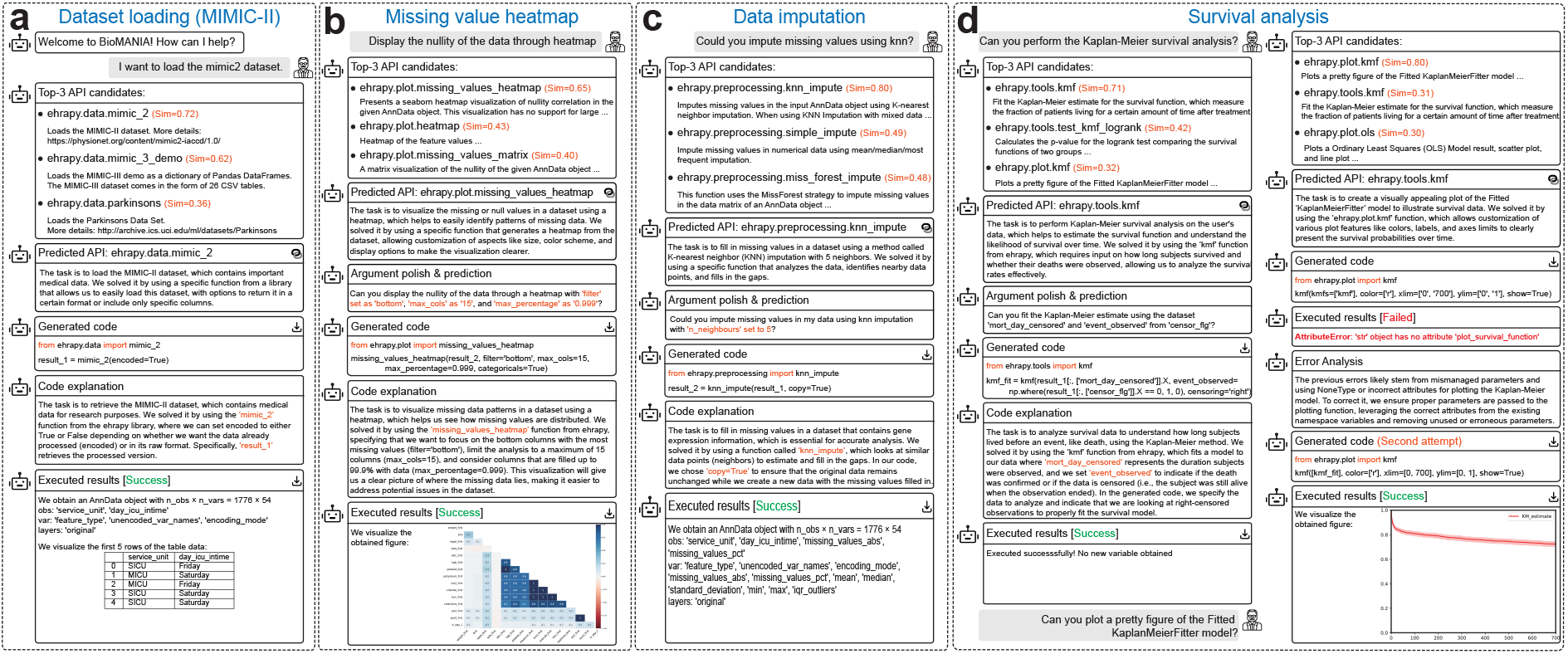
Illustrative examples of using BioMANIA for electronic health record data analysis. These examples encompass a wide range of electronic health record data analysis functionalities, including (**a**) Data loading, (**b**) Missing value visualization, (**c**) Missing value imputation, (**d**) Survival analysis.

### A.4 Failure cases of code generation using GPT-4

**GPT-4 Failure Example 1: Batch correction**

**Prompt:**

Provide the python code for batch correction with the API only from Scanpy. Apply it to the built-in pbmc3k processed dataset and pbmc68k reduced dataset.

**GPT-4 Turbo Answer:**

To perform batch correction on single-cell RNA sequencing data using the Scanpy library, we can employ different approaches, but one common method is to use the `Harmony` integration tool. `scanpy.external.pp.harmony integrate()` can be useful for batch correction by integrating datasets from different batches. Below is an illustrative Python code to perform batch correction using Scanpy on two datasets: `pbmc3k_processed` (which is assumed here after common preprocessing) and `pbmc68k reduced` datasets.

First, ensure you have the necessary libraries installed:

~~~
pip install scanpy anndata pandas numpy matplotlib scikit - learn seaborn
pip install harmonypy
~~~

Now, let’s script the batch correction using Harmony through Scanpy:

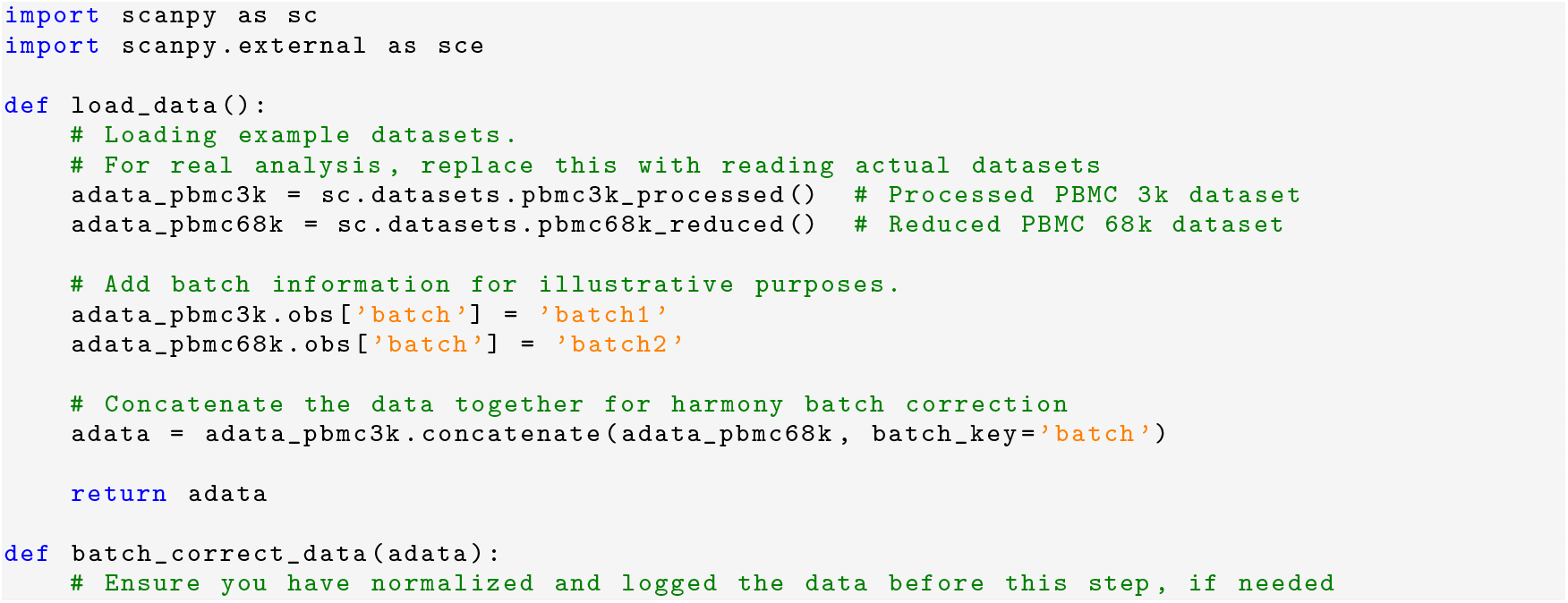
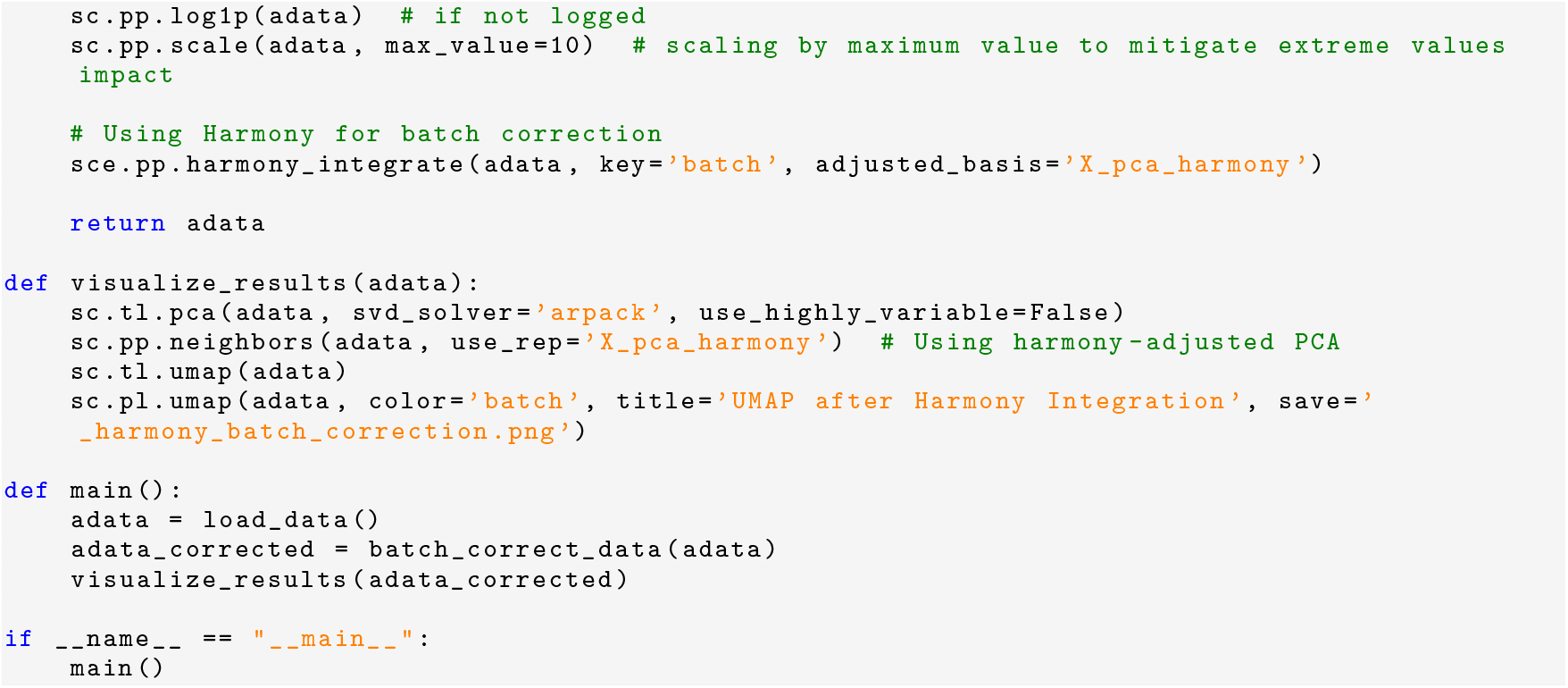

Explanation:

1. **Data Loading and Concatenation**: We load the `pbmc3k_processed` and `pbmc68k reduced` datasets. Here it’s presumed they are directly accessible via Scanpy’s dataset functions, which might not be the case outside theoretical or example contexts. Concatenate the datasets indicating that the concat will form distinct batches.
2. **Batch Correction**: We prepare the data by scaling and logging (if needed). Apply Harmony integration (from the scanpy external module) to correct for batch effects, using the batch labels for alignment.
3. **Visualization**: Post-correction visualizations using PCA and UMAP to display the effect of the batch correction.

The function `sce.pp.harmony integrate()` performs the actual integration (batch correction) work. This function operates assuming that the PCA has been performed on the dataset. Ensure any step relevant specifically to your data condition (like extra preprocessing) is included.

Remember to examine each step and adapt it to your specific datasets and conditions ensuring data compatibility and preprocessing. The provided code assumes direct use without specific compatibility adjustments.

**Note:**

The generated code raises the error `ValueError: Input X contains NaN.` in the codeline `sc.tl.pca(adata, svd_solver=‘arpack’)`. This is due to the reason that `adata` contains NaN values and needs to be preprocessed. GPT can refer to [this GitHub issue](https://github.com/scverse/scanpy/issues/2163) to debug.

**GPT-4 Failure Example 2: Trajectory inference**

**Prompt:**

Provide the python code for trajectory inference with the API only from Scanpy. Apply it to the built-in pbmc3k processed dataset.

**GPT-4 Turbo Answer:**

To perform trajectory inference on single-cell RNA sequencing data using Scanpy, you can utilize the PAGA (Partition-based Graph Abstraction) algorithm, which provides a way to infer paths through the data. The example below uses the `pbmc3k` dataset, which is a standard dataset for demonstration purposes in single-cell RNA sequencing analysis.

First, make sure you have the necessary packages installed. You’ll need `scanpy` primarily. You can install it using pip if you don’t have it already:

~~~
pip install scanpy
~~~

Below is the complete Python code, performing basic preprocessing, clustering, computing PAGA, and visualizing the trajectory based on Scanpy’s capabilities.

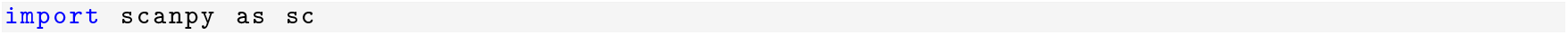
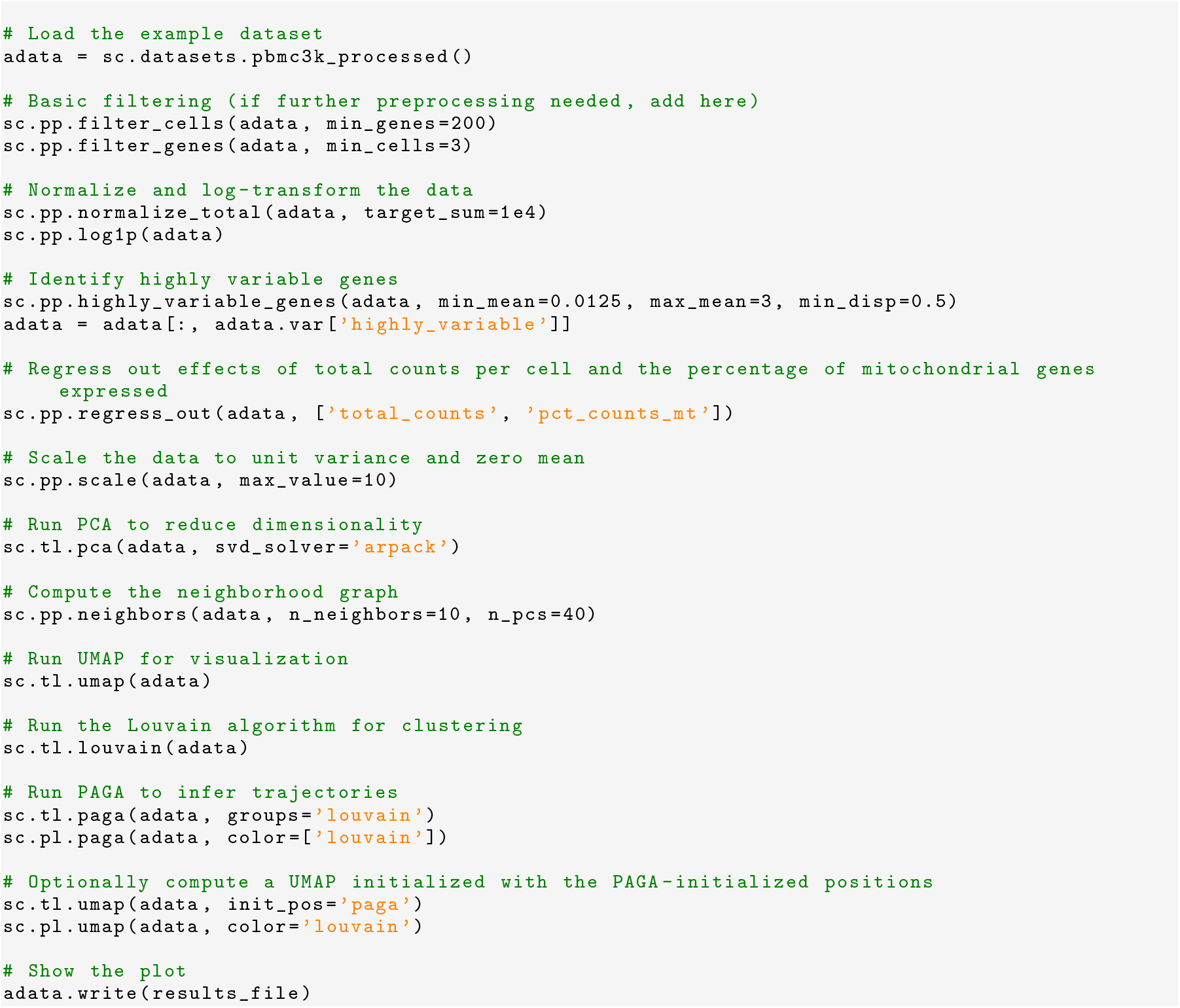

**In this code:**

1. Data is loaded using Scanpy’s built-in function (`sc.datasets.pbmc3k_processed()`).
2. Data is filtered by cells and genes to ensure robust analysis.
3. Normalization and log transformation are applied so that gene expression levels across cells can be more comparable.
4. Highly variable genes are identified to focus the analysis on genes that contribute most to variability among cells.
5. Linear regressions correct for confounding factors (total counts and mitochondrial gene expression).
6. PCA (Principal Component Analysis) is used for dimensionality reduction—a standard step in scRNA-seq analysis.
7. A neighborhood graph is computed to reflect the relationships (similarity) between cells.
8. The Louvain algorithm clusters the cells.
9. PAGA is used to infer and visualize the developmental trajectories among the discovered cell states (clusters).
10. Visualization is performed using UMAP, with PAGA paths overlayed to guide the initialization of UMAP coordinates, offering insightful perspectives on the data structure.

This approach provides a trajectory analysis of single-cell RNA data, inferring possible developmental pathways and relationships among cell types based on gene expression profiles. Adjustments might be necessary based on more specific needs or data characteristics.

**Note:**

API `scanpy.pp.highly_variable genes` will raise error `TypeError: putmask: first argument must be an array` for some dataset `pbmc3k_processed`, while other dataset `pbmc3k` will not. This is also an open issue submitted by other users.

**GPT-4 Failure Example 3: Gene ranking**

**Prompt:**

Provide the python code for plotting the ranking of genes with the API only from Scanpy. Apply it to the built-in pbmc3k processed dataset.

**GPT-4 Turbo Answer:**

Below is a Python example that uses Scanpy to process the `pbmc3k` dataset and then plots the gene rankings. This example first preprocesses the data (for normalization and selection of highly variable genes) and then ranks the genes based on their scores using PCA as part of the analysis. Finally, the gene rankings are plotted using `sc.pl.rank_genes_groups`. Here are the steps outlined:

1. Load the necessary libraries.
2. Load the example dataset.
3. Preprocess the data (normalization, logarithmic transformation, identifying highly variable genes).
4. Perform PCA to find the principal components.
5. Cluster the cells.
6. Find marker genes for each cluster.
7. Plot the ranking of genes.

Ensure that Scanpy and its dependencies are properly installed in your Python environment. You can install Scanpy using pip:

~~~
pip install scanpy
~~~

Here’s the code:

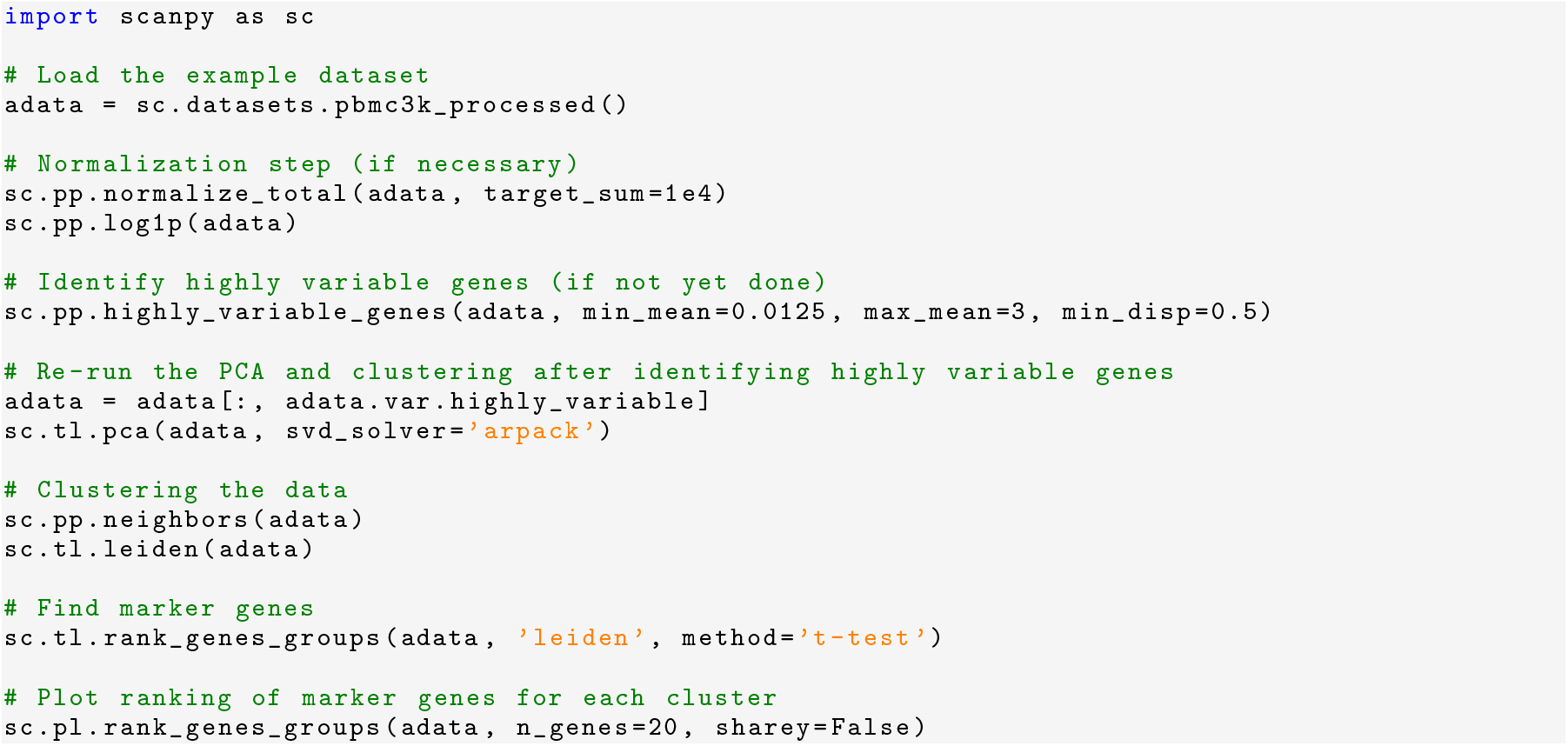

**Notes:**

- The `sc.datasets.pbmc3k_processed()` method fetches a dataset that has been preprocessed already, which includes normalization and the identification of highly variable genes. Some steps above are repeated for educational purposes and might not affect the pre-processed dataset.
- Adjust the clustering, PCA, and marker gene detection based on the specific analysis needs and the characteristics of the dataset.
- The number of genes (`n genes`) shown in the plot can be adjusted.

This code will provide a visual representation of the top differentially expressed genes across the identified clusters, which can help in understanding the biological heterogeneity of the cell population. Adjust parameters and steps based on your specific analysis requirements and to dive deeper into specific aspects of the dataset.

**Note:**

When the targeted task requires more pre-request steps, it has a higher probability of generating errors. It meets the same error `TypeError: putmask: first argument must be an array` by `scanpy.pp.highly_variable genes`.

**GPT-4 Failure Example 4: Violin visualization**

**Prompt:**

Provide the python code for visualizing the genes expression values through violin figure with the API only from Scanpy. Apply it to the built-in pbmc3k processed dataset.

**GPT-4 Turbo Answer:**

To visualize gene expression values using a violin plot in Scanpy, you’ll need to import the library and load a standard processed dataset. Below is detailed Python code that uses Scanpy to visualize gene expression in a violin plot from the built-in `pbmc3k` dataset.

First, ensure you have installed Scanpy and other necessary libraries, like pandas and numpy. If you haven’t installed these packages yet, you can install them via pip:

~~~
pip install scanpy pandas numpy matplotlib
~~~

Now, here’s a step-by-step Python script to visualize gene expression using a violin plot:

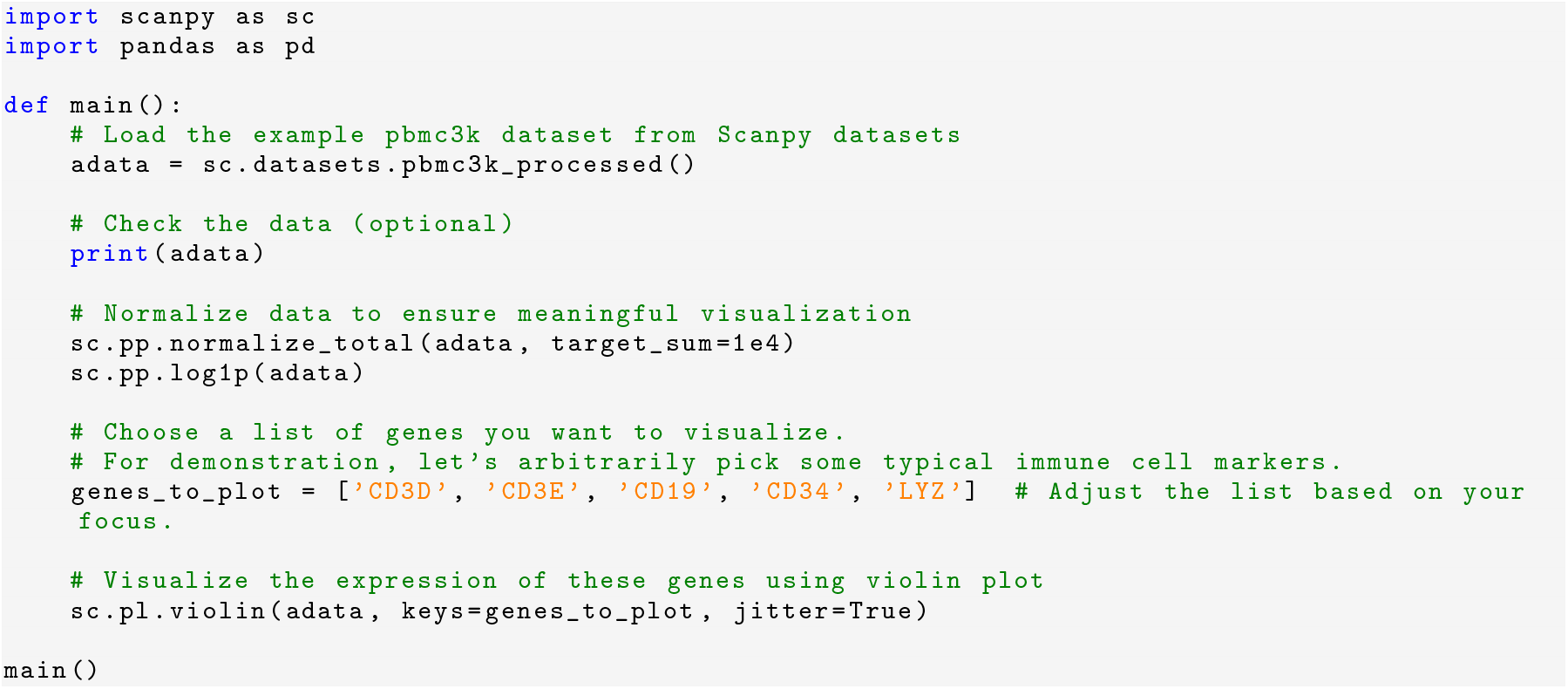

**Description of Steps:**

1. **Import necessary libraries**: Import Scanpy as well as other dependencies.
2. **Loading the Data**: The `sc.datasets.pbmc3k_processed()` function fetches the preprocessed dataset having 3k PBMCs from 10X Genomics.
3. **Normalize the data**: Before visualization, normalization is performed to scale the data while keeping the relative differences in gene expression values intact.
4. **Gene Selection**: Define genes you are interested in. Replace `‘CD3D’, ‘CD3E’, ‘CD19’, ‘CD34’, ‘LYZ’` with other genes of interest as required.
5. **Violin Plot Visualization**: Use `sc.pl.violin()` from Scanpy to create violin plots of these genes to observe their expression profiles in the dataset.
6. **Jitter**: Enabled jitter to better visualize individual data points.

**Considerations:**

- **Gene Names**: Ensure that gene names used in the plots are correct and they exist in the dataset. You can check available genes via `adata.var_names`.
- **Additional Adjustments**: Depending on your specific requirements or dataset, you might need additional preprocessing steps, or you might want to adjust the layout or aesthetics of the plot.

This code provides a basic outline and can be modified based on further specifications of the analysis or visual requirements.

**Note:**

The generated code raises the error `KeyError: “Could not find keys ‘[‘CD34’]’ in columns of ‘adata.obs’ or in ‘adata.raw.var_names.’”`. This occurs because it didn’t know which attribute indeed exists in the `adata` object.

**GPT-4 Failure Example 5: Quality control**

**Prompt:**

Provide the python code for calculating quality control metrics with the API only from Scanpy. Apply it to the built-in pbmc3k dataset.

**GPT-4 Turbo Answer:**

To calculate quality control metrics for single-cell RNA sequencing data using Scanpy, and applying it to the built-in `pbmc3k` dataset, you can follow these steps in Python. First, ensure you have Scanpy installed. If not, you can install it using pip:

~~~
pip install scanpy
~~~

Next, you can load the PBMC 3k dataset directly from Scanpy, calculate the quality control metrics, plot them, and filter_cells based on these metrics. Here is how you can do it:

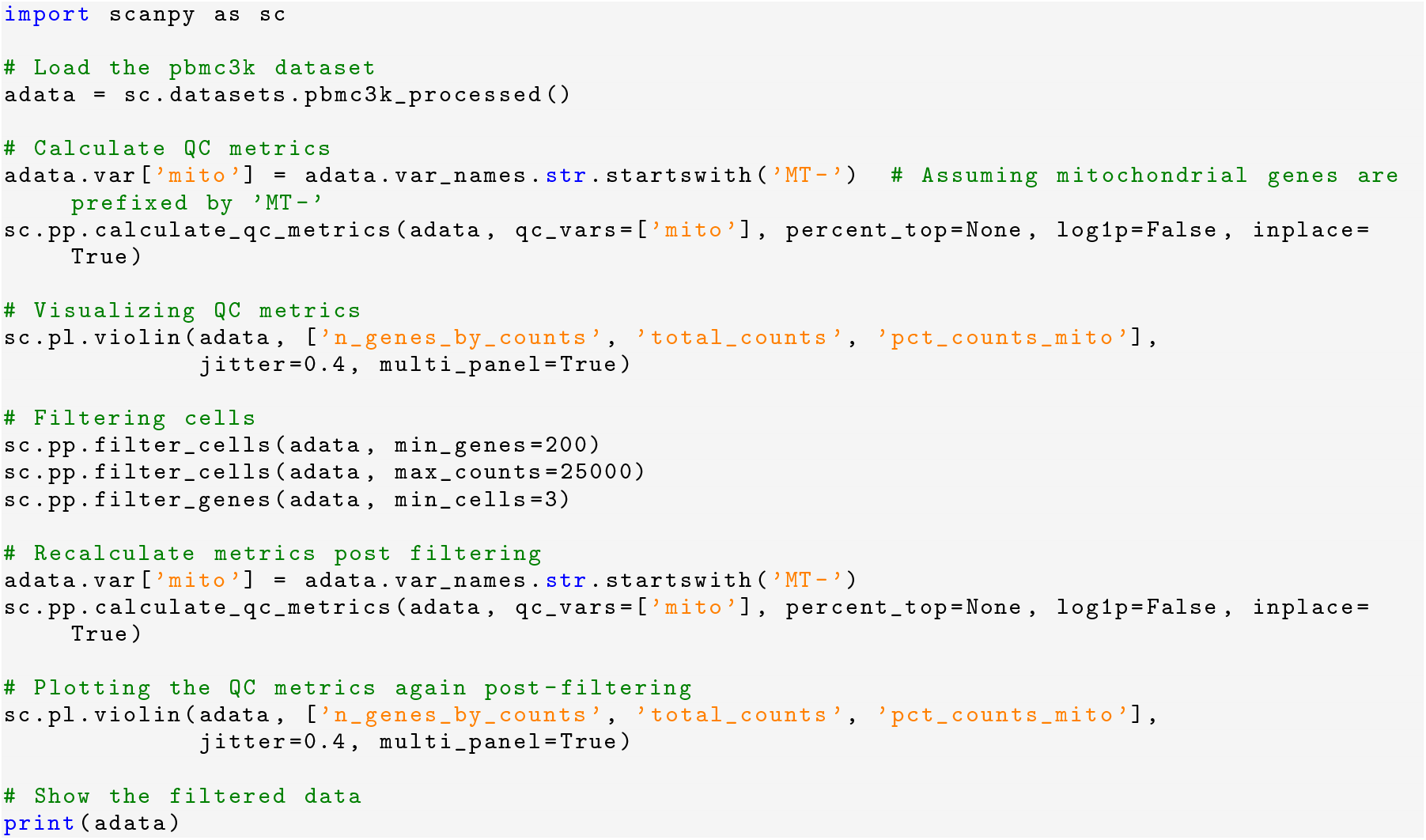

**Explanation:**

1. **Loading the Data**: `sc.datasets.pbmc3k_processed()` loads a preprocessed dataset of PBMC (Peripheral Blood Mononuclear Cells) obtained from 10X Genomics. The dataset is already normalized and log-transformed.
2. **Calculate QC Metrics**: We’re adding a column to locate mitochondrial genes (commonly used as a quality control metric in single-cell data). Using `scanpy.pp.calculate_qc_metrics`, we compute the number of genes expressed, total counts, and percentage of mitochondrial genes for each cell.
3. **Visualizing QC Metrics**: Using `scanpy.pl.violin`, we plot these metrics to visualize potential outliers and get a sense of the overall data quality.
4. **Filtering Data**: We filter out cells with fewer than 200 detected genes or more than 25,000 total counts to remove low-quality cells. This step may need adjustments based on specific datasets or research requirements.
5. **Recalculation and Revisualization Post Filtering**: After filtering the dataset, it’s a good practice to recalculate and plot QC metrics again to confirm the data’s quality has improved and check the effect of the filtering.

The steps above provide a basic framework for applying QC metrics using Scanpy to a single-cell RNA sequencing dataset. Adjustments may be needed depending on specific data characteristics or research requirements.

**Note:**

The generated code didn’t raise any errors, but the preprocessing steps such as `scanpy.pp.filter_cells(adata, max_counts=25000)` show that nothing is filtered. The parameters predicted by GPT might not be reasonable enough to filter out the data.

**GPT-4 Failure Example 6: Heatmap visualization**

**Prompt:**

Provide the python code for visualizing the genes expression values through heatmap with the API only from Scanpy. Apply it to the built-in pbmc3k processed dataset.

**GPT-4 Turbo Answer:**

To visualize gene expression values as a heatmap in Python using the Scanpy API, first, you’ll need to install the Scanpy library if it’s not already installed in your Python environment. Scanpy is a popular library for analyzing single-cell RNA sequencing data, which includes functions to preprocess data, perform clustering, and visualize results including generating heatmaps.

Here is how you could do it, step by step, including loading a built-in dataset (such as the `pbmc3k_processed` dataset), selecting genes, and plotting a heatmap.

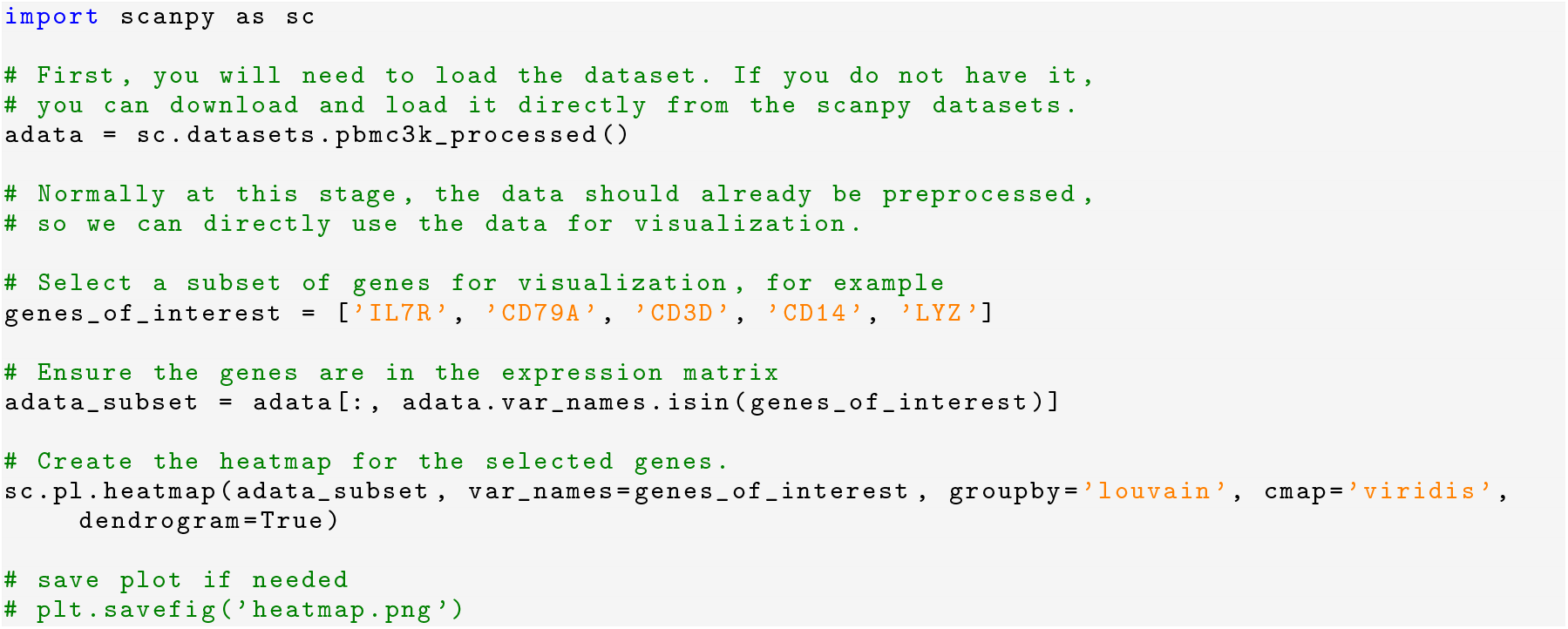

Here’s the detailed explanation of what the code does:

1. We import the necessary library, `scanpy`.
2. We load the `pbmc3k_processed` dataset. This dataset is often used as a benchmark in single-cell RNA-seq analyses and comes preprocessed (i.e., normalized, log-transformed, etc.).
3. We specify a list of gene names that we are particularly interested in. This must be based on the biological question you wish to explore.
4. We use a subset of the `adata` object that only includes the selected genes to make the heatmap more informative and focused.
5. We create a heatmap using `scanpy.pl.heatmap`. The `groupby` parameter is set to ‘louvain’, which refers to the Louvain community detection method used in the clustering step of processing this dataset. `cmap` ‘viridis’ can be replaced with other colormap names depending on preference.
6. Optional: You can save the plot as an image file using matplotlib’s `savefig()`.

This should give you a clear heatmap of how the selected genes are expressed across the computed Louvain clusters in the provided single-cell dataset.

**Note:**

The generated code raises the error `ValueError: Distance matrix ‘X’ must be symmetric`. This code works without the preprocessing step `adata_subset = adata[:, adata.var_names.isin(genes_of_interest)]`, which means GPT doesn’t know the constraint on the input of the API `scanpy.pl.heatmap`. Also, the preprocessing step of GPT generated code is not reasonable as it filters out almost all cells while only remaining one cell `CD79A`.

**GPT-4 Failure Example 7: Moran’s I Metric**

**Prompt:**

Provide the python code for computing Moran’s I metric of the data with the API only from Scanpy. Apply it to the built-in pbmc3k processed dataset.

**GPT-4 Turbo Answer:**

To compute Moran’s I metric using Python, specifically within the context of single-cell RNA sequencing data, we will use the Scanpy library. Moran’s I is a measure of spatial autocorrelation which, in the context of single-cell data, can provide insights into the spatial arrangements or patterns of gene expression.

First, if you haven’t already installed Scanpy, you can install it using pip:

~~~
pip install scanpy
~~~

Below is a step-by-step Python code on how to load the `pbmc3k` dataset, preprocess it, compute Moran’s I for a gene, and visualize it. Note that Moran’s I itself is not directly included in Scanpy’s standard functions but can be computed using the `scanpy.metrics.morans_i` function, provided you have an adequate spatial dataset. For demonstration, we’ll treat the `pbmc3k` dataset as if it has spatial data by creating some mock spatial data.

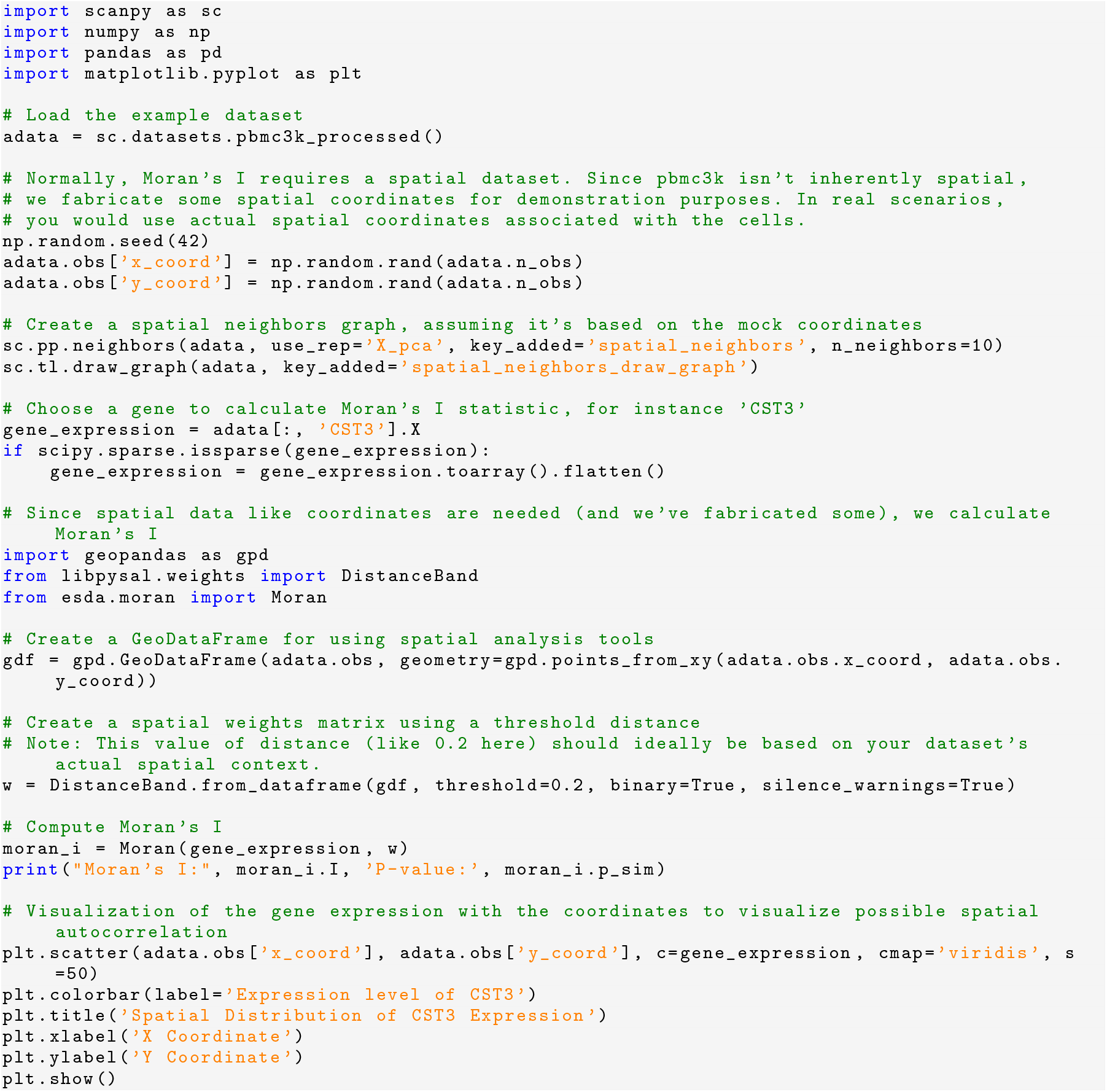

In this example, the Moran’s I statistic would be calculated using mocked spatial data. Note that genuine spatial data (from experiments like spatial transcriptomics) should be used for meaningful biological insights. The visualization provides a mock view on how gene expression patterns might look in a spatial data context. Adjust the threshold and methods as per your actual spatial data and experiment needs.

**Note:**

The generated code has two issues. The first issue is that it assumes `key_added` is the parameter of `scanpy.tl.draw graph`, while it is not. As a result, it raises the error `TypeError: ‘key_added’ is an invalid keyword argument for this function`. Another issue is that it uses the `Moran` function from the external library `esda.moran` instead of calling its API `scanpy.metrics.morans_i`. This API exists in the 1.8.x version of Scanpy, which was [released](https://pypi.org/project/scanpy/#history) before April 1, 2022. So this API should be recognized by GPT.

**GPT-4 Failure Example 8: Cell density embedding**

**Prompt:**

Provide the python code for plotting the density of cells in UMAP embedding with the API only from Scanpy. Apply it to the built-in pbmc3k processed dataset.

**GPT-4 Turbo Answer:**

To plot the density of cells in UMAP embedding using the Scanpy library and applying it to the built-in `pbmc3k` dataset, you first need to install Scanpy if you haven’t done so yet. You can install it via pip:

~~~
pip install scanpy
~~~

Now, here’s the Python code that will load the `pbmc3k` dataset, perform the necessary preprocessing, compute the UMAP embedding, and plot the density of cells on the UMAP plots.

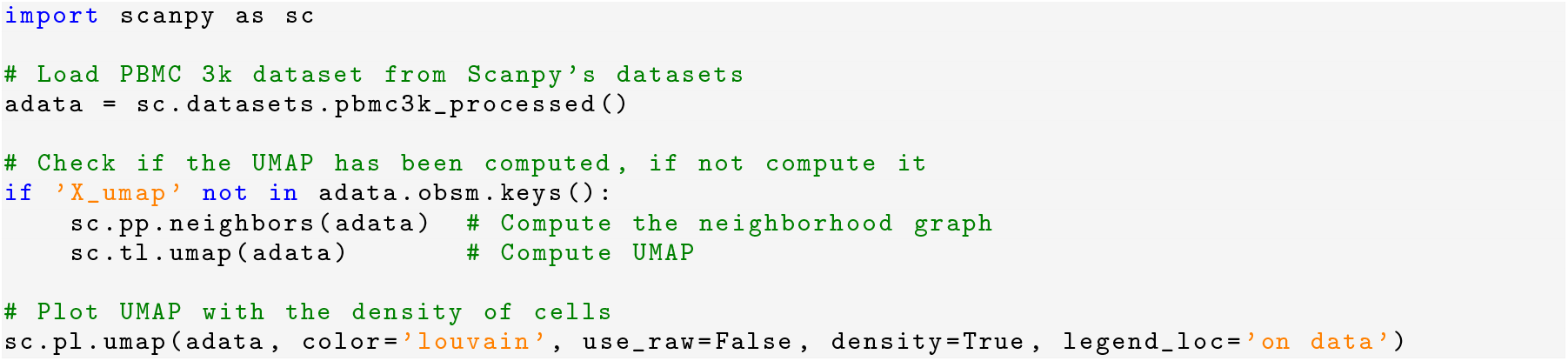

**Explanation of Code:**

1. The dataset `pbmc3k_processed` is preprocessed, meaning it includes PCA results and some clustering (usually stored in `adata.obs`).
2. Before plotting, it checks if UMAP embeddings (`X_umap`) are already in the AnnData object (`adata`). If they are not, it computes a neighborhood graph and then UMAP embeddings.
3. The `sc.pl.umap()` function from Scanpy is used with `density=True` to plot cell densities, `color=‘louvain’` to color cells by their Louvain cluster, and `use_raw=False` to indicate that the UMAP should be plotted using processed data.

This will create a UMAP visualization color-coded by Louvain clusters, overlayed with a density plot that highlights areas with higher cell concentrations. Ensure that `legend_loc=‘on data’` is set to add cluster identifiers directly on the plot for better visual cues of cluster locations in high-density regions.

**Note:**

The generated code raises an error `AttributeError: PathCollection.set() got an unexpected keyword argument ‘density’`, which indicates that the API `scanpy.pl.umap` should not treat `density` as its parameter.

**GPT-4 Failure Example 9: Hierarchically-clustered heatmap**

**Prompt:**

Provide the python code for drawing a hierarchically clustered heatmap with the API only from Scanpy. Apply it to the built-in pbmc3k processed dataset.

**GPT-4 Turbo Answer:**

To draw a hierarchically clustered heatmap using the Scanpy library, you can follow these steps involving loading the built-in `pbmc3k` dataset, performing necessary preprocessing, and finally plotting the heatmap. Below is the complete Python code to achieve this.

1. Import necessary libraries.
2. Load the dataset.
3. Preprocess the data (normalization, log transformation, detecting highly variable genes).
4. Perform PCA for dimensionality reduction.
5. Compute the neighborhood graph.
6. Cluster the data.
7. Identify marker genes.
8. Plot heatmap with hierarchical clustering.

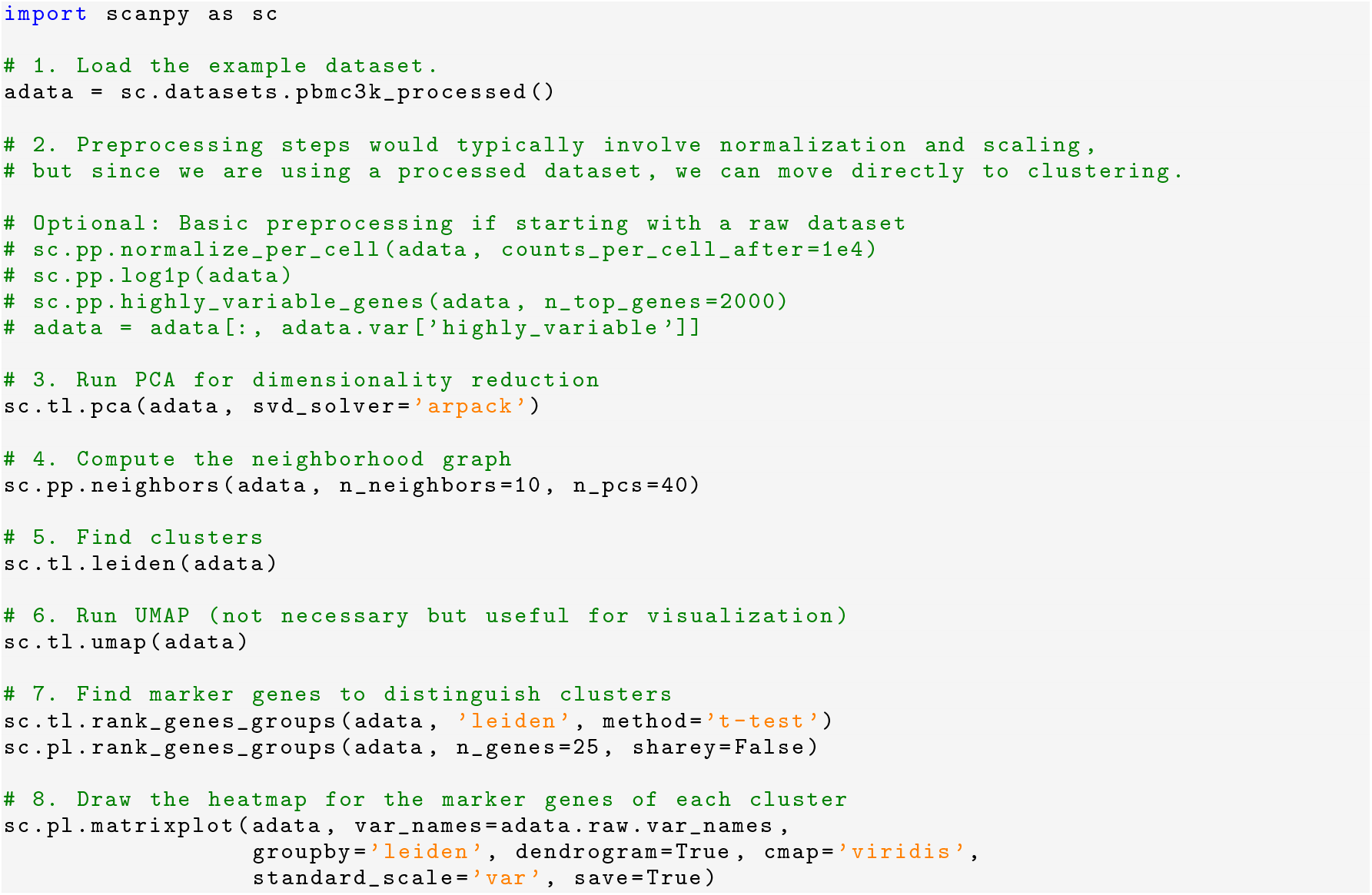

**Explanation of Key Steps:**

1. **Load Data**: The `pbmc3k_processed` dataset provided by Scanpy is used.
2. **Preprocessing**: This typically involves normalization, logarithmic transformation, and selection of highly variable genes. However, since the dataset is already processed, these steps are optional.
3. **PCA**: Principal component analysis reduces the dimensionality of the data, retaining essential features.
4. **Neighbors Computation**: To understand the local structure of the data, a neighborhood graph is computed.
5. **Clustering**: The Leiden algorithm identifies relatively densely connected groups of cells.
6. **Marker Genes**: Identifying marker genes helps in understanding the defining characteristics of each cluster.
7. **UMAP**: A visualization tool to represent the data in a low-dimensional space. This is useful for examining how clusters arrange spatially.
8. **MatrixPlot**: This function creates a heatmap of gene expression, with options for hierarchical clustering of rows (genes) and columns (samples/groups).

This pipeline will result in a heatmap where clusters of cells can be observed based on their expression profiles and hierarchical groupings both of samples and genes. The `dendrogram=True` in the `matrixplot` function enables hierarchical clustering based on the data. Adjust the `var_names` parameter to focus on specific genes or use the results from `rank_genes_groups` to explore markers for each cluster. Ensure you have an adequate version of Scanpy installed that supports all function calls (usually Scanpy *≥* 1.4).

**Note:**

There are three issues with the GPT-generated code. One is that it didn’t use `scanpy.pl.clustermap`, which was already developed in Scanpy 1.8.x version. Another issue is that it raises the error `ValueError: Image size of 507648×380 pixels is too large. It must be less than 2^16 in each direction.`, which suggests that the data needs preprocessing (like filtering). The final issue is its optional preprocessing code, which is commented out. This code raises a `TypeError: putmask: first argument must be an array` when calling `scanpy.pp.highly_variable genes(adata, n top genes=2000)`.

**GPT-4 Failure Example 10: Dotplot visualization**

**Prompt:**

Provide the python code for visualizing the genes expression values through dotplot with the API only from Scanpy. Apply it to the built-in pbmc3k processed dataset.

**GPT-4 Turbo Answer:**

To visualize gene expression data using a dotplot from the Scanpy API, and especially applying this visualization on the built-in `pbmc3k_processed` dataset, you should follow these steps. The steps include loading the necessary data, preprocessing if needed, and finally plotting the data using the `sc.pl.dotplot` function from Scanpy. Below is a detailed Python script example that demonstrates how to achieve this:

1. **Import necessary libraries**
2. **Load the dataset**
3. **Preprocess data** (if required, for example, normalize or identify highly variable genes)
4. **Plot the data using a dotplot**

First, ensure you have the `scanpy` package installed. You can install it using pip if it’s not installed:

~~~
pip install scanpysss
~~~

Here is a comprehensive Python script:

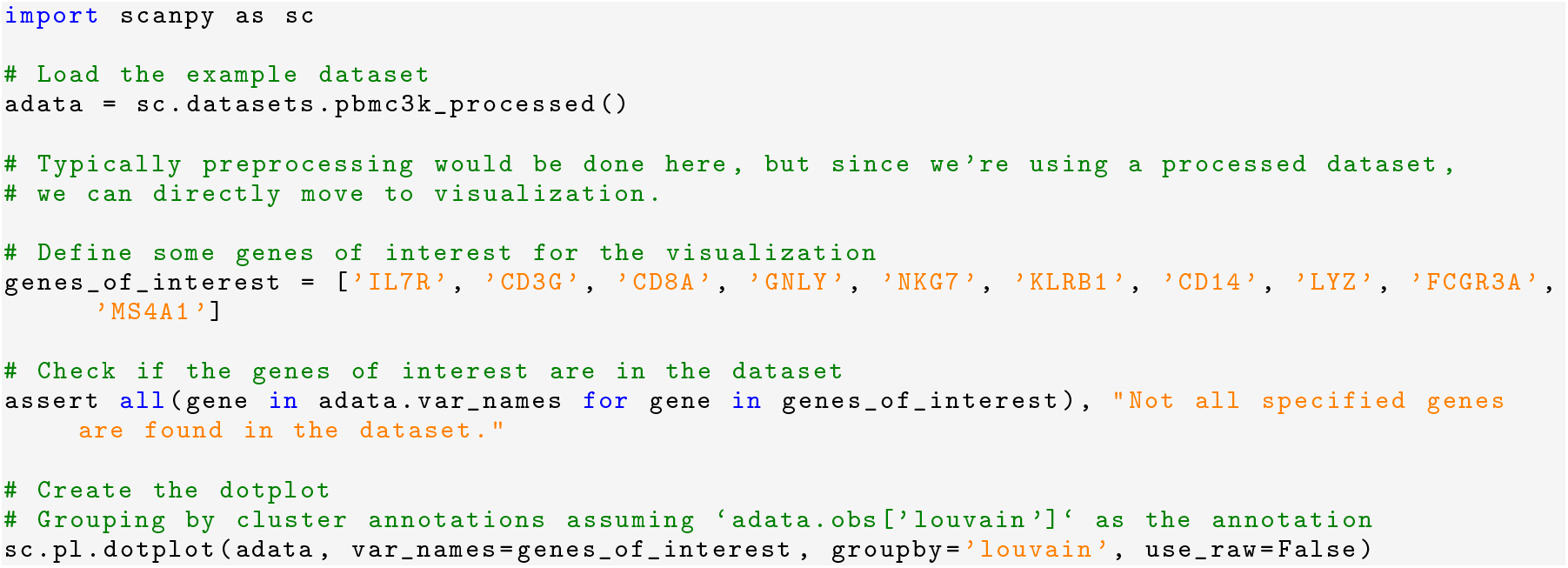

**Explanation:**

1. **Import Scanpy**: This is needed for processing and visualization.
2. **Load Dataset**: `sc.datasets.pbmc3k_processed()` loads the processed PBMC dataset from 10x Genomics. This dataset is handy for demonstrations and comes pre-loaded with cluster annotations, gene filters, and some normalization.
3. **Define Genes of Interest**: These are the genes you want to focus on in your dotplot. Modify this list based on your research focus.
4. **Create Dotplot**: Using the `sc.pl.dotplot` function, the data is visualized. The genes specified are plotted against clusters identified (`groupby=‘louvain’`). `use_raw=False` uses the processed data as opposed to raw count data.

Just run the code to visualize the expression of the chosen genes across different cell clusters identified in the dataset. The assumptions include the dataset having Louvain cluster assignments already performed, and genes listed are already filtered and relevant in this context.

Remember, for a different dataset or different stages of preprocessing, you might need to adjust the steps, including normalizing the data or re-running clustering. This script assumes that such steps are already completed in the `pbmc3k_processed` dataset.

**Note:**

The error `AssertionError: Not all specified genes are found in the dataset.` occurs because GPT writes an assertion that the given gene list exists in the built-in dataset. It will raise error as GPT doesn’t know exactly which genes exist in the provided dataset.

**GPT-4 Failure Example 11: Tracksplot Visualization**

**Prompt:**

Provide the python code for visualizing the genes expression values through tracksplot with the API only from Scanpy. Apply it to the built-in pbmc3k processed dataset.

**GPT-4 Turbo Answer:**

To create a genes expression visualization using a tracks plot for the `pbmc3k` dataset using the Scanpy API, you’ll first need to load the dataset, preprocess it as necessary, and then use the `sc.pl.tracksplot` function to generate the plot. Below I’ll guide you through the complete process, including necessary preprocessing steps like normalization, principal component analysis, and computing the neighborhood graph, which are typical for single-cell RNA-seq data analysis.

First, you need to ensure that you have Scanpy installed in your Python environment. If it’s not installed, you can install it using pip:

~~~
pip install scanpy
~~~

Now, here’s a complete example of how to handle the `pbmc3k` dataset and visualize specific gene expressions with tracksplot:

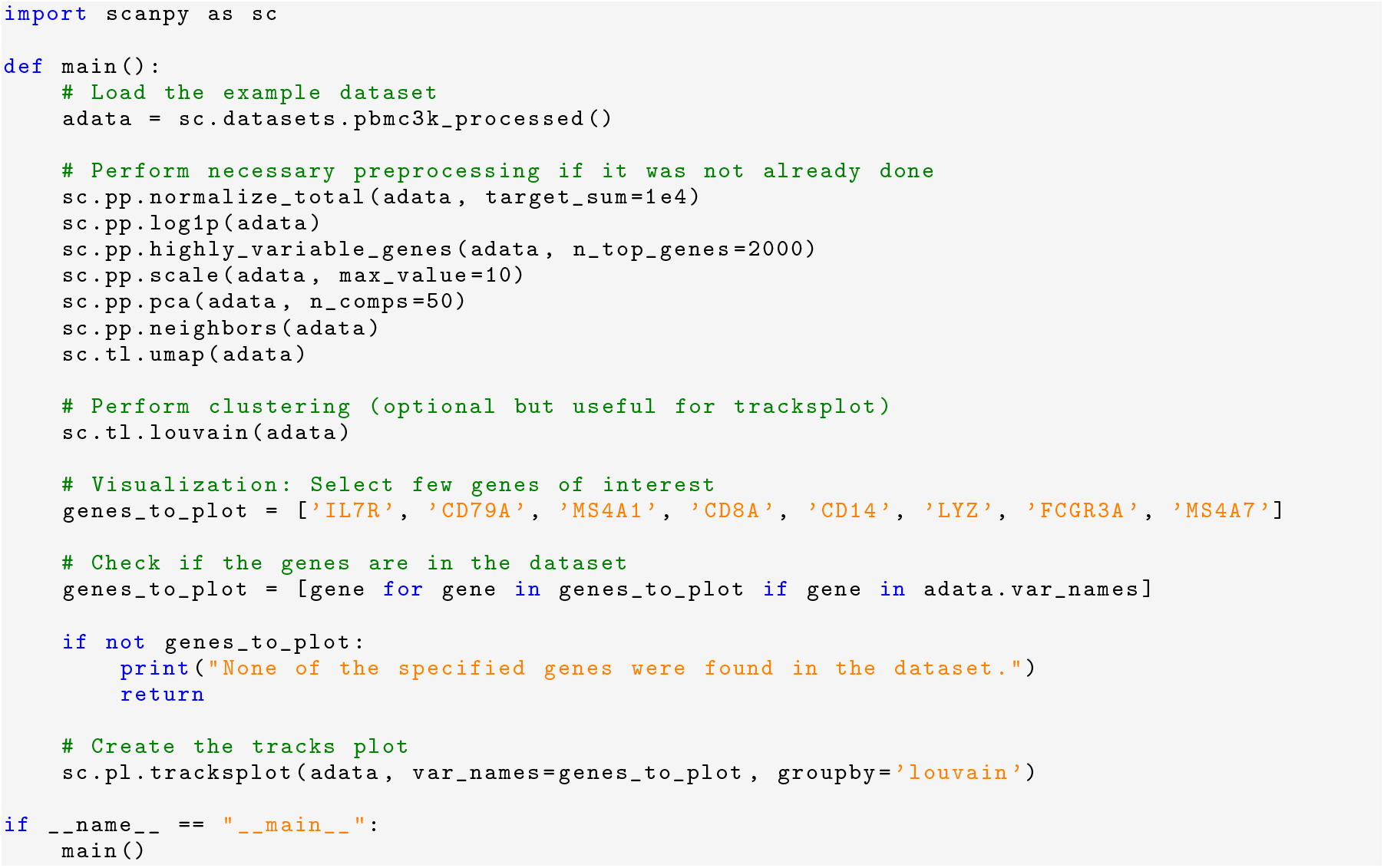

**Explanation:**

1. **Data Loading**: We load the processed version of the `pbmc3k` dataset directly from Scanpy’s dataset collection. This dataset has been pre-processed, including normalization and log-transformation.
2. **Check and Preprocess (if necessary)**: Though `sc.datasets.pbmc3k_processed()` generally contains a preprocessed dataset, the code snippet includes checks and preprocessing steps to ensure all necessary preprocessing has been done. This includes normalization, logarithmic transformation, selection of highly variable genes, scaling, PCA, computation of neighborhood graphs, UMAP for dimension reduction, and clustering using the Louvain algorithm.
3. **Gene Selection**: You can customize `genes_to_plot` with the genes you are interested in visualizing.
4. **Tracks Plot**: The `sc.pl.tracksplot` function generates a tracks plot. Here, it is grouped by the Louvain clusters which categorizes the cells into different clusters.

Make sure that the gene names specified in `genes_to_plot` are correctly named and present in the dataset. You can adjust the list of genes based on your specific genes of interest for the PBMC dataset. This code will run in any standard Python environment where Scanpy is installed.

**Note:**

This generated code contains two issues. The first issue arises with `scanpy.pp.highly_variable genes`, leading to the error `TypeError: putmask: first argument must be an array`. The second issue is related to `scanpy.pp.pca`, which raises the error `ValueError: Input X contains NaN`.

**GPT-4 Failure Example 12: Stacked violin ranking visualization**

**Prompt:**

Provide the python code for visualizing the ranking of genes expression values through stacked violin figure with the API only from Scanpy. Apply it to the built-in pbmc3k processed dataset.

**GPT-4 Turbo Answer:**

Below, I provide a Python code snippet to visualize the ranking of gene expression values using a stacked violin plot utilizing the Scanpy API. This snippet applies to the built-in `pbmc3k_processed` dataset in Scanpy. Before running this script, make sure you have installed the necessary packages (`scanpy`, `anndata`, and related dependencies) in your Python environment.

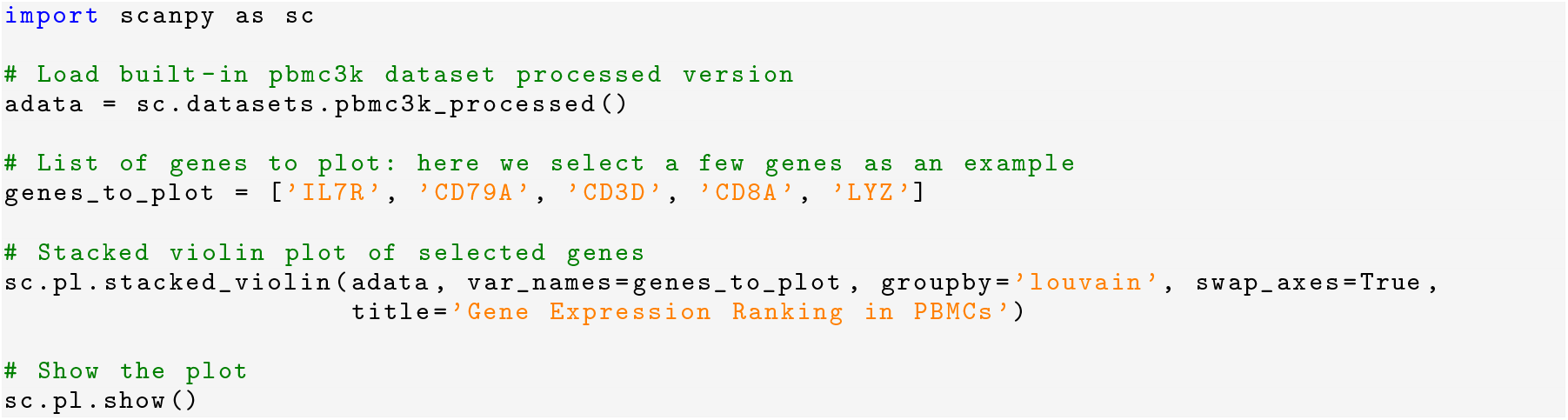

**Explanation:**

1. **Importing Scanpy**: We first import Scanpy, which is a widely-used library for single-cell analysis.
2. **Loading Data**: We load the `pbmc3k_processed` dataset directly from Scanpy’s datasets. This dataset is preprocessed, including normalization and clustering, representing a typical workflow result.
3. **Gene Selection**: Here, a list `genes_to_plot` is defined containing the gene names for which the expression is to be visualized. You may modify this list based on the genes of interest or focus on top expressed genes identified via previous analysis.
4. **Stacked Violin Plot**: The `sc.pl.stacked violin` function is used to create the plot. Important parameters include:

- `var_names`: Names of the variables or genes.
- `groupby`: The category over which to group the data, here the `louvain` clustering results.
- `swap_axes`: With this set to True, genes become rows and categories become columns, which might fit better for presentations depending on the plot’s orientation preference.
- `title`: Setting up the title of the plot.

5. **Display the Plot**: `sc.pl.show()` is used to render the plot, although this is optional in most environments as plots are displayed automatically by default.

Make sure that the gene names used in `genes_to_plot` exist within the dataset. You can check the genes available in your dataset using `adata.var_names` to customize the `genes_to_plot` list further.

**Note:**

The generated code has already finished the task. However, it treats `scanpy.pl.stacked violin` as a rendering function and mistakenly tries to use `scanpy.pl.show()` to display the figure. But `scanpy.pl.show()` is not a valid API from Scanpy. The figure will be displayed automatically once the `sc.pl.stacked violin` function is executed.

### A.5 Prompts used by BioMANIA

#### A.5.1 Prompt for task planning

\# **Task Description**

\## **Common part**

Create step-by-step task plan with subtasks to achieve the goal.

Tone: Vary the tone appropriately across tasks, such as polite, straightforward, or casual.

Subtask length: Each subtask should consist of around 10 words.

Subtask scope: Each subtask should invoke only one API from the {lib} PyPI library. Split subtasks if they contain more than one action. For example, use “Please filter the data” alongside “Could you normalize the data?” instead of “Please filter and normalize.” Use “Please load/filter/normalize dataset1” and “Can you load/filter/normalize dataset2” instead of “How to load/filter/normalize dataset1 and dataset2.”. Use “Integrate two datasets” when integration requires two datasets.

Subtask order: Ensure prerequisites appear before subsequent API calls.

Number of subtasks: Create approximately 5 tasks based on complexity. The visualization step should appear last and not be included in previous steps.

Focus on Operations: Concentrate on actions and objectives without over-explaining the object being operated on. For instance, “Please create scatter plot in UMAP basis” emphasizes the action rather than the object (data). Use “Could you process the data” instead of “Could you process the data for spatial analysis.”

Avoid Dataset Mentions: Aside from the data-loading step, avoid directly mentioning the dataset. For example, use “Could you draw scatter plot for the data” instead of “Could you draw scatter plot for the pbmc3k data?”. Notice that never mention keywords like ‘pbmc3k’, ‘pbmc68k’ except for the data loading step.

Goal Oriented Structure: State the goal at the beginning of each subtask. Be clear and concise, omit API names, and focus on key actions only.

Only respond in JSON format strictly enclosed in double quotes, adhering to the Response Format. Exclude any extraneous content from your response.

Examples:

Goal: use Squidpy for spatial analysis of Imaging Mass Cytometry data, focusing on the spatial distribution and interactions of cell types

Response:

~~~
{
“plan”: [
“step 1: Load pre-processed Imaging Mass Cytometry data.”,
“step 2: Could you show cell type clusters in spatial context?”,
“step 3: Please calculate co-occurrence of cell types across spatial dimensions.”,
“step 4: Compute neighborhood enrichment.”,
“step 5: Can you plot neighborhood enrichment results?”
]
}
~~~

Now finish the goal with the following information:

Goal: {goal_description}

Response Format: `{“plan”: [“step 1: content”, “step 2: content”, …]}`

\## **With_or_without_data**

\### **With_data**

{Common_part}. Input: Your data is provided in the format ‘file path: file description’ from {data_list}

\### **Without_data**

{Common_part }. You have no local data. Please use only APIs that access built-in datasets and avoid those requiring local data loading.

\# **Example_Input**:

Lib: Scanpy

Goal_description: Could you integrate pbmc3k processed and pbmc68k processed dataset, and conduct batch correction?

\# **Example Output**:

~~~
{
“plan”: [
“step 1: Please load processed pbmc3k dataset.”,
“step 2: Could you load processed pbmc68k dataset?”,
“step 3: Please integrate the two datasets.”,
“step 4: Could you perform batch correction on the combined data?”,
“step 5: Can you create scatter plot in UMAP basis?”
]
}
~~~

#### A.5.2 Prompt for task polishing

\# **Task Description**

Refine the subtask description by integrating essential parameters and their values from the docstring, ensuring their values are appropriate for the next steps in the code execution. Inherit any valid parameter values from the current subtask, verifying their accuracy and relevance. Check the docstring for API dependencies, required optional parameters, parameter conflicts, duplication, and deprecations. If the main goal provides a method name and the subtask can use this method to accomplish its goal, include the method name in the polished subtask. Include only the parameters with explicitly assigned values; avoid stating default values or parameters with vague values.

Provide only the refined subtask description. Avoid including any extraneous information.

—

Example:

original subtask description: Can you normalize the data and identify ‘target sum’ as median of total counts, ‘key’ with specific value, ‘max num’ as ‘10’, ‘log’ as ‘True’

thought: we delete some parameters, remove ‘target sum’ because the parameter ‘target sum’ just repeats its description and wasn’t assign valid value, remove ‘key’ as it wasn’t assign valid value, and the default value of the parameter ‘log’ is ‘True’ so we remove it, while only the ‘max num’ as ‘10’ is valid and non default and executable.

refined subtask description: Can you normalize the data and set ‘max num’ as ‘10’

Example:

Details to consider

Namespace variables:

~~~
{
“ result_1 “:
 “{
   ‘ type ‘: ‘ AnnData ‘,
   ‘ value ‘: Ann Data object with n_obs x n_vars = 3798 × 36601
     obs: ‘ in_tissue ‘, ‘ array_row ‘, ‘ array_col ‘
     var: ‘ gene_ids ‘, ‘ feature_types ‘, ‘ genome ‘
     uns: ‘ spatial ‘, ‘pca ‘
     obsm : ‘ spatial ‘, ‘ X_pca ‘
     varm : ‘PCs ‘
 }”
}
~~~

Extract necessary parameter details and constraints from API Docstring:

~~~
def squidpy. gr. ripley (adata =$, cluster_key =@, mode =$, spatial_key =@, metric=@, n_neigh =@,
 n_simulations =@, n_observations =@, max_dist=@, n_steps=@, seed =@, copy =@):}
~~~

original subtask description: Can you calculate Ripley’s statistics?

refined subtask description: Can you calculate Ripley’s statistics with ‘cluster key’ set as ‘array row’?

Example:

main goal: Use Scanpy to finish trajectory inference using the PAGA method. original subtask description: Please perform trajectory inference.

refined subtask description: Please perform trajectory inference using the PAGA method.

Example:

Main goal: Use Scanpy to conduct gene annotation on dataset 3k PBMCs. original subtask description: Load the built-in dataset.

refined subtask description: Load the built-in dataset 3k PBMCs.

—

Details to consider

Understand context and dependencies from past executed code: {execution_history}

Ensure parameter compatibility for existing namespace variables: {namespace_variables}

Extract necessary parameter details and constraints from API Docstring: {API_docstring}

Main goal (that includes this subtask as a step): {main_goal}

original Subtask description: {current_subtask}

refined subtask description:

\# **Example Input**:

`Execution_history = “““`

~~~
from scanpy. datasets import ebi_expression_atlas
result_1 = ebi_expression_atlas (‘E-GEOD -98816 ‘)
from scanpy. pp import filter_genes
result_2 = filter_genes (result_1, min_counts =100, inplace = True)
“““
~~~

Namespace variables =

~~~
{“ result_1 “:
 “{
   ‘ type ‘: ‘ AnnData ‘,
   ‘ value ‘: Ann Data object with n_obs x n_vars = 3186 × 21904
     obs: ‘ Sample Characteristic [ organism ]’, ‘ Sample Characteristic Ontology Term [
 organism ]’, ‘ Sample Characteristic [ strain ]’, ‘ Sample Characteristic Ontology Term [ strain
 ]’, ‘ Sample Characteristic [ age]’, ‘ Sample Characteristic Ontology Term [ age]’, ‘ Sample
 Characteristic [ developmental stage ]’, ‘ Sample Characteristic Ontology Term [ developmental
 stage ]’, ‘ Sample Characteristic [ organism part]’, ‘ Sample Characteristic Ontology Term [
 organism part]’, ‘ Sample Characteristic [ genotype ]’, ‘ Sample Characteristic Ontology Term [
 genotype ]’, ‘ Factor Value [ single cell identifier]’, ‘ Factor Value Ontology Term [ single
 cell identifier ]’
     var: ‘n\ _counts ‘
 }
}
~~~

`API docstring = “““`

~~~
def scanpy. pp. filter_cells (data : AnnData, min_counts : Optional[ int] = None, min_genes: Optional[ int] = None, max_counts :
 Optional[ int] = None, max_genes: Optional[ int] = None,
 inplace : bool = True, copy : bool = False):} \\
 “““ Filter cell outliers based on counts and numbers of genes expressed.
 For instance, only keep cells with at least ‘ min_counts ‘ counts or ‘
 min_genes ‘ genes expressed. This is to filter measurement outliers,
 i. e. “ unreliable “ observations.
 Only provide one of the optional parameters ‘ min_counts ‘, ‘ min_genes ‘, ‘
 max_counts ‘, ‘ max_genes ‘ per call.
 Parameters
 ----------
 data
 The (annotated) data matrix of shape ‘ n_obs ‘ x ‘ n_vars ‘. Rows correspond to cells and columns to genes.
 min_counts
 Minimum number of counts required for a cell to pass filtering.
 min_genes
 Minimum number of genes expressed required for a cell to pass filtering.
 max_counts
 Maximum number of counts required for a cell to pass filtering.
 max_genes
 Maximum number of genes expressed required for a cell to pass filtering.
 inplace
 Perform computation inplace or return result.
 Returns
 -------
 Depending on ‘ inplace ‘, returns the following arrays or directly subsets
 and annotates the data matrix :
 cells_subset
   Boolean index mask that does filtering. ‘ True ‘ means that the cell is kept. ‘ False ‘
 means the cell is removed.
number_per_cell
   Depending on what was thresholded (‘ counts ‘ or ‘ genes ‘), the array stores ‘ n_counts ‘
or ‘ n_cells ‘ per gene .”““
~~~

`Main goal = “““How can I filter out low-quality genes and cells using Scanpy?”““`

`Current subtask = “““Next, filter out low-quality cells using the ‘filter_cells’ method by specifying the appropriate thresholds for counts or genes expressed.”““`

\# **Example Output**:

Next, filter out low-quality cells using the `filter_cells` method by setting ‘min_counts’ as 100 and ‘inplace’ as True.

#### A.5.3 Prompt for task summarization

\# **Task Description**

Help to summarize the task with its solution in layman tone, it should be understandable by non professional programmers. Starts from description of the user query {user_query} and includes the selected API function’s Docstring {API_docstring}. Please use the template as ‘The task is …, we solved it by ‘ The response should be no more than two sentences

\# **Example Input**:

User_query: Let’s visualize the clustering results using the ‘leiden’ method to assess the impact of batch correction.

API_docstring:

~~~
def scanpy. tl. leiden (adata : AnnData, resolution : float = 1, restrict_to : Optional[ tuple [ str,
 Sequence [ str ]]] = None, random_state : Optional[ Union [ int, Random State ]] = 0, key_added :
 str = “ leiden “, adjacency : Optional[ spmatrix ] = None, directed :
 bool = True, use_weights : bool = True, n_iterations : int = -1, partition_type : Optional[ type [ Mutable Vertex Partition
 ]] = None, neighbors_key : Optional[ str] = None, obsp : Optional[ str] = None, copy : bool =
 False):
 “““
 Cluster cells into subgroups [ Traag18 ] _.
 Cluster cells using the Leiden algorithm [ Traag18 ]_,
 an improved version of the Louvain algorithm [ Blondel08 ] _.
 It has been proposed for single - cell analysis by [ Levine15 ] _.
 This requires having ran : func:’∼ scanpy. pp. neighbors ‘ or
 : func:’∼ scanpy. external. pp. bbknn ‘ first.
 Parameters
 ----------
 adata : Ann Data
   The annotated data matrix.
 resolution : float, optional
   A parameter value controlling the coarseness of the clustering.
   Higher values lead to more clusters.
   Set to ‘ None ‘ if overriding ‘ partition_type ‘
   to one that doesn ‘ t accept a ‘ resolution_parameter ‘.
 random_state : int or Random State, optional
   Change the initialization of the optimization.
 restrict_to : tuple [ str, Sequence [ str]], optional
   Restrict the clustering to the categories within the key for sample
   annotation, tuple needs to contain ‘(obs_key, list_of_categories) ‘.
 key_added : str, optional
   ‘ adata. obs ‘ key under which to add the cluster labels.
 adjacency : spmatrix, optional
   Sparse adjacency matrix of the graph, defaults to neighbors connectivities.
 directed : bool, optional
   Whether to treat the graph as directed or undirected.
 use_weights : bool, optional
   If ‘ True ‘, edge weights from the graph are used in the computation
   (placing more emphasis on stronger edges).
 n_iterations : int, optional
   How many iterations of the Leiden clustering algorithm to perform.
   Positive values above 2 define the total number of iterations to perform,
   -1 has the algorithm run until it reaches its optimal clustering.
 partition_type : type [ MutableVertexPartition ], optional
   Type of partition to use.
   Defaults to : class :’∼ leidenalg. RBConfiguration Vertex Partition ‘.
   For the available options, consult the documentation for
   : func:’∼ leidenalg. find_partition ‘.
 neighbors_key : str, optional
   Use neighbors connectivities as adjacency.
   If not specified, leiden looks. obsp [‘ connectivities ‘] for connectivities
   (default storage place for pp. neighbors).
   If specified, leiden looks
 . obsp [. uns[ neighbors_key ][‘ connectivities_key ‘]] for connectivities.
 obsp : str, optional
   Use. obsp [ obsp ] as adjacency. You can ‘ t specify both ‘
   obsp ‘ and ‘ neighbors_key ‘ at the same time.
 copy : bool, optional
   Whether to copy ‘ adata ‘ or modify it inplace.
 ** partition_kwargs
   Any further arguments to pass to ‘∼ leidenalg. find_partition ‘
   (which in turn passes arguments to the ‘ partition_type ‘).
 Returns
 -------
 adata. obs[ key_added ] : pandas. Series
   Array of dim (number of samples) that stores the subgroup id
   (‘‘0 ‘‘, ‘‘1 ‘‘, …) for each cell.
 adata. uns[‘ leiden ‘][‘ params ‘] : dict
   A dict with the values for the parameters ‘ resolution ‘, ‘ random_state ‘,
   and ‘ n_iterations ‘.
 “““
~~~

\# **Example Output**:

The task is to visualize the results of grouping cells into clusters using the ‘leiden’ method, which helps us see how well our batch correction worked. We solved it by using the ‘scanpy.tl.leiden’ function that clusters the data based on a set of parameters, allowing us to analyze the cell groups effectively.

#### A.5.4 Prompt for instruction generation

\# **Instruction**:

Generate 10 examples that use the function {API_name} (…) to train an intelligent code assistant for computational biology. Each example should be a JSON dictionary with fields `“instruction”` and `“code”`. The `“instruction”` field should be a single line of language instruction that a user might say to the code assistant. The `“code”` field should be a single line of code that the code assistant should generate based on the user’s instruction.

Be specific when writing the user instructions—craft them as if a human user would naturally speak to the assistant. Do not include the function name or parameter names explicitly. Vary the tone of the queries to include polite, straightforward, and casual styles. Emphasize the intent and the methods of the function, clearly distinguishing between the Target and Contrasting descriptions.

For example, given a Target description `“Scatter plot in UMAP basis”` and a Contrasting description `[“Scatter plot in PHATE basis”]`, the generated example `“Please create a scatter plot in the UMAP basis”` should convey the intent `“create a scatter plot”` and the detail `“in the UMAP basis”`, while avoiding specifics related to `“PHATE”`.

Now I provide my target and contrasting descriptions to you:

Target_description: {`target description`}

Contrasting descriptions: {`contrasting descriptions`}

Refer to the following code documentation for more information on what each function does and the distinctions between Target and Contrasting descriptions:

Docstring: {`API docstring`}

Now, generate the 10 examples following the instructions above:

\# **Example_Input**:

API_name: “scanpy.pl.ranking(…)”,

Target_description: “Plot rankings. See, for example, how this is used in pl.pca_loadings.”,

Contrasting_descriptions:

~~~
{
 “ Plot PCA results. The parameters are the ones of the scatter plot. Call pca_ranking
 separately .”,
 “ Plot ranking of genes using dotplot plot (see : func: scanpy. pl. dotplot).”,
 “ Scatter plot in PCA coordinates. Use the parameter annotate_var_explained to annotate the
   explained variance .”,
 “ Principal component analysis [ Pedregosa 11 ]. Computes PCA coordinates, loadings and
 variance decomposition .”,
 “ Plot ranking of genes using matrixplot plot (see : func: scanpy. pl. matrixplot).”
}
~~~

API_docstring:

~~~
def scanpy. pl. ranking (adata : AnnData, attr: Literal[ var, obs, uns, varm, obsm ], keys: Union [
 str, Sequence [ str]], color: None = black, dictionary : None = None, include_lowest : None =
 False):
 “““ Plot rankings. See, for example, how this is used in pl. pca_loadings.
 Parameters
 ----------
 adata
   The data .}
 attr
   The attribute of Ann Data that contains the score .}
 keys
   The scores to look up an array from the attribute of adata .”““}
~~~

\# **Example Output**:

[{“instruction”: “Please plot rankings using the specified keys from the attribute of the data.”, “code”: “scanpy.pl.ranking(adata, attr, keys)” },

{“instruction”: “Generate a rankings plot based on the provided attribute and keys.”, “code”: “scanpy.pl.ranking(adata, attr, keys)” },

{“instruction”: “I need to visualize rankings, can you use the attribute data for this plot?”, “code”: “scanpy.pl.ranking(adata, attr, keys)” },

{“instruction”: “Plot rankings from the specified data attribute using the given keys.”, “code”: “scanpy.pl.ranking(adata, attr, keys)” },

{“instruction”: “Could you create a rankings plot utilizing attribute data and specific keys?”, “code”: “scanpy.pl.ranking(adata, attr, keys)” },

{“instruction”: “Show me rankings with the attribute data and chosen keys in the plot.”, “code”: “scanpy.pl.ranking(adata, attr, keys)” },

{“instruction”: “I want to view rankings, can you plot this using the attribute data and keys provided?”, “code”: “scanpy.pl.ranking(adata, attr, keys)” },

{“instruction”: “Plot rankings using the attribute data and keys you have.”, “code”: “scanpy.pl.ranking(adata, attr, keys)” },

{“instruction”: “Please create a rankings plot by specifying the attribute and keys to use.”, “code”: “scanpy.pl.ranking(adata, attr, keys)” },

{“instruction”: “Generate a plot of rankings based on the attribute and keys provided for the data.”, “code”: “scanpy.pl.ranking(adata, attr, keys)” }]

#### A.5.5 Prompt for API prediction

\# **Task Description**:

Select the candidate that conveys the most information from the given instruction, and return it as a list in the format `[function1]`. Learn from the examples provided on how to choose candidates with maximum informative content.

Incontext example: {incontext_example}

Now finish the selection task for the below instruction. Never mess up with the examples above. Function candidates: {function_candidate}

Instruction: {query}

Function:

\# **Example_Input**: **Incontext Example = “““**:

function candidates:

0:scanpy.Neighbors.compute transitions, description: Compute transition matrix.

1:scanpy.Neighbors.compute eigen, description: Compute eigen decomposition of transition matrix.

2:scanpy.tl.pca, description: Principal component analysis [Pedregosa11]. Computes PCA coordinates, loadings and variance decomposition. Uses the implementation of *scikit-learn* [Pedregosa11].

Instruction: Can you compute the eigen decomposition of the transition matrix with a random state of 123?

Function: `[scanpy.Neighbors.compute eigen]`

—

function candidates:

0:scanpy.pl.MatrixPlot.style, description: Modifies plot visual parameters. 1:scanpy.pl.StackedViolin.style, description: Modifies plot visual parameters. 2:scanpy.pl.DotPlot.style, description: Modifies plot visual parameters.

Instruction: Fine-tune the edge line width to 0.2 Function: `[scanpy.pl.MatrixPlot.style]`

—

function candidates:

0:scanpy.Neighbors.to igraph, description: Generate igraph from connectivities.

1:scanpy.pl.draw graph, description: Scatter plot in graph-drawing basis.

2:scanpy.tl.draw graph, description: Force-directed graph drawing [Islam11] [Jacomy14] [Chippada18]. An alternative to tSNE that often preserves the topology of the data better. This requires to run `scanpy.pp.neighbors` first.

Instruction: Create a force-directed graph using the ‘Fa’ layout for me. Function: `[scanpy.tl.draw graph]`

—

function candidates:

0:scanpy.tl.embedding density, description: Calculate the density of cells in an embedding (per condition). Gaussian kernel density estimation is used to calculate the density of cells in an embedded space.

1:scanpy.pl.embedding, description: Scatter plot for user specified embedding basis (e.g. umap, pca, etc). 2:scanpy.pl.embedding density, description: Plot the density of cells in an embedding (per condition).

Instruction: I would like to visualize the gaussian kernel density estimates over condition, can you plot it? Function: `[scanpy.pl.embedding density]`

—

function candidates:

0:scanpy.pl.rank_genes_groups dotplot, description: Plot ranking of genes using dotplot plot (see

`scanpy.pl.dotplot`).

1:scanpy.pl.rank_genes_groups, description: Plot ranking of genes.

2:scanpy.pl.pca_loadings, description: Rank genes according to contributions to PCs. Instruction: Rank genes using contributions to PCs and display the plot.

Function: `[scanpy.pl.pca_loadings]`

**”””**

Function candidate:

0:”scanpy.pl.ranking”: “Plot rankings.See, for example, how this is used in pl.pca ranking.”

1:”scanpy.pl.rank_genes_groups”: “Plot ranking of genes.”

2:”scanpy.pl.rank_genes_groups dotplot”: “Plot ranking of genes using dotplot plot (see

:func:’ scanpy.pl.dotplot’)”

Query: “Please visualize the rankings based on my data. #

**Example Output**:

[“scanpy.pl.ranking”]

#### A.5.6 Prompt for argument prediction

##### A.5.6.1 Argument prediction for Literal type

\# **Parameter type: Literal**

\## **Task Description**:

Determine from the API DOCSTRING and USER QUERY whether each parameter needs a predicted value. Based on the parameter type and meaning, infer if it needs to fill in a value or modify the default value, and return all the parameters that need prediction or modification with their predicted values.

Instructions:

1. Extract values and assign to parameters: If there are values explicitly appearing in the USER QUERY, such as those quoted with “, values with underscores like value value, or list/tuple/dict like `[‘A’, ‘B’]`, they are likely to be actual parameter values. When handling these values: Boolean values: If we are predicting boolean values, you can ignore these explicitly mentioned values. Integer/float/str/list/tuple values: If we are predicting these types of values, you can assign them to the most corresponding parameters based on the modifiers of the extracted values.
2. Extract and convert values if necessary: Some values might not be obvious and require conversion (e.g., converting “Diffusion Map” to “diffmap” if “Diffusion Map” appears in the user query, as this is the correct usage for some API. Another example is converting “blank space delimiter” to `“delimiter=‘ ‘”`). Ensure that the returned values are correctly formatted, case-sensitive, and directly usable in API calls.
3. Identify relevant parameters: From the USER QUERY and API DOCSTRING, identify which parameters require value extraction or default value modification based on their type and meaning.
4. Never generate fake values: Only include values explicitly extracted in the USER QUERY or that can be clearly inferred. Do not generate fake values for remaining parameters.
5. Response format: Format results as `“param name”: “extracted value”`, ensuring they are executable and match the correct type from the API DOCSTRING. For example, return `[‘a’, ‘b’]` for list type instead of Tuple `(‘a’, ‘b’)`. Only return parameters with non-empty values (i.e., not None).
6. Avoid complexity: Avoid using ‘or’ in parameter values and ensure no overly complex escape characters. The format should be directly loadable by `json.loads`.
7. Distinguish similar parameters: For similar parameters (e.g., ‘knn’ and ‘knn max’), distinguish them in meaning and do not mix them up. If there are two parameter candidates with the same meaning for one value, fill in both with the extracted value.
8. Maintain Format and transfer format if necessary: Ensure the final returned values match the expected parameter type, including complex structures like `[(“keyxx”, 1)]` for parameters like “obsm keys” with types like `Iterable[tuple[str, int]]`. If USER QUERY contains list, tuple, or dictionary values, extract them as a whole and keep the format as a value.

Find the candidate with the same level of informativeness. If there are multiple candidates containing the keyword, but the query provides only minimal information, select the candidate with the least information content. If a parameter name is mentioned without a value, skip it.

Examples:

USER QUERY: `“… with basis ‘Diffusion Map’” =>`{`“basis”:’diffmap’`}

USER QUERY: `“… blank space delimiter” =>`{`“delimiter”:’ ‘`}

Now start extracting the values with parameters from USER QUERY based on parameters descriptions and default value from API DOCSTRING

USER QUERY: {user_query}

API DOCSTRING: {API_docstring}

param list: {parameters_name_list}

Return parameter with extracted value in format: { `“param1”:”value1”, “param2”:”value2”, …` }.

Only return JSON format answer in one line, do not return other descriptions.

\# **Example_Input**:

User query: Could you annotate the highly variable genes with ‘flavor’ set as ‘cell_ranger’?

API_docstring:

~~~
def scanpy. pp. highly_variable_genes (flavor: Literal[ seurat, cell_ranger, seurat_v 3 ] = seurat):
 “““ Annotate highly variable genes [ Satija15 ] _ [ Zheng17 ] _ [ Stuart19 ] _. Expects logarithmized
   data, except when ‘ flavor=‘ seurat_v3 ‘‘, in which count
 data is expected.
 Depending on ‘ flavor ‘, this reproduces the R-implementations of Seurat
 [ Satija15 ]_, Cell Ranger [ Zheng17 ]_, and Seurat v3 [ Stuart19 ] _.
 For the dispersion - based methods ([ Satija15 ] _ and [ Zheng17 ] _), the normalized
 dispersion is obtained by scaling with the mean and standard deviation of
 the dispersions for genes falling into a given bin for mean expression of
 genes. This means that for each bin of mean expression, highly variable
 genes are selected.
 For [ Stuart19 ]_, a normalized variance for each gene is computed. First, the data
 are standardized (i. e., z-score normalization per feature) with a regularized
 standard deviation. Next, the normalized variance is computed as the variance
 of each gene after the transformation. Genes are ranked by the normalized variance.
 See also ‘ scanpy. experimental. pp. _highly_variable_genes ‘ for additional flavours
 (e. g. Pearson residuals).
 Parameters
 ----------
 flavor
   Choose the flavor for identifying highly variable genes. For the dispersion
 based methods in their default workflows, Seurat passes the cutoffs whereas
 Cell Ranger passes ‘ n_top_genes ‘.
 “““
~~~

Parameters name list:

~~~
[‘ flavor ‘]
~~~

\# **Example Output**:

~~~
{‘ flavor ‘: ‘ cell_ranger ‘}
~~~

##### A.5.6.2 Argument prediction for Bool type

\# **Parameter type: Boolean**

\## **Task Description**:

Instructions:

Determine the state (True/False) based on the full USER QUERY and the parameter description, rather than just a part of it. If the ‘True/False’ value is not explicitly mentioned but the meaning is implied in the USER QUERY, still predict the parameter.

Examples:

USER QUERY: `“… with logarithmic axes” =>`{`“log”:”True”`}

USER QUERY: `“… without logarithmic axes” =>` { `“log”:”False”`}

USER QUERY: `“… hide the default legends and the colorbar legend” =>`{`“show”:”False”, “show colorbar”:”False”`}

USER QUERY: `“… and create a copy of the data structure” =>` {`“copy”: “True”`}, do not predict “data”

USER QUERY: {user_query}

API DOCSTRING: {API_docstring}

param list: {parameters_name_list}

Return parameter with extracted value in format: {`“param1”:”value1”, “param2”:”value2”, …` }.

Only return JSON format answer in one line, do not return other descriptions.

\# **Example_Input**:

User query: Could you conduct principal component analysis with ‘zero center’ set as ‘False’? API docstring:

~~~
def scanpy. pp. pca(zero_center : Optional[ bool] = True, return_info : bool = False,
 use_highly_variable : Optional[ bool] = None, copy : bool = False, chunked : bool = False):
“““ Principal component analysis [ Pedregosa 11 ] _. Computes PCA coordinates, loadings and
 variance decomposition.
Uses the implementation of * scikit - learn * [ Pedregosa 11 ] _.
 .. versionchanged :: 1.5.0
   In previous versions, computing a PCA on a sparse matrix would make a dense copy of
   the array for mean centering.
   As of scanpy 1.5.0, mean centering is implicit.
   While results are extremely similar, they are not exactly the same.
   If you would like to reproduce the old results, pass a dense array.
 Parameters
 ----------
 zero_center
   If ‘ True ‘, compute standard PCA from covariance matrix.
 If ‘ False ‘, omit zero - centering variables
 (uses : class :’∼ sklearn. decomposition. Truncated SVD ‘),
 which allows to handle sparse input efficiently.
 Passing ‘ None ‘ decides automatically based on sparseness of the data.
 return_info
   Only relevant when not passing an : class :’∼ anndata. AnnData ‘:
 see “** Returns **”.
 use_highly_variable
   Whether to use highly variable genes only, stored in
 ‘. var[‘ highly_variable ‘]’.
 By default uses them if they have been determined beforehand.
 copy
   If an : class :’∼ anndata. AnnData ‘ is passed, determines whether a copy
 is returned. Is ignored otherwise.
 chunked
   If ‘ True ‘, perform an incremental PCA on segments of ‘ chunk_size ‘.
 The incremental PCA automatically zero centers and ignores settings of
 ‘ random_seed ‘ and ‘ svd_solver ‘. If ‘ False ‘, perform a full PCA.
 “““
~~~

Parameters name list:

~~~
[‘ zero_center ‘, ‘ return_info ‘, ‘ use_highly_variable ‘, ‘ copy ‘, ‘ chunked ‘]
~~~

\# **Example Output**:

~~~
{‘ zero_center ‘: ‘ False ‘}
~~~

##### A.5.6.3 Argument prediction for Other type

\# **Parameter type: int, float, str, list[str], tuple[str] and all other types**

\## **Task Description**:

Instructions:

Choose only parameters with explicitly mentioned values in the USER QUERY. If a parameter name is mentioned without a value (e.g., “a specific group/key” where “group/key” is the parameter name but no value is given), ignore that parameter.

Examples:

USER QUERY: `“… with chunk size 6000” =>` {`“chunked”:”True”, “chunk size”:”6000”`}

USER QUERY: `“… with groups [‘A’, ‘B’, ‘C’]” =>` {`“groups”:[‘A’, ‘B’, ‘C’]`}

USER QUERY: `“… with at least 100 counts.” =>` {`“min_counts”: 100`}

USER QUERY: `“… for a specific layer ‘X new’?” =>` {`“layer”: “X new”` }, do not assign

“X new” to “X”

USER QUERY: { user query}

API DOCSTRING: {API_docstring}

param list: {parameters_name_list}

Return parameter with extracted value in format: { `“param1”:”value1”, “param2”:”value2”, …` }.

Only return JSON format answer in one line, do not return other descriptions.

\# **Example_Input**:

User query: Can you draw a UMAP plot for the clusters using ‘color’ set as ‘louvain’? API docstring:

~~~
def scanpy. pl. umap (color: Optional[ Union [ str, Sequence [ str ]]] = None, gene_symbols : Optional[
 str] = None, edges_width : float = 0.1, edges_color : Union [ str, Sequence [ float], Sequence [
 str ]] = grey, neighbors_key : Optional[ str] = None, groups: Optional[ str] = None,
 components : Union [ str, Sequence [ str ]] = None, dimensions: Optional[ Union [ tuple [ int, int],
 Sequence [ tuple [ int, int ]]]] = None, layer: Optional[ str] = None, scale_factor : Optional[
 float] = None, color_map : Optional[ Union [ Colormap, str ]] = None, cmap : Optional[ Union [
 Colormap, str ]] = None, palette : Optional[ Union [ str, Sequence [ str], Cycler ]] = None,
 na_color: Union [ str, tuple [ float, Ellipsis ]] = lightgray, size : Optional[ Union [float,
 Sequence [ float ]]] = None, legend_loc : str = right margin, legend_fontoutline : Optional[ int
 ] = None, colorbar_loc : Optional[ str] = right, vmax : Optional[ Union [ str, float, Callable [
 Sequence [ float], float], Sequence [ Union [ str, float, Callable [ Sequence [ float], float ]]]]] =
  None, vmin : Optional[ Union [ str, float, Callable [ Sequence [ float], float], Sequence [ Union [
 str, float, Callable [ Sequence [ float], float ]]]]] = None, vcenter: Optional[ Union [ str,
 float, Callable [ Sequence [ float], float], Sequence [ Union [ str, float, Callable [ Sequence [
 float], float ]]]]] = None, norm : Optional[ Union [ Normalize, Sequence [ Normalize ]]] = None,
 outline_width : tuple [ float, float] = (0.3, 0.05), outline_color : tuple [ str, str] = (‘ black
 ‘, ‘ white ‘), ncols: int = 4, hspace : float = 0.25, wspace : Optional[ float] = None, title :
 Optional[ Union [ str, Sequence [ str ]]] = None):
 “““ Scatter plot in UMAP basis.
 Parameters
 ----------
 color
   Keys for annotations of observations / cells or variables/ genes, e. g.,
 ‘‘ ann1 ‘‘ or ‘[‘ ann1 ‘, ‘ ann2 ‘]’.
 gene_symbols
   Column name in ‘. var ‘ Data Frame that stores gene symbols. By default ‘ var_names ‘
 refer to the index column of the ‘. var ‘ Data Frame. Setting this option allows
 alternative names to be used.
 edges_width
   Width of edges.
 edges_color
   Color of edges. See : func:’∼ networkx. drawing. nx_pylab. draw_networkx_edges ‘.
 neighbors_key
   Where to look for neighbors connectivities.
 If not specified, this looks. obsp [‘ connectivities ‘] for connectivities
 (default storage place for pp. neighbors).
 If specified, this looks
. obsp [. uns[ neighbors_key ][‘ connectivities_key ‘]] for connectivities.
 groups
   Restrict to a few categories in categorical observation annotation.
 The default is not to restrict to any groups.
 components
   For instance, ‘[‘1, 2 ‘, ‘2, 3 ‘]’. To plot all available components use ‘ components =‘ all ‘‘.
 dimensions
   0 - indexed dimensions of the embedding to plot as integers. E. g. [(0, 1), (1, 2)].
 Unlike ‘ components ‘, this argument is used in the same way as ‘ colors ‘, e. g. is
 used to specify a single plot at a time. Will eventually replace the components
 argument.
 layer
   Name of the Ann Data object layer that wants to be plotted. By default
 adata. raw. X is plotted. If ‘ use_raw = False ‘ is set, then ‘ adata .X’ is plotted.
 If ‘ layer ‘ is set to a valid layer name, then the layer is plotted. ‘ layer ‘
 takes precedence over ‘ use_raw ‘.
 color_map
   Color map to use for continous variables. Can be a name or a
 : class :’∼ matplotlib. colors. Colormap ‘ instance (e. g. ‘“ magma ‘“, ‘“ viridis”‘
 or ‘ mpl. cm. cividis ‘), see : func:’∼ matplotlib. cm. get_cmap ‘.
 If ‘ None ‘, the value of ‘ mpl. rcParams [“ image. cmap “]’ is used.
 The default ‘ color_map ‘ can be set using : func:’∼ scanpy. set_figure_params ‘.
 palette
   Colors to use for plotting categorical annotation groups.
 The palette can be a valid : class :’∼ matplotlib. colors. Listed Colormap ‘ name
 (‘‘ Set2 ‘‘, ‘‘ tab20 ‘‘, …), a : class :’∼ cycler. Cycler ‘ object, a dict mapping
 categories to colors, or a sequence of colors. Colors must be valid to
 matplotlib. (see : func:’∼ matplotlib. colors. is_color_like ‘).
 If ‘ None ‘, ‘ mpl. rcParams [“ axes. prop_cycle “]’ is used unless the categorical
 variable already has colors stored in ‘ adata. uns [“{ var} _colors “]’.
 If provided, values of ‘ adata. uns [“{ var} _colors “]’ will be set.
 na_color
   Color to use for null or masked values. Can be anything matplotlib accepts as a
 color. Used for all points if ‘ color=None ‘.
 size
   Point size. If ‘ None ‘, is automatically computed as 120000 / n_cells.
 Can be a sequence containing the size for each cell. The order should be
 the same as in adata. obs.
 legend_loc
   Location of legend, either ‘‘ on data ‘‘, ‘‘ right margin ‘‘ or a valid keyword
 for the ‘loc ‘ parameter of : class :’∼ matplotlib. legend. Legend ‘.
 legend_fontoutline
   Line width of the legend font outline in pt. Draws a white outline using
 the path effect : class :’∼ matplotlib. patheffects. with Stroke ‘.
 colorbar_loc
   Where to place the colorbar for continous variables. If ‘ None ‘, no colorbar is added.
 vmax
   The value representing the upper limit of the color scale. The format is the same as
 for ‘ vmin ‘.
 vmin
   The value representing the lower limit of the color scale. Values smaller than vmin
 are plotted
 with the same color as vmin. vmin can be a number, a string, a function or ‘ None ‘. If
 vmin is a string and has the format ‘pN ‘, this is interpreted as a vmin = percentile (N).
 For example vmin =‘ p1 .5 ‘ is interpreted as the 1.5 percentile. If
 vmin is function, then vmin is interpreted as the return value of the function over the list of values to plot.
 For example to set vmin tp the mean of the values to plot, ‘ def my_vmin (values): return
 np. mean (values)’ and then set ‘ vmin = my_vmin ‘. If vmin is None (default) an automatic
 minimum value is used as defined by matplotlib ‘ scatter ‘ function. When making multiple
 plots, vmin can be a list of values, one for each plot. For example ‘ vmin =[0.1, ‘p1 ‘, None
, my_vmin ]’
 vcenter
   The value representing the center of the color scale. Useful for diverging colormaps.
 The format is the same as for ‘ vmin ‘.
 Example : sc. pl. umap (adata, color=‘ TREM2 ‘, vcenter=‘ p50 ‘, cmap =‘ RdBu_r ‘) outline_width
   Tuple with two width numbers used to adjust the outline. The first value is the width
 of the border color as a fraction of the scatter dot size (default: 0.3). The second value
 is
 width of the gap color (default: 0.05).
 outline_color
 Tuple with two valid color names used to adjust the add_outline. The first color is
 the
 border color (default: black), while the second color is a gap color between the
 border color and the scatter dot (default: white).
 ncols
   Number of panels per row.
 hspace
   Adjust the height of the space between multiple panels.
 wspace
   Adjust the width of the space between multiple panels.
 title
   Provide title for panels either as string or list of strings,
 e. g. ‘[‘ title1 ‘, ‘ title2 ‘, …] ‘.
 “““
~~~

Parameters_name_list:

~~~
[‘ color ‘, ‘ gene_symbols ‘, ‘ edges_width ‘, ‘ edges_color ‘, ‘ neighbors_key ‘, ‘ groups ‘, ‘ components ‘, ‘
 dimensions ‘, ‘ layer ‘, ‘ scale_factor ‘, ‘ color_map ‘, ‘ cmap ‘, ‘ palette ‘, ‘ na_color ‘, ‘
 size ‘, ‘ legend_loc ‘, ‘ legend_fontoutline ‘, ‘ colorbar_loc ‘, ‘ vmax ‘, ‘ vmin ‘, ‘ vcenter ‘, ‘
 norm ‘, ‘ outline_width ‘, ‘ outline_color ‘, ‘ ncols ‘, ‘ hspace ‘, ‘ wspace ‘, ‘ title ‘]
~~~

\# **Example Output**:

~~~
{‘ color ‘: ‘ louvain ‘}
~~~

#### A.5.7 Prompt for code explanation

\# **Task Description**

Help to summarize the task with its solution in layman tone, it should be understandable by non professional programmers. Starts from description of the user query {user_query} and includes the selected API function {API_docstring}. The generated code is {execution code} Please use the template as ‘The task is …, we solved it by ‘

The response should be no more than three sentences. Additionally, the interpretation encompasses explanations for the parameters utilized in the generated code.

\# **Example_Input**:

User query: Could you visualize the gene expression values through dotplot with `adata` set as `result 1`, `var_names` set as a valid subset of `result 1.var_names`, and `groupby` set as `louvain`?

API_docstring:

~~~
def scanpy. pl. dotplot(adata : AnnData, var_names: Union [ str, Sequence [ str], Mapping [ str, Union [ str, Sequence [ str ]]]], groupby : Union[
 str, Sequence [ str]], use_raw : Optional[ bool] = None,
  log: bool = False, num_categories : int = 7, expression_cutoff : float = 0.0,
 mean_only_expressed : bool = False, cmap : str = Reds, dot_max : Optional[ float] = None,
 dot_min : Optional[ float] = None, standard_scale : Optional[ Literal[ var, group ]] = None,
 smallest_dot : Optional[ float] = 0.0, title : Optional[ str] = None, colorbar_title : Optional
 [ str] = Mean expression
in group, size_title : Optional[ str] = Fraction of cells
in group (%), figsize : Optional[ tuple [ float, float ]] = None, dendrogram : Union [ bool, str] =
 False, gene_symbols : Optional[ str] = None, var_group_positions : Optional[ Sequence [ tuple [
 int, int ]]] = None, var_group_labels : Optional[ Sequence [ str ]] = None, var_group_rotation :
 Optional[ float] = None, layer: Optional[ str] = None, swap_axes: Optional[ bool] = False,
 dot_color_df : Optional[ Data Frame ] = None, show : Optional[ bool] = None, save : Optional[
 Union [ str, bool ]] = None, ax: Optional[ _AxesSubplot ] = None, return_fig : Optional[ bool] =
 False, vmin : Optional[ float] = None, vmax : Optional[ float] = None, vcenter: Optional[ float
 ] = None, norm : Optional[ Normalize ] = None):
 “““ Makes a * dot plot* of the expression values of ‘ var_names ‘.
 For each var_name and each ‘ groupby ‘ category a dot is plotted.
 Each dot represents two values: mean expression within each category
 (visualized by color) and fraction of cells expressing the ‘ var_name ‘ in the
 category (visualized by the size of the dot). If ‘ groupby ‘ is not given,
 the dotplot assumes that all data belongs to a single category.
 .. note ::
   A gene is considered expressed if the expression value in the ‘ adata ‘ (or ‘
   adata. raw ‘) is above the specified threshold which is zero by default.
 An example of dotplot usage is to visualize, for multiple marker genes,
 the mean value and the percentage of cells expressing the gene
 across multiple clusters.
 This function provides a convenient interface to the :
 class :’∼ scanpy. pl. DotPlot ‘ class. If you need more flexibility, you should use : class :’∼ scanpy. pl. DotPlot ‘
 directly.
 Parameters
 ----------
 adata
   Annotated data matrix.
 var_names
   ‘ var_names ‘ should be a valid subset of ‘ adata. var_names ‘.
   If ‘ var_names ‘ is a mapping, then the key is used as label
   to group the values (see ‘ var_group_labels ‘). The mapping values
   should be sequences of valid ‘ adata. var_names ‘. In this
   case either coloring or ‘ brackets ‘ are used for the grouping
   of var_names depending on the plot. When ‘ var_names ‘ is a mapping,
   then the ‘ var_group_labels ‘ and ‘ var_group_positions ‘ are set.
   groupby
   The key of the observation grouping to consider.
 use_raw
   Use ‘raw ‘ attribute of ‘ adata ‘ if present.
 log
   Plot on logarithmic axis.
 num_categories
   Only used if groupby observation is not categorical. This value
   determines the number of groups into which the groupby observation
   should be subdivided.
 categories_order
   Order in which to show the categories. Note : add_dendrogram or add_totals
   can change the categories order.
 figsize
   Figure size when ‘ multi_panel =True ‘.
   Otherwise the ‘ rcParam [‘ figure. figsize ]’ value is used.
   Format is (width, height)
 dendrogram
   If True or a valid dendrogram key, a dendrogram based on the hierarchical
   clustering between the ‘ groupby ‘ categories is added.
   The dendrogram information is computed using : func:’ scanpy. tl. dendrogram ‘.
   If ‘ tl. dendrogram ‘ has not been called previously the function is called
   with default parameters.
 gene_symbols
   Column name in ‘. var ‘ Data Frame that stores gene symbols.
   By default ‘ var_names ‘ refer to the index column of the ‘. var ‘ Data Frame.
   Setting this option allows alternative names to be used.
 var_group_positions
   Use this parameter to highlight groups of ‘ var_names ‘.
   This will draw a ‘ bracket ‘ or a color block between the given start and end
   positions. If the parameter ‘ var_group_labels ‘ is set, the corresponding
   labels are added on top / left. E. g. ‘ var_group_positions =[(4, 10)]’
   will add a bracket between the fourth ‘ var_name ‘ and the tenth ‘ var_name ‘.
   By giving more positions, more brackets/ color blocks are drawn.
 var_group_labels
   Labels for each of the ‘ var_group_positions ‘ that want to be highlighted.
 var_group_rotation
   Label rotation degrees.
   By default, labels larger than 4 characters are rotated 90 degrees.
 layer
   Name of the Ann Data object layer that wants to be plotted. By default adata. raw. X is
 plotted.
   If ‘ use_raw = False ‘ is set, then ‘ adata .X’ is plotted. If ‘ layer ‘ is set to a valid
 layer name,
   then the layer is plotted. ‘ layer ‘ takes precedence over ‘ use_raw ‘.
 title
   Title for the figure
 colorbar_title
   Title for the color bar. New line character (\ n) can be used.
 cmap
   String denoting matplotlib color map.
 standard_scale
   Whether or not to standardize the given dimension between 0 and 1, meaning for
   each variable or group, subtract the minimum and divide each by its maximum.
 swap_axes
   By default, the x axis contains ‘ var_names ‘ (e. g. genes) and the y axis
   the ‘ groupby ‘ categories. By setting ‘ swap_axes ‘ then x are the
   ‘ groupby ‘ categories and y the ‘ var_names ‘.
 return_fig
   Returns : class:’ DotPlot ‘ object. Useful for fine – tuning
   the plot. Takes precedence over ‘ show = False ‘.
 size_title
   Title for the size legend. New line character (\ n) can be used.
 expression_cutoff
   Expression cutoff that is used for binarizing the gene expression and
   determining the fraction of cells expressing given genes. A gene is
   expressed only if the expression value is greater than this threshold.
 mean_only_expressed
   If True, gene expression is averaged only over the cells
   expressing the given genes.
 dot_max
   If none, the maximum dot size is set to the maximum fraction value found
   (e. g. 0.6). If given, the value should be a number between 0 and 1.
   All fractions larger than dot_max are clipped to this value.
 dot_min
   If none, the minimum dot size is set to 0. If given,
   the value should be a number between 0 and 1.
   All fractions smaller than dot_min are clipped to this value.
 smallest_dot
   If none, the smallest dot has size 0.
   All expression levels with ‘ dot_min ‘ are plotted with this size.
 show
   Show the plot, do not return axis.
 save
   If ‘ True ‘ or a ‘str ‘, save the figure.
   A string is appended to the default filename.
   Infer the filetype if ending on {‘‘. pdf ‘‘, ‘‘. png ‘‘, ‘‘. svg ‘‘}.
 ax
   A matplotlib axes object. Only works if plotting a single component.
 vmin
   The value representing the lower limit of the color scale. Values smaller than vmin
 are plotted
   with the same color as vmin.
 vmax
   The value representing the upper limit of the color scale. Values larger than vmax
 are plotted
   with the same color as vmax.
 vcenter
   The value representing the center of the color scale. Useful for diverging colormaps.
 norm
   Custom color normalization object from matplotlib. See ‘
   https://matplotlib.org/stable/tutorials/colors/colormapnorms.html ‘ for details.
 kwds
   Are passed to : func:’ matplotlib. pyplot. scatter ‘.
 Returns
 -------
 If ‘ return_fig ‘ is ‘ True ‘, returns a : class :’∼ scanpy. pl. DotPlot ‘ object, else if ‘ show ‘ is false, return axes dict
 See also
 --------
 : class :’∼ scanpy. pl. DotPlot ‘: The DotPlot class can be used to to control
    several visual parameters not available in this function.
 : func:’∼ scanpy. pl. rank_genes_groups_dotplot ‘: to plot marker genes identified using the : func:’∼ scanpy. tl. rank_genes_groups ‘ function.
 Examples
 --------
 Create a dot plot using the given markers and the PBMC example dataset grouped by the category ‘ bulk_labels ‘.
 .. plot ::
   : context: close - figs
   import scanpy as sc
   adata = sc. datasets. pbmc68 k_reduced ()
   markers = [‘ C1QA ‘, ‘ PSAP ‘, ‘ CD79A ‘, ‘ CD79B ‘, ‘ CST3 ‘, ‘LYZ ‘]
   sc. pl. dotplot(adata, markers, groupby =‘ bulk_labels ‘, dendrogram = True)
 Using var_names as dict:
 .. plot ::
   : context: close - figs
   markers = {‘T-cell ‘: ‘ CD3D ‘, ‘B-cell ‘: ‘ CD79A ‘, ‘ myeloid ‘: ‘ CST3 ‘} sc. pl. dotplot(adata, markers, groupby =‘ bulk_labels ‘, dendrogram = True)
   Get DotPlot object for fine tuning
 .. plot ::
   : context: close - figs
   dp = sc. pl. dotplot(adata, markers, ‘ bulk_labels ‘, return_fig = True)
   dp. add_totals (). style (dot_edge_color =‘ black ‘, dot_edge_lw =0.5). show ()
 The axes used can be obtained using the get_axes () method
 .. code - block :: python
   axes_dict = dp. get_axes ()
   print(axes_dict)”““ ‘. The generated code is ‘ result_plot = dotplot(result_1, var_names
 = list(result_1. var_names [0 :10 ]), groupby =‘ louvain ‘, use_raw = False, show = True) ‘.
~~~

Execution code:

~~~
result_plot = dotplot(result_1, var_names= list(result_1. var_names [0 :10 ]), groupby =‘ louvain ‘,
 use_raw = False, show = True)
~~~

\# **Example Output**:

The task is to create a visual representation, called a dot plot, that shows the expression levels of specific genes across different cell groups in a dataset. We solved it by using the ‘scanpy.pl.dotplot’ function, specifying a list of the first ten genes from our data (‘var_names’), grouping the results by the ‘louvain’ clustering method (‘groupby’), and opting not to use the raw data (‘use_raw=False’) for clarity in the results, while also displaying the plot on the screen (‘show=True’). This approach allows us to easily see which genes are expressed in each cell group and to what extent, helping in biological analysis and interpretation of the data.

#### A.5.8 Prompt for code troubleshooting

\# **Task Description**

\## **With or without solution from github issue**

\### With solution

`possible solution info = f”Possible solution from similar issues from Github Issue Discussion:` {`possible_solution`}”

\### Without solution

`possible solution info = ““`

\## **With or without tutorial examples**

\### With tutorial examples

possible solution info = f”Here are some examples. Identify the key prerequisite APIs or the correct usage of the target API to address the bug. Do not copy their parameters; predict based on the variable information in our namespace: {`API_examples`}.”

\### Without tutorial examples

`possible solution info = ““`

\## **Common Part**

Task: Analyze and correct the most recent failed Python script attempt based on the provided traceback information.

Make corrections considering all previous failed attempts and the associated error details, ensuring that prior mistakes are not repeated. You must make some valid and meaningful change, and keep the key correction step you did correctly in past attempt. Please ensure each time you make a correction, you are moving towards the correct solution.

Include all necessary library imports at the beginning.

Ensure the correct execution order for API dependencies like pandas, numpy, and AnnData, based on the traceback error and variables.

Use only variables from successful executions.

Remove unnecessary or incorrect parameters, ensuring required ones are in proper order. Adjust misused attributes or values, especially for AnnData object-specific attributes.

Preprocess data if needed using relevant tools or APIs before the main API call. Remove unnecessary optional parameters causing errors.

Respond with the corrected code in JSON format.

Refer to namespace variables for their values and attributes; avoid variables with None. The function returned variable should not be the same name as the function name.

Common Errors to Address:

Import Verification: Confirm necessary libraries are imported. API Usage: Replace or continue with the correct API as needed.

Parameter Handling: Streamline parameters to essentials, removing any incorrect or irrelevant ones. Prerequisite API Calls: Include any necessary pre-API steps.

Identify and address indirect errors by deducing the root cause. Present the logical steps in the ‘analysis’ section and the corresponding code in the ‘code’ section.

KeyError: Ensure to use the exist and correct attribute from existing variables in namespace. Or use the correct API to generate the attribute before executing this subtask. Sometimes the key error is due to that there lack of previous API.

TypeError: Sometimes the processed variable gets None, please use other variable in namespace instead of the null variable.

AttributeError: Sometimes the message ‘NoneType’ object indicates that the variable is None, verify that you’re not using a None variable and replace it with a valid one from the current namespace. Or check whether the annotations or parameters you’re accessing (e.g., .var, .obs, etc.) are indeed present in the object. It’s possible that the attribute you’re trying to use does not exist for this specific object or is incorrectly referenced.

ValueError: Be careful for choosing the attributes from ‘.obs’ and ‘.var’ of AnnData object, as some API requires annotations are from the same attribute.

Here are information:

Success execution History: { success_history_code}

Existing Namespace variables: { namespace_variables}

Current Goal: { goal_description}

History Failed Attempts with their tracebacks:

{error code}

{possible_solution_info}

{tutorial_examples}

API Docstring: {API_docstring}.

Response Format: “analysis”: “Explain how to correct the bug in 2 sentences, including the reason of the bug, and how to correct.”, “code”: “Contain the Corrected failed attempt code, exclude code from ‘success execution history’, exclude ‘save plot with timestamp()’, exclude any comments in the code. Notice to set any parameters named ‘inplace’, ‘show’ as True, while ‘copy’ as False”

\# **Example_Input**:

`Success history code = “““`

~~~
from scanpy. datasets import pbmc3 k_processed
result_1 = pbmc3 k_processed ()
from scanpy. pl import dotplot
“““
~~~

Namespace variables = “““

~~~
{“ result_1 “:
 “{
   ‘ type ‘: ‘ AnnData ‘,
   ‘ value ‘: Ann Data object with n_obs x n_vars = 2638 × 1838
     obs: ‘ n_genes ‘, ‘ percent_mito ‘, ‘ n_counts ‘, ‘ louvain ‘
     var: ‘ n_cells ‘
     uns: ‘ draw_graph ‘, ‘ louvain ‘, ‘ louvain_colors ‘, ‘ neighbors ‘, ‘pca ‘, ‘
 rank_genes_groups ‘
     obsm : ‘ X_pca ‘, ‘ X_tsne ‘, ‘ X_umap ‘, ‘ X_draw_graph_fr ‘
     varm : ‘PCs ‘
     obsp : ‘ distances ‘, ‘ connectivities ‘
 }”
}
“““
~~~

`Goal_description = “““`

`Could you visualize the gene expression values through dotplot with ‘adata’ set as ‘result 1’, ‘var_names’ set as a valid subset of ‘result 1.var_names’, and ‘groupby’ set as ‘louvain’?`

`“““`

`Error code = “““`

~~~
failed attempt 1:
result_2 = dotplot(result_1, result_1. var_names, ‘ louvain ‘, use_raw = result_1, standard_scale =‘
   var ‘, show = True)
 traceback : ‘ Traceback (most recent call last):
 File “/ home / z6 dong / Bio Chat/ refer/ src/ BioMANIA / src/ inference / execution_UI. py”, line 502, in
   execute_api_call
  exec(api_call_code, locals (), globals ())
 File “ < string >“, line 1, in <module >
 File “/ home / z6 dong / anaconda3 / envs/ biomania / lib / python 3 .10/ site - packages/ scanpy / plotting /
   _dotplot. py”, line 940, in dotplot
   dp = DotPlot(
 File “/ home / z6 dong / anaconda3 / envs/ biomania / lib / python 3 .10/ site - packages/ scanpy / plotting /
   _dotplot. py”, line 131, in
   Base Plot. __init__(__init__
 File “/ home / z6 dong / anaconda3 / envs/ biomania / lib / python 3 .10/ site - packages/ scanpy / plotting /
   _baseplot_class. py”, line 110, in __init__
   self. categories, self. obs_tidy = _prepare_dataframe (
 File “/ home / z6 dong / anaconda3 / envs/ biomania / lib / python 3 .10/ site - packages/ scanpy / plotting /
   _anndata. py”, line 1926, in _prepare_dataframe
   obs_tidy = get. obs_df(
 File “/ home / z6 dong / anaconda3 / envs/ biomania / lib / python 3 .10/ site - packages/ scanpy / get/ get. py”,
   line 291, in obs_df
   _get_obs_rep (adata, layer= layer, use_raw = use_raw),
 File “/ home / z6 dong / anaconda3 / envs/ biomania / lib / python 3 .10/ site - packages/ scanpy / get/ get. py”,
   line 394, in _get_obs_rep
   raise Type Error(f” use_raw expected to be bool, was { type (use_raw)}.”)
Type Error: use_raw expected to be bool, was <class ‘ anndata. _core. anndata. AnnData ‘ >.
‘
“““
~~~

`Possible solution info = ““`

`tutorial examples = “““`

~~~
import scanpy as sc
from matplotlib. pyplot import rc_context
sc. set_figure_params (dpi =100, color_map =“ viridis_r “)
sc. settings. verbosity = 0
sc. logging. print_header ()
pbmc = sc. datasets. pbmc68 k_reduced ()
# inspect pbmc contents
pbmc
# rc_context is used for the figure size, in this case 4 x4
with rc_context ({“ figure. figsize “: (4, 4)}):
sc. pl. umap (pbmc, color =“ CD79A “)
color_vars = [
 “ CD79A “,
 “ MS4A1 “,
 “ IGJ”,
 “ CD3D “,
 “ FCER 1 A “,
 “ FCGR 3 A “,
 “ n_counts”,
 “ bulk_labels “,
]
with rc_context ({“ figure. figsize “: (3, 3)}):
 sc. pl. umap (pbmc, color= color_vars, s=50, frameon = False, ncols =4, vmax =“ p99 “)
# compute clusters using the leiden method and store the results with the name ‘ clusters ‘
sc. tl. leiden (
 pbmc,
 key_added =“ clusters”,
 resolution =0.5,
 n_iterations =2,
 flavor =“ igraph “,
 directed = False,
)
with rc_context ({“ figure. figsize “: (5, 5)}):
 sc. pl. umap (
   pbmc,
   color =“ clusters”,
   add_outline =True,
   legend_loc =“ on data “,
   legend_fontsize =12,
   legend_fontoutline =2,
   frameon = False,
   title =“ clustering of cells”,
   palette =“ Set1 “,
)
marker_genes_dict = {
 “B-cell “: [“ CD79A “, “ MS4A1 “],
 “ Dendritic “: [“ FCER 1 A “, “ CST3 “],
 “ Monocytes “: [“ FCGR 3 A “],
 “ NK “: [“ GNLY “, “ NKG7 “],
 “ Other “: [“ IGLL1 “],
 “ Plasma “: [“ IGJ”],
 “T-cell “: [“ CD3D “],
}
sc. pl. dotplot(pbmc, marker_genes_dict, “ clusters”, dendrogram = True)
# create a dictionary to map cluster to annotation label
cluster2 annotation = {
 “0”: “ Monocytes”,
 “1”: “ NK”,
 “2”: “T-cell”,
 “3”: “ Dendritic”,
 “4”: “ Dendritic”,
 “5”: “ Plasma “,
 “6”: “B-cell”,
 “7”: “ Dendritic”,
 “8”: “ Other”,
}
# add a new ‘. obs ‘ column called ‘ cell type ‘ by mapping clusters to annotation using pandas ‘
 map ‘ function
pbmc. obs [“ cell type “] = pbmc. obs [“ clusters “]. map (cluster2 annotation). astype (“ category “)
sc. pl. dotplot(pbmc, marker_genes_dict, “ cell type “, dendrogram = True)
sc. pl. umap (
 pbmc,
 color =“ cell type “,
 legend_loc =“ on data “,
 frameon = False,
 legend_fontsize =10,
 legend_fontoutline =2,
)
with rc_context ({“ figure. figsize “: (4.5, 3)}):
 sc. pl. violin (pbmc, [“ CD79A “, “ MS4A1 “], groupby =“ clusters “)
with rc_context ({“ figure. figsize “: (4.5, 3)}):
 sc. pl. violin (
   pbmc,
   [“ n_genes”, “ percent_mito “],
   groupby =“ clusters”,
   stripplot= False, # remove the internal dots
   inner =“ box”, # adds a boxplot inside violins
)
ax = sc. pl. stacked_violin (
 pbmc, marker_genes_dict, groupby =“ clusters”, swap_axes= False, dendrogram = True
)
sc. pl. matrixplot (
 pbmc,
 marker_genes_dict,
 “ clusters”,
 dendrogram =True,
 cmap =“ Blues”,
 standard_scale =“ var”,
 colorbar_title =“ column scaled \ nexpression “,
)
# scale and store results in layer
pbmc. layers [“ scaled “] = sc. pp. scale (pbmc, copy = True). X
sc. pl. matrixplot (
 pbmc,
 marker_genes_dict,
 “ clusters”,
 dendrogram =True,
 colorbar_title =“ mean z-score “,
 layer =“ scaled “,
 vmin =-2,
 vmax =2,
 cmap =“ RdBu_r”,
)
import matplotlib. pyplot as plt
fig, (ax1, ax2, ax3) = plt. subplots (1, 3, figsize =(20, 4), gridspec_kw ={“ wspace “: 0.9})
ax1 _dict = sc. pl. dotplot(
 pbmc, marker_genes_dict, groupby =“ bulk_labels “, ax=ax1, show = False
)
ax2 _dict = sc. pl. stacked_violin (
 pbmc, marker_genes_dict, groupby =“ bulk_labels “, ax=ax2, show = False
)
ax3 _dict = sc. pl. matrixplot(
 pbmc, marker_genes_dict, groupby =“ bulk_labels “, ax=ax3, show = False, cmap =“ viridis”
)
ax = sc. pl. heatmap (
 pbmc, marker_genes_dict, groupby =“ clusters”, cmap =“ viridis”, dendrogram = True
)
ax = sc. pl. heatmap (
 pbmc,
 marker_genes_dict,
 groupby =“ clusters”,
 layer =“ scaled “,
 vmin =-2,
 vmax =2,
 cmap =“ RdBu_r”,
 dendrogram =True,
 swap_axes=True,
 figsize =(11, 4),
)
ax = sc. pl. tracksplot(pbmc, marker_genes_dict, groupby =“ clusters”, dendrogram = True)
sc. tl. rank_genes_groups (pbmc, groupby =“ clusters”, method =“ wilcoxon “)
sc. pl. rank_genes_groups_dotplot (pbmc, n_genes =4)
sc. pl. rank_genes_groups_dotplot (
 pbmc,
 n_genes =4,
 values_to_plot =“ logfoldchanges “,
 min_logfoldchange =3,
 vmax =7,
 vmin =-7,
 cmap =“ bwr”,
)
sc. pl. rank_genes_groups_dotplot (
 pbmc,
 n_genes =30,
 values_to_plot =“ logfoldchanges “,
 min_logfoldchange =4,
 vmax =7,
 vmin =-7,
 cmap =“ bwr”,
 groups =[“3”, “7”],
)
sc. pl. rank_genes_groups_matrixplot (
 pbmc, n_genes =3, use_raw = False, vmin =-3, vmax =3, cmap =“ bwr”, layer =“ scaled “
)
sc. pl. rank_genes_groups_stacked_violin (pbmc, n_genes =3, cmap =“ viridis_r “)
sc. pl. rank_genes_groups_heatmap (
 pbmc,
 n_genes =3,
 use_raw = False,
 swap_axes=True,
 vmin =-3,
 vmax =3,
 cmap =“ bwr”,
 layer =“ scaled “,
 figsize =(10, 7),
 show = False,
)
sc. pl. rank_genes_groups_heatmap (
 pbmc,
 n_genes =10,
 use_raw = False,
 swap_axes=True,
 show_gene_labels = False,
 vmin =-3,
 vmax =3,
 cmap =“ bwr”,
)
sc. pl. rank_genes_groups_tracksplot (pbmc, n_genes =3)
with rc_context ({“ figure. figsize “: (9, 1.5) }):
 sc. pl. rank_genes_groups_violin (pbmc, n_genes =20, jitter= False)
# compute hierarchical clustering using PCs (several distance metrics and linkage methods are
 available).
sc. tl. dendrogram (pbmc, “ bulk_labels “)
ax = sc. pl. dendrogram (pbmc, “ bulk_labels “)
ax = sc. pl. correlation_matrix (pbmc, “ bulk_labels “, figsize =(5, 3.5))
“““
~~~

API docstring = “““

~~~
def scanpy. pl. dotplot(adata : AnnData, var_names: Union [ str, Sequence [ str], Mapping [ str, Union [
 str, Sequence [ str ]]]], groupby : Union [ str, Sequence [ str]], use_raw : groupby :, Optional[ bool] = None,
 log: bool = False, num_categories : int = 7, expression_cutoff : float = 0.0,
 mean_only_expressed : bool = False, cmap : str = Reds, dot_max : Optional[ float] = None,
 dot_min : Optional[ float] = None, standard_scale : Optional[ Literal[ var, group ]] = None,
 smallest_dot : Optional[ float] = 0.0, title : Optional[ str] = None, colorbar_title : Optional
 [ str] = Mean expression
in group, size_title : Optional[ str] = Fraction of cells
in group (%), figsize : Optional[ tuple [ float, float ]] = None, dendrogram : Union [ bool, str] =
 False, gene_symbols : Optional[ str] = None, var_group_positions : Optional[ Sequence [ tuple [
 int, int ]]] = None, var_group_labels : Optional[ Sequence [ str ]] = None, var_group_rotation :
 Optional[ float] = None, layer: Optional[ str] = None, swap_axes: Optional[ bool] = False,
 dot_color_df : Optional[ Data Frame ] = None, show : Optional[ bool] = None, save : Optional[
 Union [ str, bool ]] = None, ax: Optional[ _AxesSubplot ] = None, return_fig : Optional[ bool] =
 False, vmin : Optional[ float] = None, vmax : Optional[ float] = None, vcenter: Optional[ float
 ] = None, norm : Optional[ Normalize ] = None):
 “““ Makes a * dot plot* of the expression values of ‘ var_names ‘.
 For each var_name and each ‘ groupby ‘ category a dot is plotted.
 Each dot represents two values: mean expression within each category
 (visualized by color) and fraction of cells expressing the ‘ var_name ‘ in the
 category (visualized by the size of the dot). If ‘ groupby ‘ is not given,
 the dotplot assumes that all data belongs to a single category.
 .. note ::
   A gene is considered expressed if the expression value in the ‘ adata ‘ (or
   ‘adata. raw ‘) is above the specified threshold which is zero by default.
 An example of dotplot usage is to visualize, for multiple marker genes,
 the mean value and the percentage of cells expressing the gene
 across multiple clusters.
 This function provides a convenient interface to the : class :’∼ scanpy. pl. DotPlot ‘
 class. If you need more flexibility, you should use : class :’∼ scanpy. pl. DotPlot ‘ directly.
 Parameters
 ----------
 adata
   Annotated data matrix.
 var_names
   ‘ var_names ‘ should be a valid subset of ‘ adata. var_names ‘.
   If ‘ var_names ‘ is a mapping, then the key is used as label
   to group the values (see ‘ var_group_labels ‘). The mapping values
   should be sequences of valid ‘ adata. var_names ‘. In this
   case either coloring or ‘ brackets ‘ are used for the grouping
   of var_names depending on the plot. When ‘ var_names ‘ is a mapping,
   then the ‘ var_group_labels ‘ and ‘ var_group_positions ‘ are set.
 groupby
   The key of the observation grouping to consider.
 use_raw
   Use ‘raw ‘ attribute of ‘ adata ‘ if present.
 log
   Plot on logarithmic axis.
 num_categories
   Only used if groupby observation is not categorical. This value
   determines the number of groups into which the groupby observation
   should be subdivided.
 categories_order
   Order in which to show the categories. Note : add_dendrogram or add_totals
   can change the categories order.
 figsize
   Figure size when ‘ multi_panel =True ‘.
   Otherwise the ‘ rcParam [‘ figure. figsize ]’ value is used.
   Format is (width, height)
 dendrogram
   If True or a valid dendrogram key, a dendrogram based on the hierarchical
   clustering between the ‘ groupby ‘ categories is added.
   The dendrogram information is computed using : func:’ scanpy. tl. dendrogram ‘.
   If ‘ tl. dendrogram ‘ has not been called previously the function is called
   with default parameters.
 gene_symbols
   Column name in ‘. var ‘ Data Frame that stores gene symbols.
   By default ‘ var_names ‘ refer to the index column of the ‘. var ‘ Data Frame.
   Setting this option allows alternative names to be used.
 var_group_positions
   Use this parameter to highlight groups of ‘ var_names ‘.
   This will draw a ‘ bracket ‘ or a color block between the given start and end
   positions. If the parameter ‘ var_group_labels ‘ is set, the corresponding
   labels are added on top / left. E. g. ‘ var_group_positions =[(4, 10)]’
   will add a bracket between the fourth ‘ var_name ‘ and the tenth ‘ var_name ‘.
   By giving more positions, more brackets/ color blocks are drawn.
 var_group_labels
   Labels for each of the ‘ var_group_positions ‘ that want to be highlighted.
 var_group_rotation
   Label rotation degrees.
   By default, labels larger than 4 characters are rotated 90 degrees.
 layer
   Name of the Ann Data object layer that wants to be plotted. By default adata. raw. X is
 plotted.
   If ‘ use_raw = False ‘ is set, then ‘ adata .X’ is plotted. If ‘ layer ‘ is set to a valid
 layer name,
   then the layer is plotted. ‘ layer ‘ takes precedence over ‘ use_raw ‘.
 title
   Title for the figure
 colorbar_title
   Title for the color bar. New line character (\ n) can be used.
 cmap
   String denoting matplotlib color map.
 standard_scale
   Whether or not to standardize the given dimension between 0 and 1, meaning for
   each variable or group, subtract the minimum and divide each by its maximum.
 swap_axes
   By default, the x axis contains ‘ var_names ‘ (e. g. genes) and the y axis
   the ‘ groupby ‘ categories. By setting ‘ swap_axes ‘ then x are the ‘
   groupby ‘ categories and y the ‘ var_names ‘.
 return_fig
   Returns : class:’ DotPlot ‘ object. Useful for fine - tuning
   the plot. Takes precedence over ‘ show = False ‘.
 size_title
   Title for the size legend. New line character (\ n) can be used.
 expression_cutoff
   Expression cutoff that is used for binarizing the gene expression and
   determining the fraction of cells expressing given genes. A gene is
   expressed only if the expression value is greater than this threshold.
 mean_only_expressed
   If True, gene expression is averaged only over the cells
   expressing the given genes.
 dot_max
   If none, the maximum dot size is set to the maximum fraction value found
   (e. g. 0.6). If given, the value should be a number between 0 and 1.
   All fractions larger than dot_max are clipped to this value.
 dot_min
   If none, the minimum dot size is set to 0. If given,
   the value should be a number between 0 and 1.
   All fractions smaller than dot_min are clipped to this value.
 smallest_dot
   If none, the smallest dot has size 0.
   All expression levels with ‘ dot_min ‘ are plotted with this size.
 show
   Show the plot, do not return axis.
 save
   If ‘ True ‘ or a ‘str ‘, save the figure.
   A string is appended to the default filename.
   Infer the filetype if ending on {‘‘. pdf ‘‘, ‘‘. png ‘‘, ‘‘. svg ‘‘}.
 ax
   A matplotlib axes object. Only works if plotting a single component.
 vmin
   The value representing the lower limit of the color scale. Values smaller than vmin
 are plotted
   with the same color as vmin.
 vmax
   The value representing the upper limit of the color scale. Values larger than vmax
 are plotted
   with the same color as vmax.
 vcenter
   The value representing the center of the color scale. Useful for diverging colormaps.
 norm
   Custom color normalization object from matplotlib. See
   ‘ https://matplotlib.org/stable/tutorials/colors/colormapnorms.html ‘ for details.
 kwds
   Are passed to : func:’ matplotlib. pyplot. scatter ‘.
 Returns
 -------
 If ‘ return_fig ‘ is ‘ True ‘, returns a : class :’∼ scanpy. pl. DotPlot ‘ object,
 else if ‘ show ‘ is false, return axes dict
 See also
 --------
 : class :’∼ scanpy. pl. DotPlot ‘: The DotPlot class can be used to to control
   several visual parameters not available in this function.
 : func:’∼ scanpy. pl. rank_genes_groups_dotplot ‘: to plot marker genes
   identified using the : func:’∼ scanpy. tl. rank_genes_groups ‘ function.
 Examples
 --------
 Create a dot plot using the given markers and the PBMC example dataset grouped by
 the category ‘ bulk_labels ‘.
 .. plot ::
   : context: close - figs
   import scanpy as sc
   adata = sc. datasets. pbmc68 k_reduced ()
   markers = [‘ C1QA ‘, ‘ PSAP ‘, ‘ CD79A ‘, ‘ CD79B ‘, ‘ CST3 ‘, ‘LYZ ‘]
   sc. pl. dotplot(adata, markers, groupby =‘ bulk_labels ‘, dendrogram = True)
 Using var_names as dict:
 .. plot ::
   : context: close - figs
   markers = {‘T-cell ‘: ‘ CD3D ‘, ‘B-cell ‘: ‘ CD79A ‘, ‘ myeloid ‘: ‘ CST3 ‘}
   sc. pl. dotplot(adata, markers, groupby =‘ bulk_labels ‘, dendrogram = True)
 Get DotPlot object for fine tuning
 .. plot ::
   : context: close - figs
   dp = sc. pl. dotplot(adata, markers, ‘ bulk_labels ‘, return_fig = True)
   dp. add_totals (). style (dot_edge_color =‘ black ‘, dot_edge_lw =0.5). show ()
 The axes used can be obtained using the get_axes () method
 .. code - block :: python
   axes_dict = dp. get_axes ()
   print(axes_dict)”““.
“““
~~~

\# **Example Output**:

~~~
{
 “ analysis “: “ The initial error occurred because the ‘ use_raw ‘ parameter was incorrectly set
   to an Ann Data object, whereas it should be a boolean value indicating whether to use the
   raw attribute of the Ann Data object. To fix this, we need to set ‘ use_raw ‘ to either True
   or False based on whether we want to utilize the raw data or not.”,
 “ code “: “ result_plot = dotplot(result_1, var_names= list(result_1. var_names [0 :10 ]), groupby =‘
   louvain ‘, use_raw = False, show = True)”
}
~~~

